# PAX6 LIMITS THE COMPETENCE OF DEVELOPING CEREBRAL CORTICAL CELLS

**DOI:** 10.1101/2022.02.03.478927

**Authors:** Martine Manuel, Kai Boon Tan, Zrinko Kozic, Michael Molinek, Tiago Sena Marcos, Maizatul Fazilah Abd Razak, Dániel Dobolyi, Ross Dobie, Beth E. P. Henderson, Neil C. Henderson, Wai Kit Chan, Michael I. Daw, John O. Mason, David J. Price

**Affiliations:** Simons Initiative for the Developing Brain, Patrick Wild Centre, University of Edinburgh, Hugh Robson Building, George Square, Edinburgh EH8 9XD, UK

**Author notes:** These authors contributed equally (Martine Manuel, Kai Boon Tan). Current address: Wolfson Institute for Biomedical Research and Department of Neuroscience, Physiology and Pharmacology, University College London, London WC1E 6BT, UK. Centre for Inflammation Research, University of Edinburgh, Queen’s Medical Research Institute, Edinburgh EH16 4TJ, UK. Co-affiliation: Zhejiang University-University of Edinburgh Institute, Zhejiang University, 718 East Haizhou Rd., Haining, Zhejiang 314400, P.R. China.

## Abstract

The development of stable specialized cell types in multicellular organisms relies on mechanisms controlling inductive intercellular signals and the competence of cells to respond to such signals. In developing cerebral cortex, progenitors generate only glutamatergic excitatory neurons despite being exposed to signals with the potential to initiate the production of other neuronal types, suggesting that their competence is limited. Here we tested the hypothesis that this limitation is due to their expression of transcription factor Pax6. We used bulk and single cell RNAseq to show that conditional cortex-specific Pax6 deletion from the onset of cortical neurogenesis allowed some progenitors to generate abnormal lineages resembling those normally found outside the cortex. Analysis of selected gene expression showed that the changes occurred in specific spatiotemporal patterns. We then compared the responses of control and Pax6-deleted cortical cells to in vivo and in vitro manipulations of extracellular signals. We found that Pax6 loss increased cortical progenitors’ competence to generate inappropriate lineages in response to extracellular factors normally present in developing cortex, including the morphogens Shh and Bmp4. Regional variation in the levels of these factors could explain spatiotemporal patterns of fate change following Pax6 deletion in vivo. We propose that Pax6’s main role in developing cortical cells is to minimize the risk of their development being derailed by the potential side-effects of morphogens engaged contemporaneously in other essential functions.

## INTRODUCTION

Gene regulatory networks (GRNs) modulated by intercellular signals control the generation of the specialized cell types that compose multicellular organisms (Britten and Davidson, 1969; Davidson, 2010). These control mechanisms affect the developmental trajectories of cells in a variety of ways to guide the production of particular cell types and prevent the emergence of alternatives.

Transcription factors whose levels vary among developing cells in precise, reproducible spatiotemporal patterns are essential components of GRNs. In some cases, their regional activation in response to inductive signals drives the production of region-specific cell types, but there are many other ways in which they can operate. For example, they can determine whether, and if so how, cells respond when confronted by inductive signals, i.e. their competence (Waddington, 1932 & 1934). Restricting the competence of cells as they develop is likely to maximize the probability of them following reproducibly their stereotypical developmental trajectories, e.g. by mitigating the effects of biochemical noise in the signals they encounter or in the intracellular pathways processing those signals (Perkins and Swain, 2009) and by preventing them responding in inappropriate ways to signalling molecules surrounding them.

The cerebral cortex is a complex amalgamation of two major neuronal cell classes generated by developmental cell lineages expressing different sets of transcription factors (Anderson et al., 1997, 2001 & 2002; Lim et al., 2018). One cell class uses the excitatory neurotransmitter glutamate to propagate neuronal activity through cortical circuits and is produced by progenitors located in the developing cerebral cortex itself. It develops from cell lineages that express transcription factors including Pax6, Neurog2 and Eomes. The other cell type uses the inhibitory neurotransmitter gamma aminobutyric acid (GABA) to refine and elaborate patterns of cortical neuronal activity and is produced by progenitors located subcortically. It develops from cell lineages that express substantially different sets of transcription factors. Pax6 is one of the first transcription factors to be expressed differentially between the progenitors of excitatory and inhibitory cortical neurons (Stoykova and Gruss, 1994; Mastick et al., 1997; Caric et al., 1997), making it a good candidate to be involved in regulating the likelihood of cortical progenitors adopting an excitatory neuronal fate.

The *Pax6* gene emerged 500-700 million years ago and has been conserved through all triproblastic animal lineages, where it is involved in many neural and non-neural processes (Callaerts et al., 1997; Simpson and Price, 2002). Its expression in the developing brain of extant vertebrates and invertebrates indicates that it acquired important functions very early in this organ’s evolution. In mammalian embryos, it is activated prior to neural tube closure in the anterior neuroectoderm where brain forms (Shimamura and Rubenstein, 1997). Its importance for the production of cortical excitatory neurons is demonstrated by the phenotypes of constitutively mutant mouse embryos unable to make functional Pax6. These embryos show reduced cortical expression of genes involved in excitatory neuron production and increased cortical expression of genes involved in the development of subcortically-derived cell types including inhibitory interneurons (Toresson et al., 2000; Yun et al., 2001; Heins et al., 2002; Scardigli et al., 2003; Schuurmans et al., 2004; Kroll and O’Leary, 2005; Holm et al., 2007; Quinn et al., 2009; Guo et al., 2019). We set out to discover what *Pax6* does in cortical progenitors to help govern their normal production of excitatory neurons.

We began by examining the effects of inducing cortex-specific *Pax6* loss-of-function in cortical progenitors using population and single cell transcriptomics followed by expression analysis of selected genes in tissue sections. The response was dichotomous: many *Pax6*-null progenitors continued to generate excitatory neurons that made cortical layers relatively normally, while others adopted abnormal developmental trajectories, the nature of which varied with age and cortical location. Subsequent in vivo and in vitro experiments revealed that Pax6 blocks the deviant trajectories by reducing the ability of cortical cells to react abnormally to substances normally present - and carrying out other essential functions - around them. We propose that the main function of Pax6 in cortical development is to imbue the process with stability and reproducibility by protecting it from potentially destabilizing signals in the cortical environment.

## RESULTS

### Removal of Pax6 from the progenitors of cortical neurons

Most cortical excitatory neurons are generated between embryonic day 12.5 (E12.5) and E16.5 in mice (Angevine and Sidman, 1961; Smart and Smart, 1982; Caviness, 1982; del Rio and Soriano, 1989; Gillies and Price, 1993). They are derived from cortical radial glial progenitors (RGPs), some directly and others indirectly via the initial production of transit-amplifying intermediate progenitors (IPs) (Noctor et al., 2004; Vasistha et al., 2015; Villalba et al., 2021). All RGPs express Pax6 (Caric et al., 1997). We used the *Emx1-Cre^ERT2^* allele (Kessaris et al., 2006) to make tamoxifen-induced cortex-specific homozygous *Pax6* conditional knock outs (*Pax6* cKOs) (Fig. 1A: Fig. S1A). Heterozygous littermates with deletion in just one *Pax6* allele served as controls; previous work on heterozygotes detected no abnormalities in cortical levels and patterns of Pax6 protein expression or cortical morphogenesis, almost certainly because known feedback mechanisms caused compensatory increases in Pax6 production from the normal allele (Manuel et al., 2007; Mi et al., 2013; Quintana et al., 2018). When we gave tamoxifen at E9.5 (tamoxifen^E9.5^), levels of normal *Pax6* mRNA in *Pax6* cKOs fell to <50% of control by E11.5, to ∼10% of control by E12.5 and to almost zero by E13.5 (Fig. S1B) and levels of Pax6 protein fell to ∼5% of control by E12.5 (Fig. S1C,D). By E12.5, Pax6 was undetectable by immunohistochemistry in almost all RGPs (except those in a narrow ventral pallial domain where *Emx1* is not expressed) (Fig. S1E,F) while a Cre reporter, *RCE^EGFP^* (Fig. S1A; Miyoshi et al., 2010), was active in most cortical cells (Fig. S1E). Thus, tamoxifen^E9.5^ ensured that the vast majority of cortical neurons was generated, directly or indirectly, from RGPs that had lost Pax6 protein.

**Figure 1:**
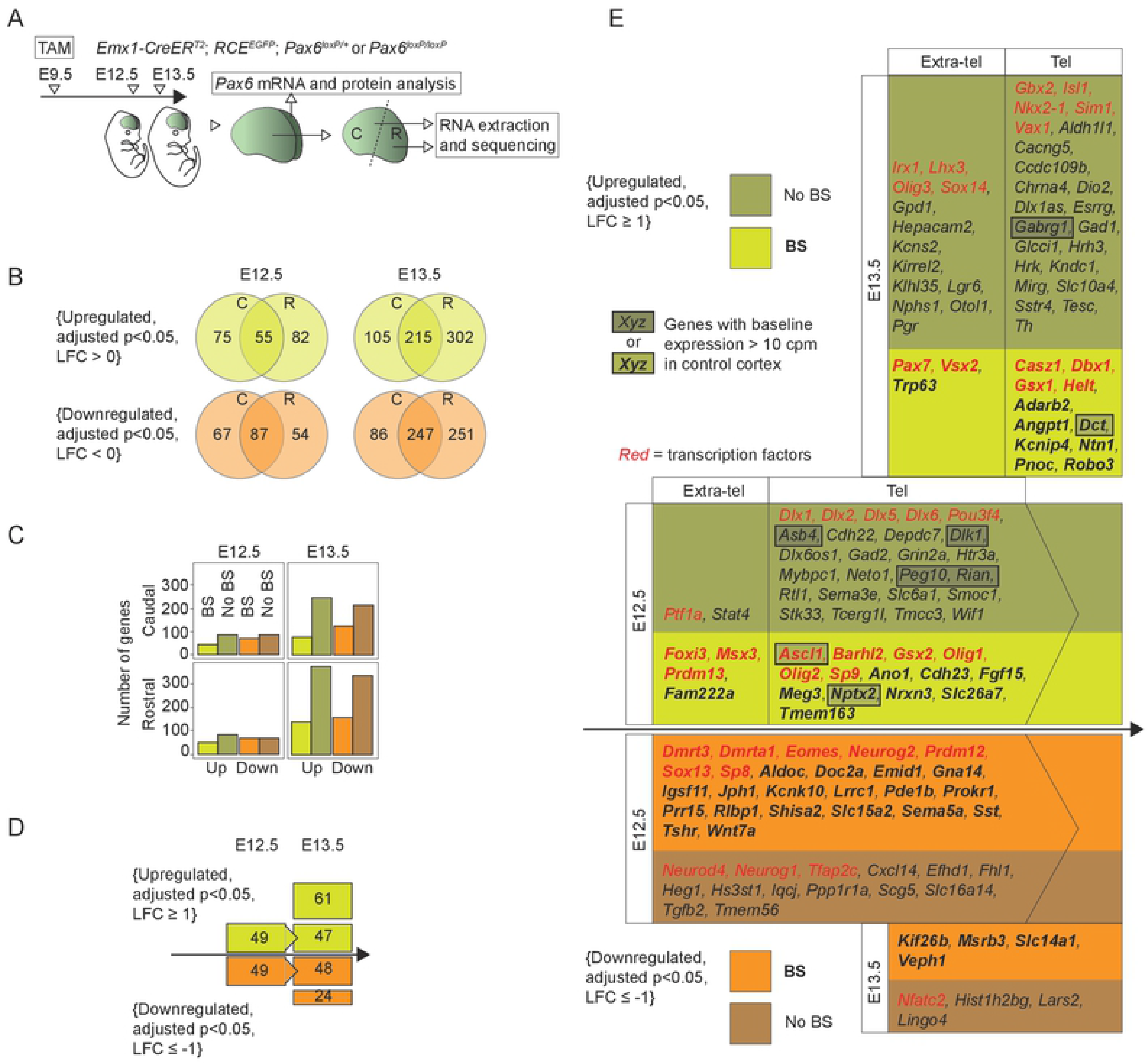
Pax6 loss altered gene expression in cortical cells. **(A)** The experimental procedure: tamoxifen (TAM) was administered at E9.5; one hemisphere from each E12.5 or E13.5 embryo was used to assess Pax6 mRNA and protein levels; the rostral and caudal halves of the other hemisphere were processed for RNAseq. **(B)** Numbers of genes with significantly upregulated or downregulated expression levels in caudal (C) and rostral (R) *Pax6* cKO cortex (LFC: log fold change). **(C)** Numbers of significantly upregulated or downregulated genes with a nearby Pax6 binding site (BS). **(D)** Changes with age in the numbers of significantly upregulated or downregulated genes with LFCs ≥ 1 or ≤ -1 in at least one of the four combinations of age and region. **(E)** Lists of upregulated and downregulated genes with LFCs ≥ 1 or ≤ -1 at E12.5 and/ or E13.5 in at least one of the four combinations of age and region. Genes were arranged according to when they first showed upregulation or downregulation (adjusted p<0.05, irrespective of LFC magnitude) and whether they had a nearby Pax6 binding site (BS). Upregulated genes were separated into those normally expressed by telencephalic cells (Tel) and those normally expressed only outside the telencephalon (Extra-tel). Upregulated genes shown in boxes were clearly expressed in control cortex, i.e. their counts per million (cpm) were > 10. Genes encoding transcription factors were listed using red font.

### Pax6 loss caused ectopic gene expression in cortical cells

We first used bulk RNAseq to study the effects of tamoxifen^E9.5^-induced *Pax6* cKO in rostral and caudal cortex at E12.5 and E13.5 (Fig. 1A). Raw data is available at the European Nucleotide Archive accession numbers PRJEB5857 and PRJEB6774. We used 4 biological replicates for each location, age and genotype; principal component analysis (PCA) on all datasets taken together showed high-level clustering by age and location (Fig. S1G).

The number of genes with significantly altered expression levels (adjusted p<0.05) in *Pax6* cKO cortex was ∼3-fold higher at E13.5 than at E12.5 (Fig. 1B; Table S1). At each age, the numbers of upregulated and downregulated genes were similar. We identified regulated genes with nearby Pax6 binding sites using published ChIPseq data from E12.5 forebrain (Sun et al., 2015). We followed their assignation of peaks to the gene with the nearest transcription start site (TSS), provided the peak lay within the genomic interval between 50kb upstream of the TSS and 50kb downstream of the transcription end site. The proportion of regulated genes with a nearby binding site was higher at E12.5 than E13.5 (Fig. 1C), suggesting an accumulation of indirect gene expression changes with age.

We then focussed on the genes that underwent the largest changes in expression levels. We included all functionally annotated genes (http://www.ensembl.org/index.html) with a log_2_ fold change (LFC) in expression ≥ 1 (for upregulated genes) or ≤ -1 (for downregulated genes) in at least one of the four combinations of age and region. This produced a subset of 183 genes: 98 genes were affected at E12.5, 95 remained so and a further 85 were added at E13.5 (Fig. 1D). To gain an initial impression of biological processes strongly associated with these genes, we passed them through the Database for Annotation, Visualization and Integrated Discovery v6.8 (DAVID v6.8; Huang et al., 2009a,b) to obtain sets of significantly enriched gene ontology (GO) terms (Table S2). Some of the GO terms obtained using the upregulated gene set described the development of cell types normally generated within the telencephalon but outside the cortex in subpallium (where cerebral cortical GABAergic interneurons and, at these ages, oligodendrocytes are made; Kessaris et al., 2006; Miyoshi et al., 2007; Lim et al., 2018). Others described the development of non-telencephalic cell types (spinal cord, inner ear, skeletal system and neural crest).

We then sorted this set of 183 genes according to: (i) whether the first sign of their upregulation or downregulation (which was defined as an adjusted p<0.05 irrespective of the magnitude of the LFC) was at E12.5 or E13.5 and (ii) whether they had a nearby Pax6 binding site (as defined above; Sun et al., 2015) (Fig. 1E). We excluded downregulated genes whose expression levels were very low in our E12.5 and E13.5 control RNAseq data and in sections of normal embryos of similar ages contained in the following gene expression databases: https://gp3.mpg.de/; https://developingmouse.brain-map.org/; http://www.informatics.jax.org/gxd. We included all upregulated genes for which there was sufficient documented evidence to allow us to reach valid conclusions on their normal sites of expression at these ages (Table S3 contains citations). We separated these upregulated genes into those normally expressed by telencephalic cells (“Tel” in Fig. 1E) and those normally expressed only outside the telencephalon (“Extra-tel” in Fig. 1E). Of the genes normally expressed in telencephalon, most had little or no expression in our control RNAseq data from E12.5 and E13.5 cortex (mean counts per million ≤ 10).

We concluded that a major consequence of *Pax6* deletion from cortical cells was to allow their ectopic activation of genes normally expressed extra-cortically, either by surrounding telencephalic cells or by cells normally located outside the telencephalon.

Our present study provides no new evidence on whether the genes with altered cortical expression following conditional Pax6 deletion are normally directly regulated by Pax6 binding to their enhancers or promoters (for previous data on this, in addition to those from Sun et al. 2015 used above, see Scardigli et al. 2003, Sansom et al. 2009 and Xie et al. 2013). A large proportion of the genes showing altered expression in our present study were transcription factors (Fig. 1E), many of which are known to regulate genes whose expression levels were altered in our *Pax6* cKOs (Corbin et al., 2000; Fode et al., 2000; Toresson et al., 2000; Yun et al., 2001 & 2002; Castro et al., 2006 & 2011; Petryniak et al., 2007; Long et al., 2009; Pinto et al., 2009; Waclaw et al., 2009; Nishida et al., 2010; Seibt et al., 2012; Kikkawa et al., 2013; Kovach et al., 2013; Wang et al., 2013; Le et al., 2017; Sessa et al., 2017; Pla et al., 2017; Guo et al., 2019). Thus, acute conditional cortex-specific Pax6 removal affected, almost certainly both directly and indirectly, the specification of the developmental trajectories of cortical progenitors and their daughters. We next examined the heterogeneity of these effects within the population of cortical progenitors.

### Pax6 loss caused cortical cell lineage progressions to diversify

We used single cell RNAseq (scRNAseq) to explore the lineage progression of cortical cells following Pax6 removal. We used the alleles described above (Fig. S1A) and separated green fluorescent protein (GFP)-expressing cells by fluorescence-activated cell sorting (FACS) from single cell suspensions of E13.5 and E14.5 *Pax6* cKO (tamoxifen^E9.5^) and control rostral cortex before carrying out scRNAseq (Fig. 2A). We focussed on rostral cortex, since it contained ∼85% of changes detected by bulk RNAseq at E13.5 (Fig. 1B). This gave four datasets of: 6,266 cells from E13.5 *Pax6* cKO; 3,744 cells from E13.5 control; 4,259 cells from E14.5 *Pax6* cKO; 4,137cells from E14.5 control. Raw data is available at the European Nucleotide Archive accession numbers PRJEB27937 and PRJEB32740.

**Figure 2:**
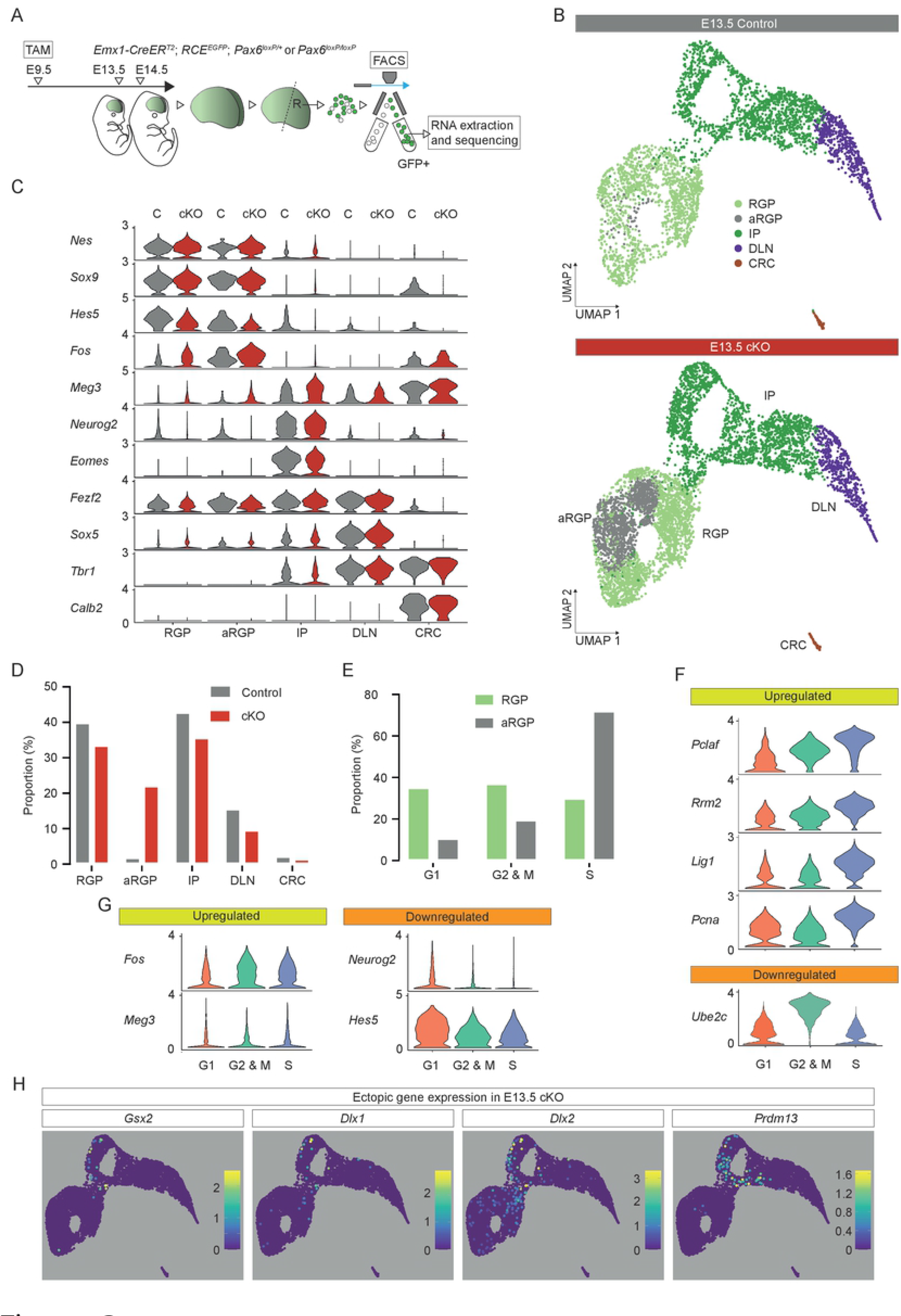
Aberrant cell type and ectopic gene expression in E13.5 *Pax6* cKO cortex. **(A)** The experimental procedure: tamoxifen (TAM) was administered at E9.5; E13.5 and E14.5 rostral (R) cortex was dissociated into single cell suspension; viable GFP+ cells were selected by fluorescence activated cell sorting (FACS) for single cell RNA sequencing (scRNAseq). **(B)** Uniform manifold approximation and projection (UMAP) plot of the scRNAseq data from *Pax6* cKO and control cells at E13.5. Data from the two genotypes were analysed together and then split for visualization. RGP, radial glial progenitor; aRGP, atypical RGP; IP, intermediate progenitor; DLN, deep layer neuron; CRC, Cajal-Retzius cell. **(C)** Violin plots of selected gene expression in each cell type from control and *Pax6* cKO E13.5 cortex. **(D)** Proportions of cells of each type in control and *Pax6* cKO E13.5 cortex. **(E)** The proportions of RGPs and aRGPs in different cell cycle phases in control and *Pax6* cKO E13.5 cortex. **(F,G)** Violin plots of the expression levels of genes that were significantly differentially expressed between RGPs and aRGPs in different cell cycle phases in control and *Pax6* cKO E13.5 cortex. **(H)** UMAP plots showing log_10_ normalized expression of selected genes that were ectopically expressed in E13.5 *Pax6* cKO cortex.

Differential expression analysis (DEA) using scRNAseq data to calculate Pax6-loss-induced LFCs in average gene expression at E13.5 correlated well with LFCs detected by bulk RNAseq in rostral E13.5 cortex (Fig. S1H), cross-validating data obtained from the two approaches.

At E13.5, comparison of *Pax6* cKO and control samples using Uniform Manifold Approximation and Projection (UMAP) dimensionality reduction indicated a high degree of similarity in their transcriptomic landscapes (Fig. 2B). Graph-based clustering combined with analysis of the expression of cell type-selective marker genes (such as *Nes*, *Sox9*, *Hes5*, *Neurog2*, *Eomes*, *Fezf2*, *Sox5, Tbr1* and *Calb2*; Lendahl et al., 1990; Weisenhorn et al, 1994; Bulfone et al., 1995; Hevner et al., 2001; Englund et al., 2005; Lai et al., 2008; Leone et al., 2008; Kovach et al., 2013; Wang et al., 2013; Kaplan et al., 2017) separated cells of both genotypes into recognized major classes: RGPs, IPs, deep layer neurons (DLNs) and Cajal-Retzius cells (CRCs) (Fig. 2B,C; Fig. S2). It also split the RGPs into two clusters, one of which contained very few cells in controls but many in *Pax6* cKOs (Fig. 2B,D). We called these cells atypical RGPs (aRGPs) and explored their distinguishing features further.

The genes whose expression levels were shown by DEA to be the most different between RGPs and aRGPs (LFCs >0.3 or <-0.3) are listed in Table S4; violin plots for two of them, *Fos* (upregulated) and *Hes5* (downregulated), are in Fig. 2C. Almost all GO terms obtained by passing this list through DAVID described processes involved in mitosis (Table S4). The remainder included terms describing cellular responses to extracellular substances and involved immediate early genes encoding AP-1 transcription factors of the Jun and Fos families, whose expression levels are known to be induced by a range of extracellular signals (Sheng and Greenberg, 1990; Radler-Pohl et al., 1993; Karin et al., 1997; Herdegen and Leah, 1998; Raivich and Behrens, 2006; Tuvikene et al., 2016). This suggested the possibility that cells exposed to the highest levels of extracellular signals developed abnormalities, an idea explored further below.

We identified each RGP’s and aRGP’s cell cycle phase by profiling its expression of known cell cycle phase-selective markers (Kowalczyk et al., 2015). We found that a relatively higher proportion of aRGPs than RGPs were in S phase (Fig. 2E). This is compatible with previous findings that Pax6 removal causes a shortening of G1, G2 and M phases (Mi et al., 2013 and 2018). The expression levels of some of the genes whose expression levels differed between aRGPs and RGPs showed large systematic variation with cell cycle phase; examples are shown in Fig. 2F. The aRGPs’ elevated expression of genes such as *Pclaf*, *Rrm2*, *Lig1* and *Pcna*, whose expression levels increased in S phase, and lowered expression of genes such as *Ube2c*, whose expression levels increased in G2 and M phases, probably reflected the relative increase in the proportions of aRGPs in S phase. However, it was hard to explain all differences between aRGPs and RGPs in this way. For example, levels of *Fos* and *Meg3*, whose expression levels were elevated in aRGPs, and *Neurog2* and *Hes5*, whose expression levels were lowered in aRGPs, showed much less variation with cell cycle phase (Fig. 2G).

At E13.5, some RGPs, aRGPs and IPs in *Pax6* cKOs showing ectopic activation of genes such as *Gsx2*, *Dlx1* and *Dlx2* (selected as examples of genes normally expressed outside the cortex but within the telencephalon: “Tel” in Fig. 1E) and *Prdm13* (an example of a gene normally expressed outside the telencephalon: “Extra-tel” in Fig. 1E) (Fig. 2H). This suggested that diversification of gene expression was occurring as cells progressed from the RGP to the IP identity. This was even clearer a day later.

At E14.5, UMAP dimensionality reduction followed by graph-based clustering combined with analysis of the expression of cell type-selective marker genes (including those used at E13.5 with the addition of layer markers such as *Cux2*, *Satb2* and *Tle4*; Yao et al., 1998; Zimmer et al., 2004; Britanova et al., 2008; Alcamo et al., 2008) separated cells of both genotypes into recognized major classes: RGPs, IPs, superficial layer 2/3 neurons (SLN-L2/3) and layer 4 neurons (SLN-L4), deep layer 5 neurons (DLN-L5) and layer 6 neurons (DLN-L6), and Cajal-Retzius cells (CRCs) (Fig. 3A,B; Fig. S3). The proportions of IPs and SLN-L2/3s were reduced in *Pax6* cKOs (Fig. 3C), in line with previous reports (Schuurmans et al., 2004; Quinn et al., 2007; Tuoc et al., 2009; Georgala et al., 2011).

**Figure 3:**
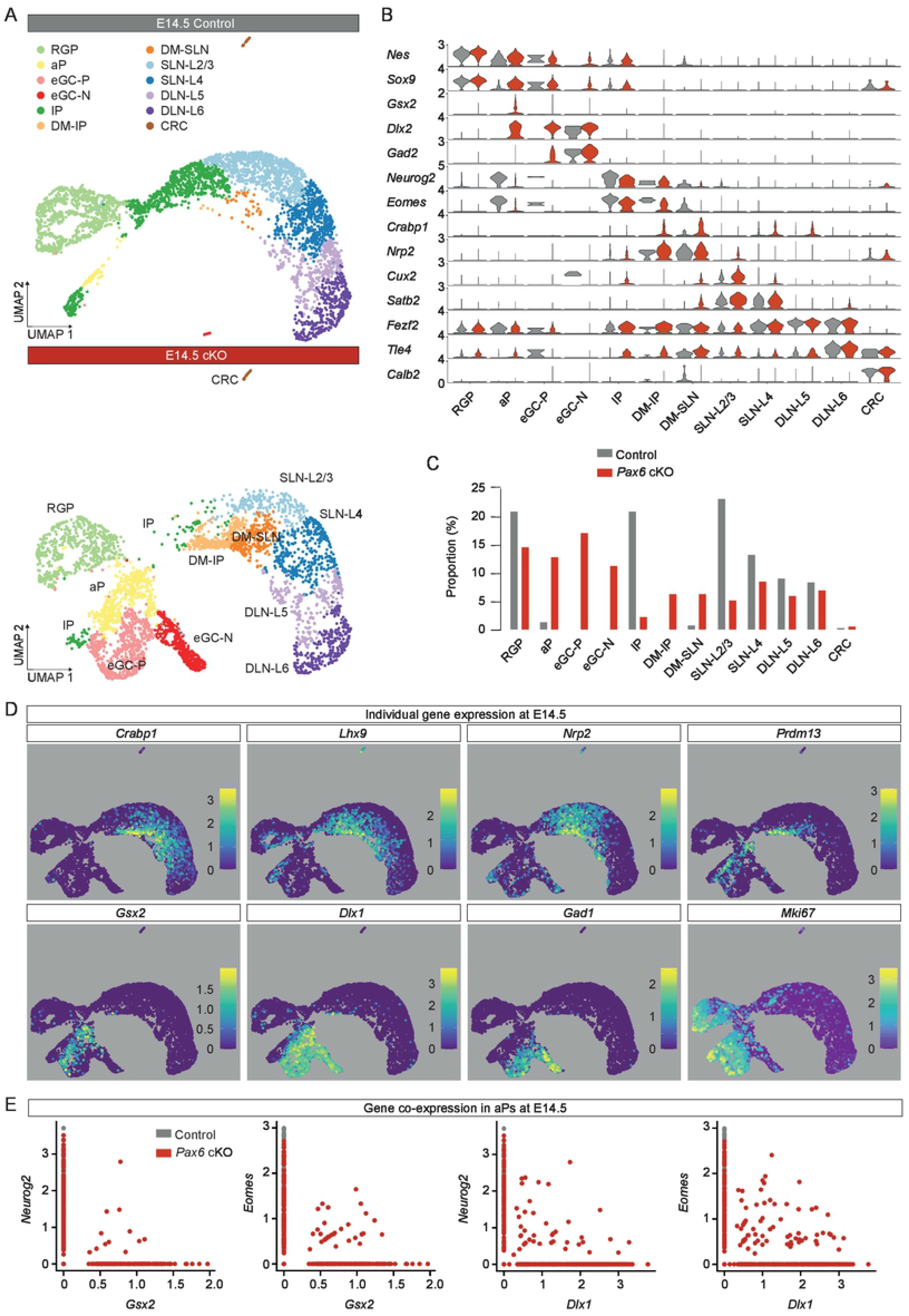
Aberrant cell types and ectopic gene expression in E14.5 *Pax6* cKO cortex. For experimental procedure, see Fig. 2A. **(A)** Uniform manifold approximation and projection (UMAP) plot of the scRNAseq data from *Pax6* cKO and control cells at E14.5. Data from the two genotypes were analysed together and then split for visualization. RGP, radial glial progenitor; aP, atypical progenitor; eGC-P and eCG-N, proliferating or non-proliferating ectopic GABAergic cells; IP, intermediate progenitor; DM-IP and DM-SLN, intermediate progenitor or superficial layer neuron in dorsomedial cortex; SLN-L2/3 and SLN-L4, layer 2/3 or layer 4 superficial layer neurons; DLN-L5 and DLN-L6, layer 5 or layer 6 deep layer neurons; CRC, Cajal-Retzius cell. **(B)** Violin plots of selected gene expression in each cell type from control and *Pax6* cKO E14.5 cortex. **(C)** Proportions of cells of each type in control and *Pax6* cKO E14.5 cortex. **(D)** UMAP plots showing log_10_ normalized expression of cell type-selective marker genes at E14.5. **(E)** Graphs showing co-expression of markers of normal cortical cells (*Neurog2* and *Eomes*) and normal subcortical cells (*Gsx2* and *Dlx1*) in some aPs in E14.5 *Pax6* cKOs.

Five additional clusters populated entirely, or very largely, by *Pax6* cKO cells had emerged. Two of them mapped between IPs and SLNs (Fig. 3A and Fig. S3) and their cells expressed high levels of genes such as *Crabp1*, *Lhx9* and *Nrp2* (Fig. 3D), characteristic of cortical cells located relatively dorsomedially (Bulchand et al., 2003; Le et al., 2007; Lin et al., 2017). We designated cells in these two clusters dorsomedial (DM); the cells in one (DM-IPs) showed greater similarity to IPs than to SLNs (e.g. in terms of *Eomes* and *Neurog2* expression: Fig. 3B and Fig. S3) while cells in the other (DM-SLNs) were more similar to SLNs than to IPs (e.g. in terms of *Cux2* and *Satb2* expression: Fig. 3B and Fig. S3). Many DM cells expressed relatively high levels of *Prdm13* (Fig. 3D; Fig. S3). Another two clusters contained cells expressing genes such as *Gsx2*, *Dlx and Gad* family members, which are normally associated with the development of GABAergic interneurons (Fig. 3B,D and Fig. S3) (Flames et al., 2007; Long et al., 2009; Pan, 2012; Wei et al., 2019). We found that these cells mapped with GABAergic interneurons and the ganglionic eminence (GE) progenitors that generated them when we integrated our E14.5 *Pax6* cKO scRNAseq dataset with data from normal E13.5 and E14.5 ventral telencephalon (Mayer et al., 2018; Fig. S4). We called them ectopic GABAergic cells (eGCs).

A major difference between the two eGC clusters was that one (which we named eGC-P) showed strong expression of markers of proliferating cells (e.g. *Mki67*) while cells in the other (which we named eGC-N) did not (Fig. 3D and Fig. S3). The fifth cluster contained proliferating cells (e.g. *Mki67*-expressing) that we called atypical progenitors (aPs). They were much more common in *Pax6* cKOs than in controls (Fig. 3C). Their gene expression profiles suggested that they were intermediate between other types of cell in both controls and *Pax6* cKOs (Fig. 3B,D and Fig. S3). For example, they were RGP-like in expressing *Nes* and *Sox9* (albeit at lower levels in both genotypes) and IP-like in expressing *Eomes* (at lower levels in *Pax6* cKO cells). In *Pax6* cKOs, they were eGC-like in expressing *Gsx2*, *Dlx* and *Gad* family members. Co-expression analysis revealed that small proportions of aPs in *Pax6* cKOs co-expressed a marker of cells undergoing normal cortical neurogenesis (*Neurog2* and *Eomes*) and a marker of eGCs (e.g. *Gsx2* and *Dlx1*) (Fig. 3E).

Cells in all clusters expressed the telencephalic marker, *Foxg1* (Fig. S3). This indicated that cells undergoing ectopic activation of genes normally expressed in non-telencephalic tissue (e.g. *Prdm13*) did not lose entirely their telencephalic identity.

We next used RNA Velocity (La Manno et al., 2018; Bergen et al., 2020) to explore the direction and speed of movement of individual *Pax6* cKO cortical cells along their predicted developmental trajectories, with particular focus on aPs (Fig. 4). For all control aPs and some *Pax6* cKO aPs, velocities were directed towards IPs (Fig. 4A,B). Whereas some of these cells in *Pax6* cKO cortex expressed markers of normal cortical neurogenesis (e.g. *Neurog2* and *Eomes*), others expressed markers of GE-derived cells (e.g. *Gsx2* and *Dlx1*) (Fig. 4C) and some were those shown previously to co-express both (Fig. 3E). This suggested that cells in the aP state were labile, with some transiently activating elements of the eGC expression profile before reverting to a more normal trajectory (further evidence for this is presented below). Other *Pax6* cKO aPs had velocities directed towards eGCs, either eGC-Ps or eGC-Ns (Fig. 4B,C). These cells showed little or no expression of *Neurog2* and *Eomes* but strong expression of *Gsx2* and *Dlx1*, suggesting that they had become more highly committed to their aberrant fates.

**Figure 4:**
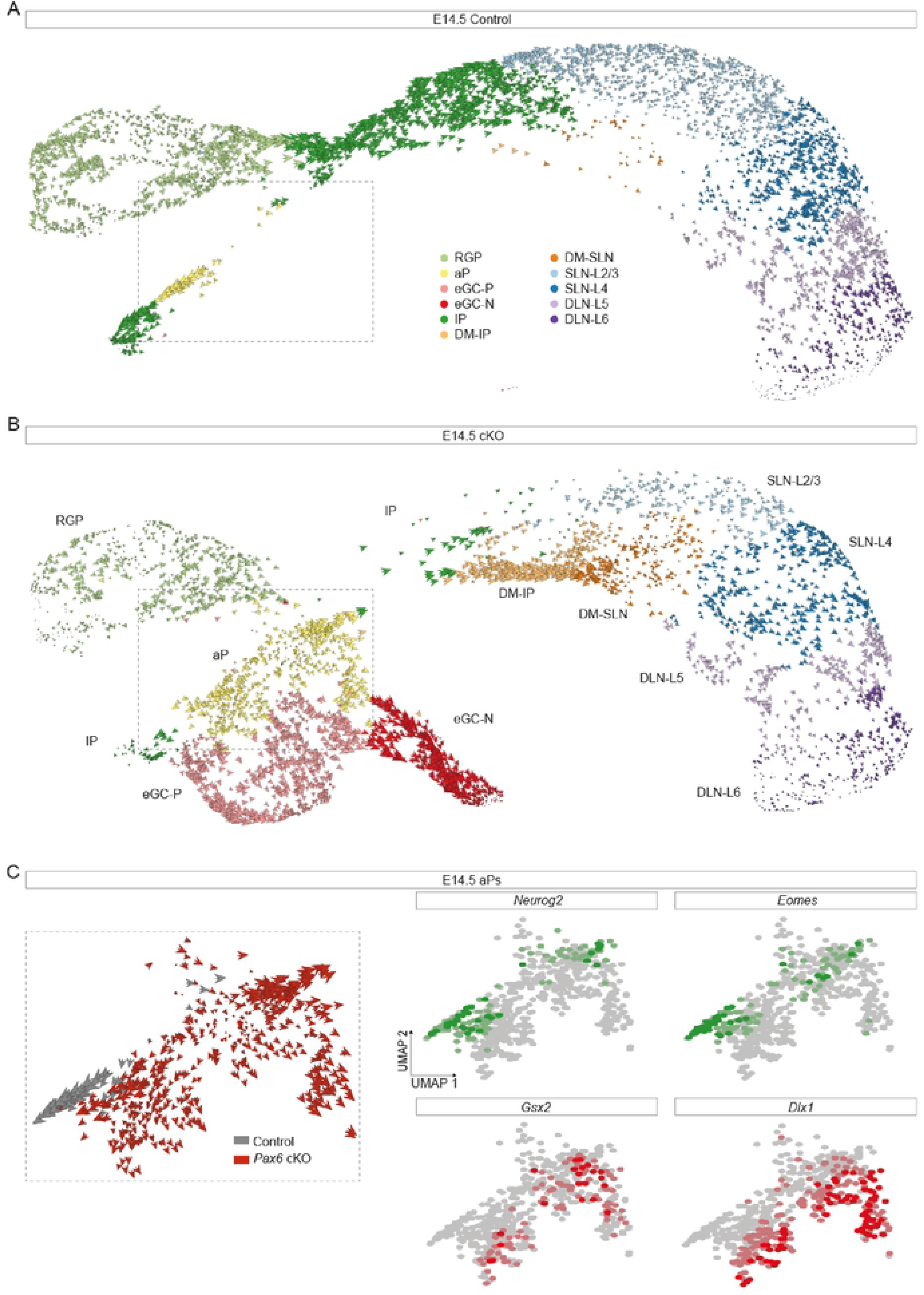
Development of abnormal cell lineages in *Pax6* cKO cortex. **(A,B)** RNA velocity analysis of individual E14.5 control and *Pax6* cKO cortical cells. Abbreviations: see Fig. 3A. **(C)** Enlargement of the boxed area in (B) showing RNA velocity analysis of individual aP control and *Pax6* cKO cells and UMAP plots of *Neurog2*, *Eomes*, *Gsx2* and *Dlx1* expression in aPs.

In summary, Pax6 removal appeared to have two major effects. First, it expanded (i) the proportion of progenitor cells in a labile state between other more highly specified progenitor states (aPs) and (ii) the proportion of cells with a relatively dorsomedial cortical identity (DMs). Second, it diversified the set of cell lineage progressions open to cortex-born cells: while some lineages remained similar to those adopted by normal cortex-born cells, others resembled those normally followed by cells outside the cortex.

### Spatiotemporal variation in the effects of Pax6 loss on selected gene expression

We next examined the effects of tamoxifen^E9.5^-induced *Pax6* deletion on spatial and temporal patterns of expression of key Pax6-regulated genes in cortical sections using in situ hybridization and immunohistochemistry. Comprehensive visualizations of the expression patterns of selected genes were obtained by combining data from serial sections such as those in Fig. 5A to generate surface-view reconstructions on representations of flattened cortical sheets (Fig. 5B,C,; for reconstruction method, see Fig. S5A).

**Figure 5:**
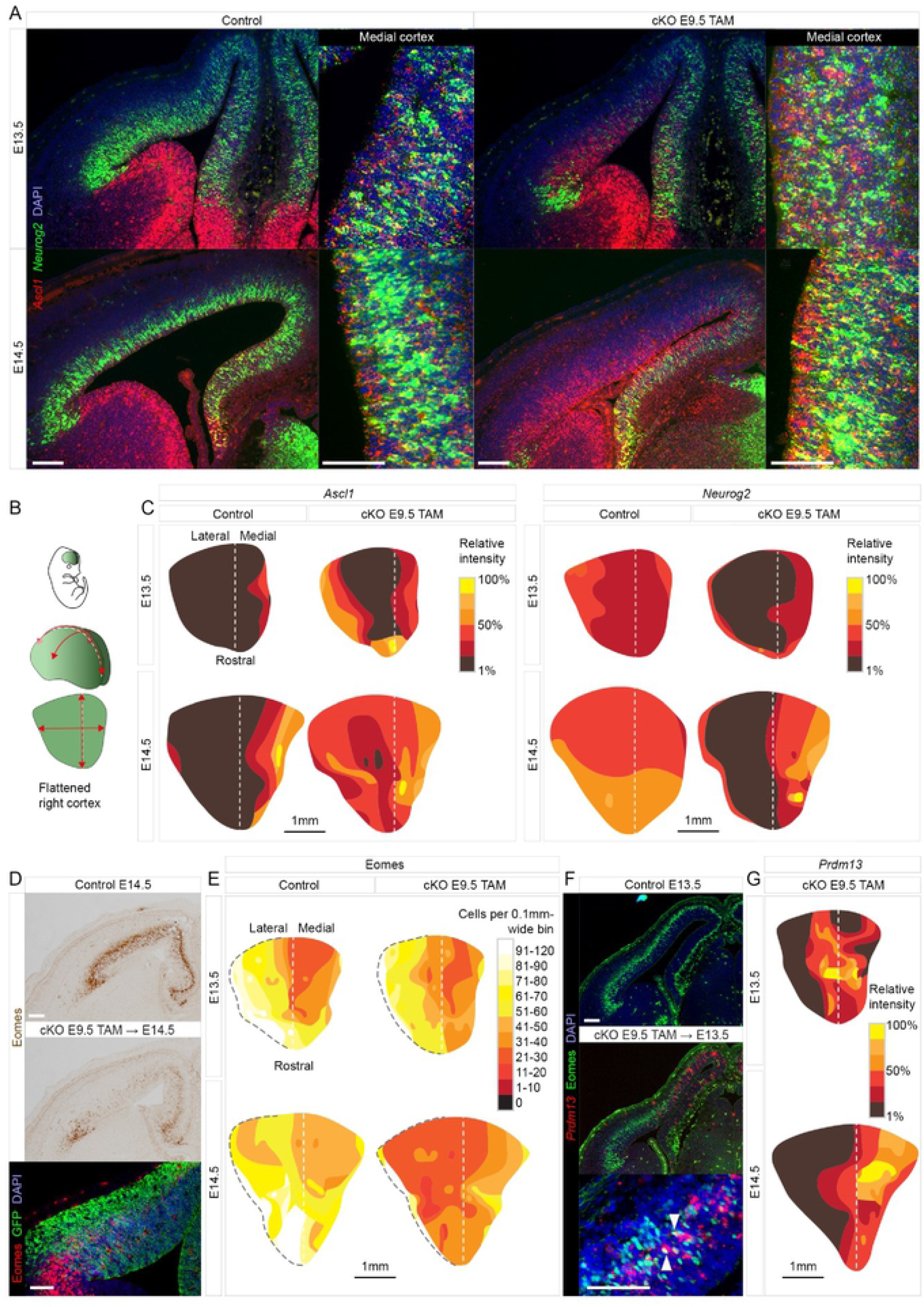
Regional changes in *Ascl1*, *Neurog2*, *Eomes* and *Prdm13* expression in *Pax6* cKO cortex. **(A)** In situ hybridizations for *Ascl1* and *Neurog2* in control and *Pax6* cKO cortex at E13.5 and E14.5. Scale bars: 0.1mm. **(B,C)** Flattened surface views of the cortex, oriented as in (B), to show the variation in the relative intensity of *Ascl1* and *Neurog2* staining across control and *Pax6* cKO cortex at E13.5 and E14.5. Method in Fig. S5A. **(D)** Colorimetric and fluorescence immunoreactivity for Eomes in control and *Pax6* cKO cortex at E14.5 (with GFP labelling *Emx1*-lineage cells). Scale bars: 0.1mm. **(E)** Flattened surface views of the cortex, oriented as in (B), to show the variation in the density of Eomes+ cells in control and *Pax6* cKO cortex at E13.5 and E14.5. Dashed black line: pallial-subpallial boundary. **(F)** Immunoreactivity for Eomes and in situ hybridization for *Prdm13* in control and *Pax6* cKO cortex at E13.5. Arrowheads indicate double-labelled cells. Scale bars: 0.1mm. **(G)** Flattened surface views of the cortex, oriented as in (B), to show the variation in the relative intensity of *Prdm13* staining in *Pax6* cKO cortex at E13.5 and E14.5.

*Ascl1* upregulation began in lateral-most cortex and spread across its entirety between E13.5 and E14.5, while *Neurog2* was downregulated in lateral cortex (except in the narrow ventral pallial domain where *Emx1^Cre^* is not expressed). Measurements of the proportions of cells expressing *Neurog2* or *Ascl1* with depth through the VZ and SVZ of lateral cortex showed that the distributions of *Neurog2*+ cells were replaced by similar distributions of *Ascl1*+ cells (Fig. S5B). Medial cortex, on the other hand, maintained levels of *Neurog2* expression that were similar to control (Fig. 5A,C) and contained a relatively high incidence of *Ascl1* and *Neurog2* co-expressing cells (Fig. 5A). Eomes downregulation was greater in lateral than in medial cortex (again, except in the narrow ventral pallial domain) (Fig. 5D,E).

Tamoxifen^E9.5^-induced *Pax6* deletion induced ectopic activation of *Prdm13* in a different pattern (Fig. 5F,G). *Prdm13*+ cells were located in the medial two thirds of the cortex, compatible with our scRNAseq analysis showing activation of *Prdm13* in DM cells (Fig. 3D). They were mainly in the SVZ, where they intermingled with Eomes+ cells, some of which co-expressed both genes (Fig. 5F). By E16.5, *Prdm13* expression remained detectable only in the most medial part of the cortex (Fig. S5C).

These results indicated that Pax6 loss had distinct effects on the expression of different genes and that the effects varied with cortical region and age.

### Pax6 loss induced eGC production in a distinct spatiotemporal pattern

We next examined cells that deviated to the eGC fate by probing for expression of *Gsx2*, *Dlx1* and *Gad1* (Fig. 6A-D). In normal cortical development, *Gsx2* becomes active only in small numbers of late-stage (E16.5 or older) cortical SVZ cells that generate cell types other than cortical neurons (Zhang et al., 2020) (these cells are seen in Fig. 6A: “Control E16.5”). Following tamoxifen^E9.5^, a wave of ectopic *Gsx2* activation was advancing rapidly across the cortex by E12.5. It began laterally and swept progressively further medially to occupy all parts of lateral cortex by E14.5, but did not extend all the way through medial cortex (Fig. 6A,E). We examined the extent to which this change depended on when tamoxifen was administered (evidence in Fig. S6A,B confirmed that tamoxifen administration at ages other than E9.5 also caused Pax6 removal from most RGPs within 3 days). We found similar distributions of Gsx2+ cells at E13.5 no matter whether tamoxifen was administered on E8.5, E9.5 or E10.5 (Fig. 6A,E) and even in E13.5 constitutive *Pax6^-/-^* mutants that had never expressed functional Pax6 (Fig. 6A). When we administered tamoxifen later, on E13.5, Gsx2+ cells were distributed throughout the entire lateral cortex 3 days later. This resembled the distributions at similarly late ages (E14.5-16.5) following early tamoxifen administration (E8.5-E10.5) and not the distributions 3 days after early tamoxifen administration (Fig. 6A).

**Figure 6:**
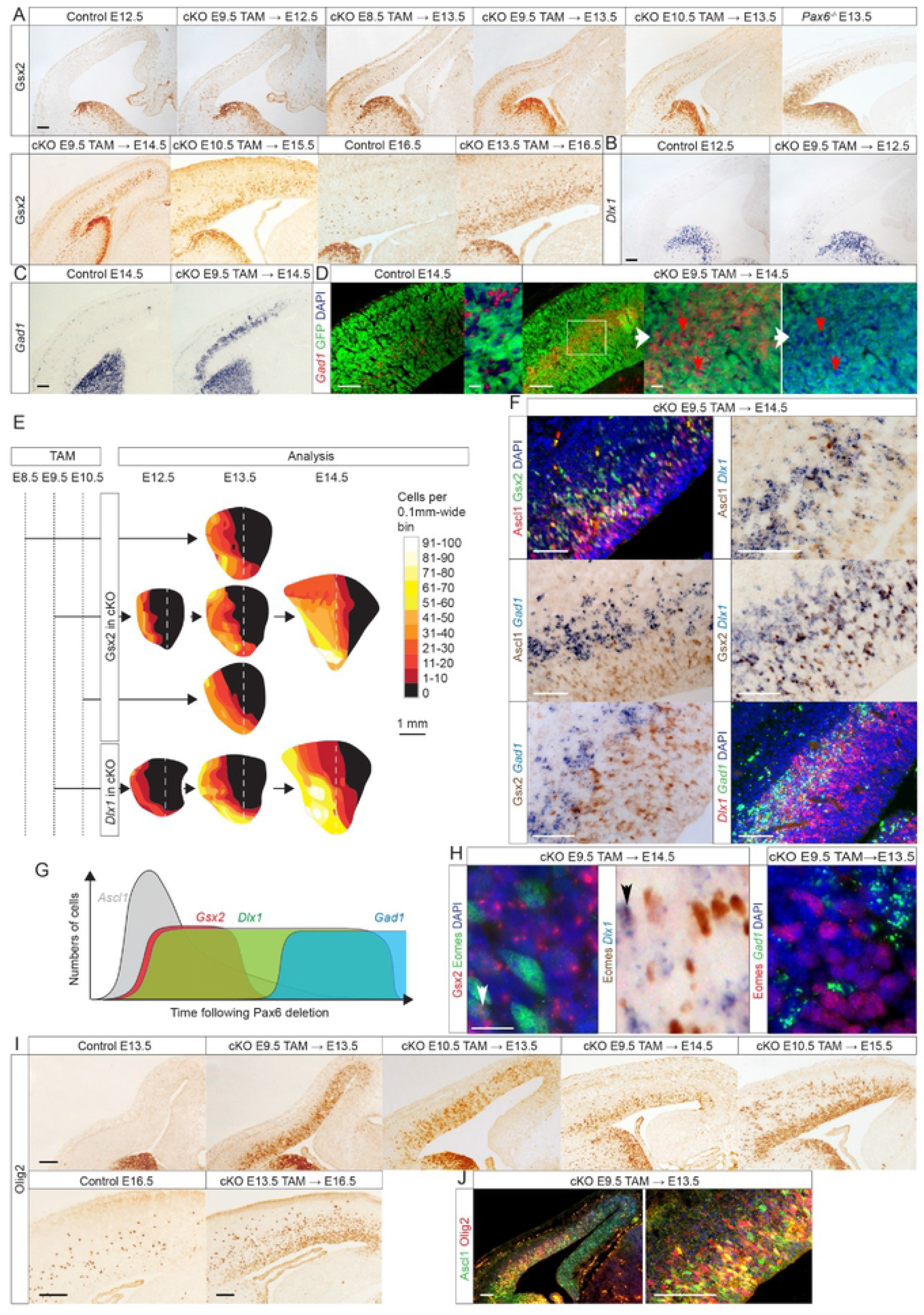
Pax6 loss induced ectopic gene expression in distinct spatiotemporal patterns. **(A)** Immunoreactivity for Gsx2 in control and *Pax6* cKO cortex between E12.5 and E16.5 following tamoxifen at E8.5, E9.5, E10.5 or E13.5 and in the cortex of E13.5 constitutive *Pax6^-/-^* embryos. Scale bar: 0.1mm. **(B,C)** In situ hybridizations for *Dlx1* and *Gad1* in control and *Pax6* cKO cortex at E12.5 and E14.5 following tamoxifen at E9.5. Scale bars: 0.1mm. **(D)** In situ hybridizations for *Gad1* and immunoreactivity for GFP in control and *Pax6* cKO cortex at E14.5 following tamoxifen at E9.5, showing enlargement of boxed area with and without *Gad1* staining. Red arrows in *Pax6* cKO cortex: examples of *Gad1*+ cells that were GFP-negative. Scale bars: 0.1mm and 0.01mm. **(E)** Flattened surface views of the cortex (method in Fig. S5A; oriented as in Fig. 5B), showing the densities of Gsx2+ and *Dlx1*+ cells in control and *Pax6* cKO cortex at E12.5, E13.5 and E14.5 following tamoxifen administration at E8.5, E9.5 or E10.5. **(F)** Fluorescence and colorimetric immunoreactivity for Ascl1 and Gsx2 and in situ hybridizations for *Dlx1* and *Gad1* in *Pax6* cKO lateral cortex at E14.5 following tamoxifen at E9.5. Scale bars: 0.1mm. **(G)** Summary of gene expression changes in (F) with time following Pax6 deletion. **(H)** Fluorescence and colorimetric immunoreactivity for Gsx2 and Eomes and in situ hybridizations for *Dlx1* and *Gad1* in *Pax6* cKO cortex at E13.5 and E14.5 following tamoxifen at E9.5. Scale bar: 0.01mm. **(I,J)** Colorimetric and fluorescence immunoreactivity for Olig2 and Ascl1 in control and *Pax6* cKO cortex between E13.5 and E16.5 following tamoxifen at E9.5, E10.5 or E13.5. Scale bars: 0.1mm.

We concluded that the spatial distribution of Gsx2+ cells depended mainly on cortical age rather than time elapsed since Pax6 removal, suggesting that cortical factors that change with age have important influences on the outcome of Pax6 removal.

Tamoxifen^E9.5^ induced a wave of ectopic *Dlx1* expression similar to that of Gsx2 expression, i.e. it was underway by E12.5 (Fig. 6B) and had spread through lateral cortex but only encroached to a limited extent into medial cortex by E14.5 (Fig. 6E). Tamoxifen^E9.5^ also led to the generation of a large population of *Gad1*+ cells in the lateral cortex (Fig. 6C,D). Most of these cells were cortically-derived (i.e. they were GFP+ *Emx1*-lineage) but they were intermingled throughout their domain with other *Gad1*+ cells that were GFP-negative subcortically-generated immigrants (arrows in Fig. 6D).

In the VZ and SVZ of *Pax6*-deleted lateral cortex, *Gsx2*, *Dlx1* and *Gad1* were activated by partially overlapping bands of cells centred progressively further basal to the ventricular surface (Fig. 6F). The Gsx2+ and *Dlx1*+ bands overlapped the basal side of the Ascl1+ band and the *Dlx1*+ band extended further basally than the Gsx2+ band. This was followed by the *Gad1*+ band, which showed considerable overlap with the *Dlx1*+ band but less overlap with the Ascl1+ and Gsx2+ bands (summarized in Fig. 6G). Where domains of expression overlapped, co-expressing cells were frequent. Small proportions of Gsx2+ or *Dlx1*+ cells co-expressed Eomes (arrows in Fig. 6H), in agreement with findings in our scRNAseq data (Figs. 3E, 4C).

We concluded that the production of eGCs unfolded in a distinct spatiotemporal pattern in mainly lateral cortex.

### Pax6 loss induced ectopic Olig2 expression largely independently of eGC production

We then examined the pattern of ectopic cortical activation of *Olig2*, which is expressed in progenitors that generate cortical interneurons and oligodendrocytes, is normally restricted to the embryonic subpallium at around E13.5 (Fig. 6I: “Control E13.5”) and later spreads as Olig2+ cells migrate into the cortex (Fig. 6I: “Control E16.5”) (Kessaris et al., 2006; Miyoshi et al., 2007). Our scRNA-seq data indicated that *Olig2* was not specifically marking eGCs, but was expressed by many additional cell types including RGPs, aPs, IPs and differentiating cells in *Pax6* cKOs (Figs. S3, S6C). Its ectopic spatiotemporal activation pattern differed from that of *Gsx2*, *Dlx1* and *Gad1* to the extent that it appeared throughout the entire lateral cortex earlier, by E13.5, but was similar in showing relatively little activation in medial cortex, even at later ages (Fig. 6I). The domain of Olig2 activation was similar in E13.5-E16.5 embryos regardless of whether tamoxifen was given at E9.5, E10.5 or E13.5. In lateral cortex, many progenitors co-expressed *Olig2* and *Ascl1* (Fig. 6J); this was supported by scRNAseq data showing that 51.8% and 67.3% of *Olig2*+ cells expressed *Ascl1* at E13.5 and E14.5 respectively. Nevertheless, our *Pax6* cKO E14.5 scRNAseq data detected *Olig2* co-expression in only a small proportion (9.6%) of cells expressing eGC markers *Gsx2*, *Dlx1*, and *Gad1* (Fig. S6C).

These findings suggested that the Pax6-loss-induced activation of *Olig2* and of eGC-expressed genes such as *Gsx2*, *Dlx1* and *Gad1* occurred largely independently. They provided further evidence of spatiotemporal variation in the effects of Pax6 loss on the ectopic activation of different genes.

### The eGCs were highly proliferative

Our scRNAseq data indicated the existence of a substantial population of proliferating eGCs in E14.5 *Pax6* cKO cortex. This was demonstrated, for example, by the rising levels of the mitotic marker *Mki67* along the inferred pseudotime trajectory of the lineage leading to eGC-P generation (Fig. 7A; trajectories were obtained using Slingshot and tradeSeq: Street et al., 2018; Tritschler et al., 2019; Van den Berge et al., 2020). To test this conclusion further, we used the *Emx1-Cre^ERT2^* allele with tamoxifen^E9.5^ to delete *Pax6* and then labelled proliferating cells by administering the S phase marker 5-ethynyl-2’-deoxyuridine (EdU) at E13.5, 30 minutes before death (Fig. 7B). We reacted sections for EdU and Gsx2, a marker of early eGCs (and also for GFP from a *Btg2*-GFP transgene that was incorporated into the mice for reasons given below) (Fig. 7C). Most Gsx2+ cells were in S phase (mean = 59.0% ±3.4 sd; n = 5 embryos from separate litters; counts were from 20 equally-spaced coronal sections through the cortex for each embryo; Source Data Table 2A), confirming their high level of proliferation.

**Figure 7:**
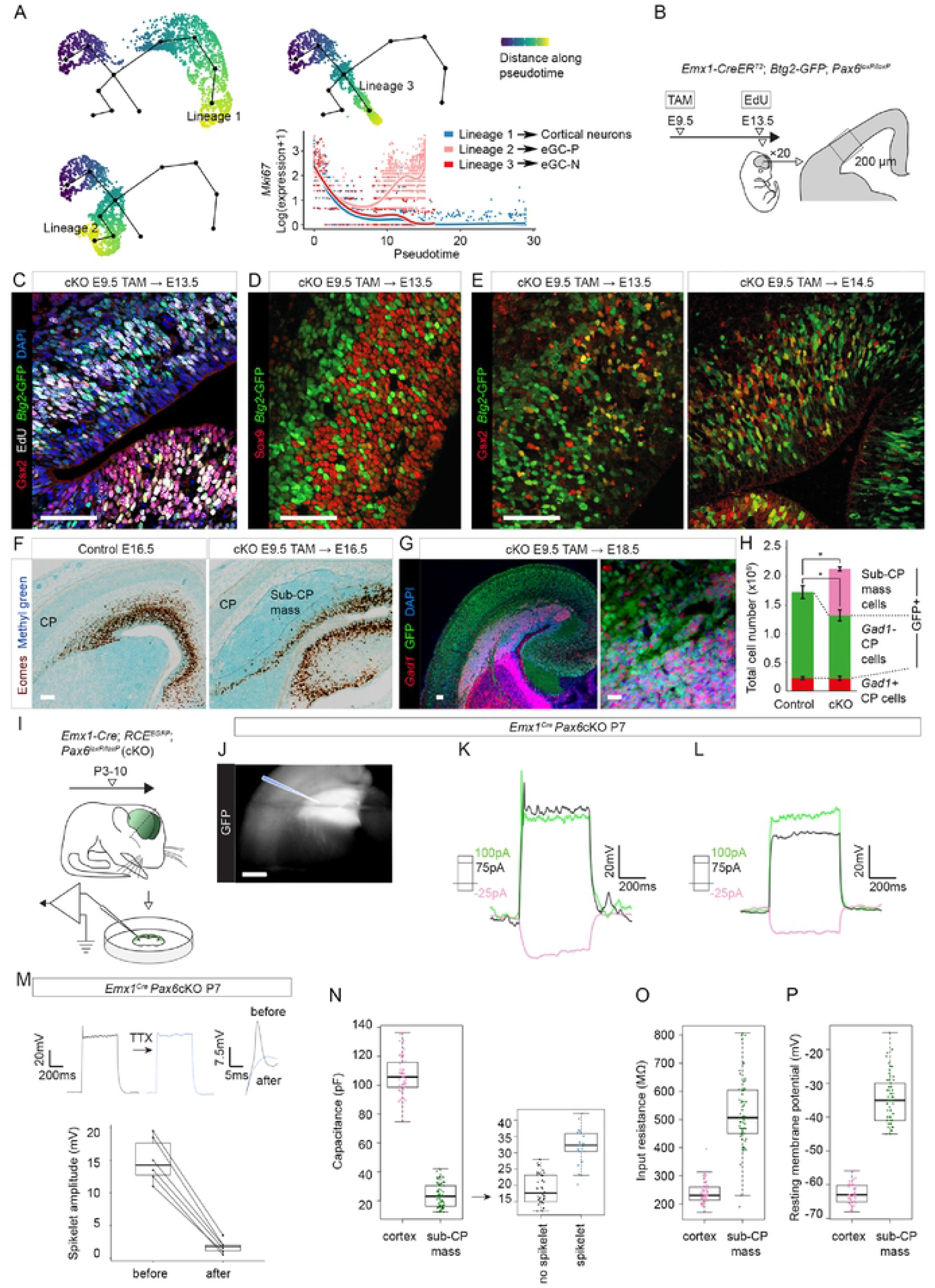
The proliferation, coalescence and electrophysiological properties of *Gsx2*-lineage eGCs. **(A)** Major pseudotemporal trajectories inferred from E14.5 *Pax6* cKO scRNAseq data (one leading to eGC-Ps, one to eGC-Ns and one to cortical glutamatergic neurons) and expression of the marker of proliferating cells, *Mki67*, along each. **(B)** The experimental procedure for EdU labelling. The *Emx1-Cre^ERT2^* allele with tamoxifen^E9.5^ was used to delete *Pax6* (embryos carried a *Btg2*-GFP transgene); 5-ethynyl-2’-deoxyuridine (EdU) was given at E13.5, 30 minutes before death; 20 coronal sections equally spaced through the brain were immunoreacted for EdU, Gsx2 and GFP; counts were made in the boxed area. **(C)** Fluorescence quadruple-staining for Gsx2, EdU, GFP (marking *Btg2*-expressing cells) and DAPI in E13.5 *Pax6* cKO cortex after the procedure in (B). Scale bar: 0.1mm. **(D)** Fluorescence double-staining for Sox9 and GFP (marking *Btg2*-expressing cells) in E13.5 *Pax6* cKO cortex after tamoxifen^E9.5^. Scale bar: 0.1mm. **(E)** Fluorescence double-staining for Gsx2 and GFP (marking *Btg2*-expressing cells) in E13.5 and E14.5 *Pax6* cKO cortex after tamoxifen^E9.5^. Scale bar: 0.1mm. **(F)** Eomes immunoreactivity and methyl green counterstaining in control and E16.5 *Pax6* cKO cortex after tamoxifen^E9.5^. Scale bar: 0.1mm. **(G)** Fluorescence immunoreactivity for GFP (*Emx1*-lineage) and in situ hybridization for *Gad1*+ cells in E18.5 *Pax6* cKO cortex after tamoxifen^E9.5^. Scale bars: 0.1mm and 0.01mm. **(H)** Quantifications of the total numbers of *Gad1*+ cells in the lateral cortical plate (CP; red: they were GFP-, subcortically-derived), of *Gad1*-cells in the CP (green: GFP+, cortical-born) and of cells in the *Pax6* cKO sub-CP masses (pink: *Gad1*+, GFP+) in control and *Pax6* cKO E18.5 embryos after tamoxifen at E9.5 (for quantification method, see Fig. S7F). Total numbers of cells were greater in *Pax6* cKO cortex (p<0.05) and numbers of lateral CP cells were reduced (p<0.02) (averages ± sem; Student’s paired t-tests; n=4 embryos of each genotype, from 4 independent litters) (Source Data Table 3). **(I)** The experimental procedure for electrophysiology (J-P). The *Emx1-Cre* allele was used to delete *Pax6*; embryos carried a GFP reporter transgene. Recordings were from sub-CP masses at P3-10. **(J)** Sub-CP mass in P7 slice prepared for electrophysiology: the cortex was GFP+ and the sub-CP mass was intensely so. Scale bar: 0.5mm. **(K,L)** Examples of responses of sub-CP mass cells to current injections (square steps, magnitudes colour-coded, 500ms duration). Membrane voltages were held at -70mV. Some cells produced small spikelets (J), others did not (K). **(M)** Tetrodotoxin (TTX; 300nM) reduced spikelet amplitudes; examples of entire response and spikelet alone before and after TTX application; effects of TTX were significant (p=0.035, Wilcoxon signed rank test; n=6 cells) (Source Data Table 4A). **(N-P)** Passive electrical properties of P3-P10 sub-CP mass cells compared to P5-P7 cortical cells from layer 5 of somatosensory area 1 (n=66 sub-CP mass cells, Source Data Table 4B; n=49 cortex cells, data for these CP cells are in Table S5). Sub-CP mass cells had significantly lower capacitance (p=2.2x10^-16^, Mann-Whitney test) and significantly higher input resistance (p=2x10^-10^, Mann-Whitney test) and resting membrane potential (p=2.2x10^-16^, Mann-Whitney test). For capacitance, values were significantly higher among sub-CP mass cells that produced spikelets (n=22 cells; n=44 produced no spikelet) (p=1.9x10^-9^, Mann-Whitney test).

We then studied the types of division that *Pax6* cKO cortical progenitors made. Previous work has shown that RGPs (Sox9+) and IPs (Eomes+) produce either post-mitotic neurons or new progenitors (Noctor et al., 2004; Gao et al., 2014; Vasistha et al., 2015; Mihalas and Hevner, 2018). Progenitors of the latter type, often described as proliferative progenitors, do not express the anti-proliferative gene *Btg2*; others, often described as neurogenic, do express *Btg2* (Iacopetti et al., 1999; Haubensak et al., 2004). We used the *Btg2*-GFP transgene (Haubensak et al., 2004) with immunohistochemistry to identify neurogenic progenitors (Fig. 7D,E). Many Gsx2+ cells expressed *Btg2* at E13.5 and E14.5, but a sizeable minority did not. Quantification in E13.5 tissue sections showed that 68.1% ±6.5 (sd) of Gsx2 protein-expressing cells were also *Btg2*-expressing (n=5 embryos from separate litters; counts were from 20 equally-spaced coronal sections through the cortex for each embryo; Source Data Table 2A). This was similar to scRNAseq data, which showed *Btg2* expression in 76.2% and 74.0% of *Gsx2*+ cells at E13.5 and E14.5 respectively. These data indicated that, overall, about a quarter of the cortical cells that activated *Gsx2* were proliferative (i.e. *Btg2*-non-expressing; their daughters would divide at least once more). The emergence in *Pax6* cKO cortex of substantial numbers of repeatedly and rapidly dividing progenitors caused a large expansion of the eGC population, described in the next section.

### Transient subcortical masses of eGCs formed beneath lateral cortex

Abnormal collections of *Gad1*+ cells coalesced beneath the cortical plate (CP) and superficial to the reduced population of Eomes+ cells in *Pax6* cKO lateral cortex between E14.5 and E16.5 (Fig. 7F; Fig. S7A). We refer to them here as sub-CP masses. Their expression of GFP, which indicated they were *Emx1*-lineage (Fig. 6C,D; Fig. 7G), combined with their *Gad1* positivity identified these cells as eGCs (Fig. 3A,D). Based on findings described above, it was likely that many of them had gone through an early transient phase of *Gsx2* expression (Fig. 6G). To confirm this, we lineage-traced cells using *Gsx2-Cre* (Kessaris et al., 2006) and the GFP Cre-reporter allele (Miyoshi et al., 2010).

Since we could not use *Gsx2-Cre* in combination with *Emx1-Cre^ERT2^*, these experiments were done in *Pax6^Sey/Sey^* (*Pax6^-/-^*) constitutive mutants (Fig. S7B-D), whose pattern of Gsx2 expression was similar to that in *Pax6* cKO cortex (Fig. 6A). We confirmed that *Gsx2*-lineage cells lost their Gsx2 protein as they moved away from the ventricular surface, activating the *Gsx2-Cre*-activated GFP reporter but no longer Gsx2 protein (Fig. S7B). In E14.5 control cortex, *Gsx2*-lineage cells were scattered and had the elongated appearance associated with migration from the Gsx2+ subpallium (Fig. S7B). In line with our prediction, most cells in the E16.5 sub-CP masses were *Gsx2*-lineage and most of their cells were *Gad1*+ (Fig. S7C). In these experiments, we also observed a population of GFP+ *Gad1*-negative neurons scattered through the CP of E16.5 *Pax6^-/-^* but not control lateral cortex: many of these neurons had the shape and apical dendrite associated with young cortical excitatory neurons (Fig. S7D). This result agreed with a prediction from our scRNAseq analysis that some cells that first expressed markers of eGCs later reverted to a cortical excitatory neuronal fate (Fig. 4C), reinforcing the suggestion of instability in the identities of *Pax6* cKO cells exiting the RGP state.

We returned to using *Pax6* cKOs (tamoxifen^E9.5^) to gain further information on the development of eGCs and sub-CP masses. Only extremely rarely did we find examples of *Emx1-*lineage (i.e. GFP+) *Gad1*+ cells in the CP of *Pax6* cKOs (an example is shown in Fig. S7E), indicating that the vast majority of eGCs were unable to contribute to the CP. We estimated the total numbers of subcortically-derived *Gad1*+ interneurons (i.e. non-*Emx1*-lineage, GFP-negative), the total numbers of cells contained in the sub-CP masses and the total numbers of all GFP+ cells in the lateral CP of control and *Pax6* cKO cortex at E18.5 (Fig. 7G,H; methodology in Fig. S7F). In *Pax6* cKOs, the numbers of *Gad1*+ GFP-negative cells in the lateral CP remained unchanged, indicating that immigration of subcortical *Gad1*+ cells into the lateral CP had proceeded normally. The numbers of GFP+ cells in the lateral CP were significantly reduced (Fig. 7H; p<0.02; Student’s paired t-test). Adding the numbers of cells in the sub-CP masses to the numbers in the lateral CP revealed that, overall, significantly more cells in total were generated in *Pax6*-deleted than in control lateral cortex (Fig. 7H; p<0.05; Student’s paired t-test). This was explicable by the switch of a significant proportion of cortical progenitors to the generation of highly proliferative eGCs that populated the sub-CP masses.

The sub-CP masses were no longer visible by postnatal day 34 (P34) (Fig. S7G). This was most likely due to the death of their cells. The proportion of cells expressing the apoptosis marker caspase-3 was much higher in the sub-CP masses than in overlying CP (Fig. S7H). It rose from 1.0% (±0.85 sd) at E14.5-16.5 to 7.1% (±4.6 sd) at P10, whereas it remained consistently very low in control cortex (mean = 0.29% ±0.21 sd, all ages combined) (Source Data Table 2B). We concluded that the very high level of proliferation among *Gsx2*-lineage cells in the lateral *Pax6* cKO cortex generated large sub-CP masses of eGCs that were eventually removed through cell death.

### Sub-CP mass cells showed immature electrophysiological properties

We tested whether sub-CP mass cells developed electrophysiological properties resembling those of interneurons by making whole-cell current-clamp recordings at P5-P10 (Fig. 7I) (Connors and Gutnick, 1990; Markram et al., 2004; Scala et al., 2021). These ages encompassed those by which normal cortical neurons have acquired the ability to generate individual or trains of action potentials (APs) in response to depolarizing stimuli (Luhmann et al., 2000; Daw et al., 2007; Yang et al., 2018). The sub-CP masses were easily identified in slices at all ages by their intense GFP expression (Fig. 7J).

The properties of the sub-CP mass cells were similar across the range of ages studied here. None of them generated mature APs. A third (22/66) produced either spikelets (spikelet peak < 10mV; spikelet amplitude = 5-25mV; little or no afterhyperpolarization, AHP; Fig. 7K) or, in two cases, underdeveloped APs (peak amplitude > 30mV and AHP > 15mV). Most (44/66) produced neither (Fig. 7L). Spikelet amplitudes were reduced by ∼90% following the addition of 300nM tetrodotoxin (TTX), which blocks the voltage gated Na^+^ channels responsible for the rising phase of the AP (Elliott and Elliott, 1993; Bean, 2007) (Fig. 7M), suggesting that spikelets were immature APs. One possibility was that the cells that produced spikelets were eGC-Ns, whereas those that did not were eGC-Ps.

The sub-CP mass cells had much lower capacitances and higher input resistances (R_in_s) and resting membrane potentials (RMPs) than P5-P7 cortical neurons recorded in layer 5 of primary somatosensory cortex (Fig. 7N-P; Fig. S9D-F; Table S5). Their relatively low capacitances were a sign that they had relatively small somas (Fig. 7N). When we split them into those that produced spikelets and those that did not we found that the former had higher capacitances, indicating that they were slightly larger (Fig. 7N). The relatively high R_in_s and RMPs of the sub-CP mass cells, neither of which differed significantly between cells that did or did not generate spikelets, were likely attributable to immaturity in the numbers of ion channels in their cell membranes (LoTurco et al., 1991; Destexhe et al., 2003; Tyzio et al., 2003; Yang et al., 2018).

We concluded that although the transcriptomes of these cells showed progress towards a GABAergic interneuron fate, they were unable to develop corresponding cellular properties. Whether this was because they had a cell autonomous inability to mature and/ or a problem with the environment in which they found themselves was not tested here.

### The production of eGCs did not depend on *Gsx2* activation

We next questioned whether early activation of *Gsx2* by eGCs contributed to their activation of genes such as *Dlx1* and *Gad1* and their repression of *Neurog2* and *Eomes*, as it does in the GEs (Toresson et al., 2000; Corbin et al., 2000; Yun et al., 2001; Waclaw et al., 2009). We carried out tamoxifen^E9.5^-induced *Pax6* and *Gsx2* cortex-specific co-deletion (double conditional KO, or dcKO; Fig. 8A; Fig. S8A). Co-deletion of *Gsx2* did not prevent the production of *Dlx1*+ and *Gad1*+ cells in similar numbers as in *Pax6* single cKOs (Fig. 8B,C). It had no detectable effect on activation of Ascl1 expression and did not reverse the Pax6-loss-induced loss of *Neurog2* from the bulk of the lateral cortex (Fig. S8B,C). While Gsx2 protein was not detected in dcKOs (Fig. 8B), mRNA from *Gsx2* exon 1 was (the deletion removed the homeodomain-encoding exon 2; Fig. S8A; Waclaw et al., 2009), allowing us to use in situ hybridization to recognize cortical cells that had activated the *Gsx2* gene even in dcKOs. Using this approach, we found no evidence that loss of Gsx2 protein from *Gsx2*+ cells caused them to upregulate Eomes expression (Fig. 8D). Nor did it cause cortical activation of *Gsx1*, which occurs in the dorsal LGE (dLGE) following Gsx2 removal (Fig. S8D) (Yun et al., 2001; Toresson and Campbell, 2001; Wang et al., 2013).

**Figure 8:**
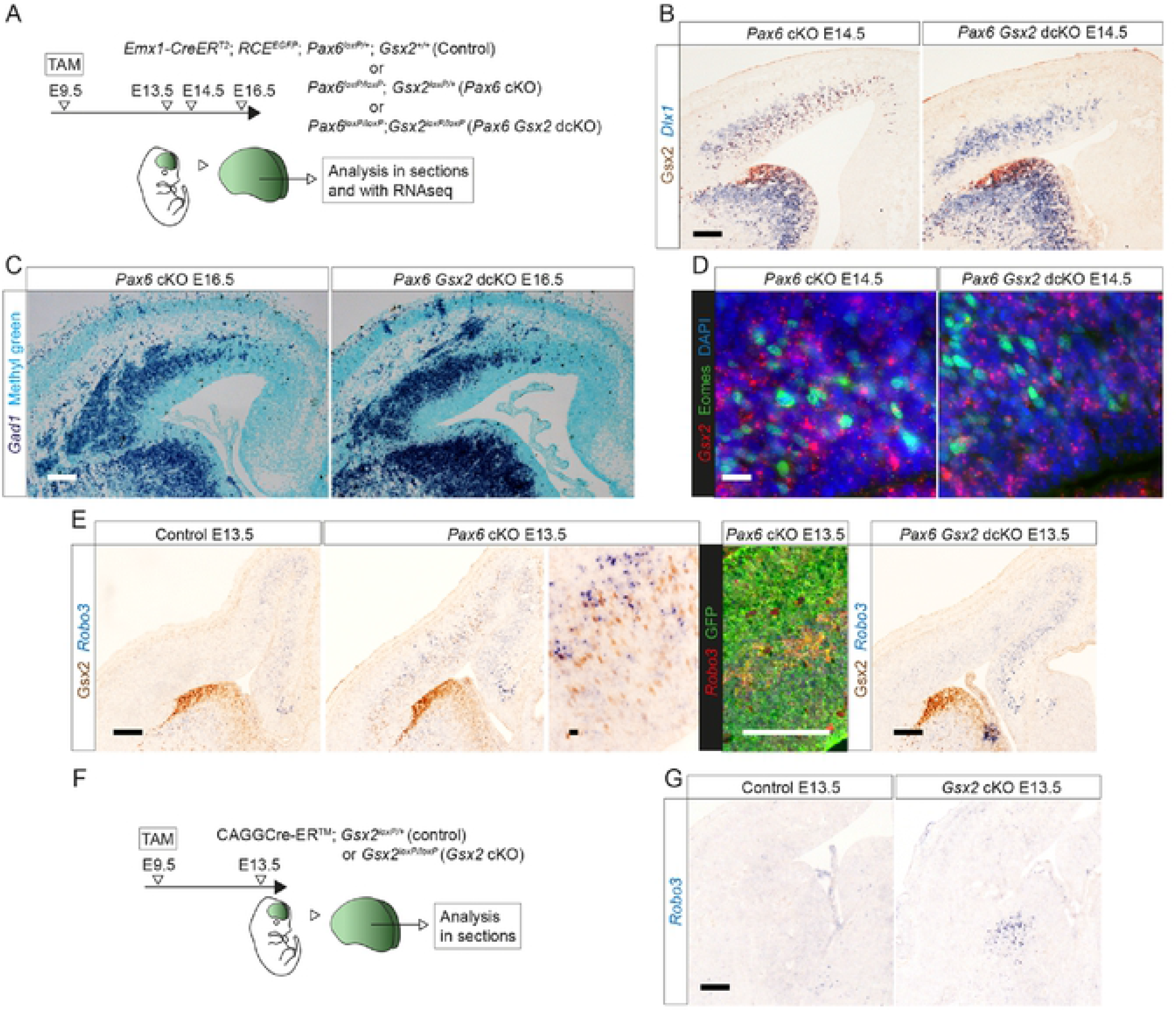
The production of eGCs did not require Gsx2 activation. **(A)** The experimental procedure for (B-E): tamoxifen (TAM) was administered at E9.5 to generate control embryos with functional alleles of both *Pax6* and *Gsx2*, a single cKO of *Pax6* or a double cKO (dcKO) of *Pax6* and *Gsx2*; brains were analysed at E13.5, E14.5 or E16.5. **(B)** Colorimetric immunoreactivity for Gsx2 and in situ hybridization for *Dlx1* in *Pax6* cKO and *Pax6 Gsx2* dcKO at E14.5. Scale bar: 0.1mm. **(C)** Colorimetric in situ hybridization for *Gad1* in *Pax6* cKO and *Pax6 Gsx2* dcKO at E16.5. Scale bar: 0.1mm. **(D)** Fluorescence in situ hybridization for *Gsx2* and immunoreactivity for Eomes in *Pax6* cKO and *Pax6 Gsx2* dcKO at E14.5. Scale bar: 0.01mm. **(E)** Colorimetric and fluorescence in situ hybridizations for *Robo3*, colorimetric immunoreactivity for Gsx2 and fluorescence immunoreactivity for GFP in control, *Pax6* cKO and *Pax6 Gsx2* dcKO at E13.5. Scale bars: 0.1mm and 0.01mm. **(F)** The experimental procedure for (G): tamoxifen (TAM) was administered at E9.5 to generate control embryos with a functional allele of *Gsx2* or a cKO of *Gsx2* throughout the embryo; brains were analysed at E13.5. **(G)** Colorimetric in situ hybridizations for *Robo3* in control and *Gsx2* cKO at E13.5. Scale bar: 0.1mm.

We obtained further evidence that Gsx2 loss had a very limited effect on the development of eGCs using RNAseq to compare gene expression levels in dcKOs versus *Pax6* single cKO cortex at E13.5. Raw data is available at the European Nucleotide Archive accession number PRJEB21105. This analysis found only two genes were significantly upregulated following deletion of Pax6 alone and significantly downregulated by co-deletion of Gsx2 (adjusted p<0.05; Fig. S8E,F), namely *Gsx2* itself and *Robo3*. Fifteen other genes showed small, significant differences in expression levels in dcKO compared to *Pax6* single cKO cortex: all were increased in dcKOs and only one of them, *Zic4*, was significantly affected (slightly increased) in *Pax6* single cKOs (Fig. S8F).

We examined the effect of *Pax6* and *Gsx2* co-deletion on *Robo3* expression more closely. In controls expressing both Pax6 and Gsx2, there was a declining medial-to-lateral gradient of *Robo3* expression across the cortex, similar to that described previously for Robo3 protein (Fig. 8E) (Barber et al., 2009). In *Pax6*-deleted cortex, there was additional *Robo3* expression in cells partly intermingled with, and partly basal to, the Gsx2+ cells in lateral cortex (Fig. 8E). Few cells appeared to be double-labelled, agreeing with detection by scRNAseq of Gsx2 in only 0.2% of *Robo3*+ cells at E13.5 and 6.2% at E14.5. These *Robo3*+ cells were cortex-born (i.e. they were GFP+) (Fig. 8E). This additional *Robo3* expression in lateral cortex was reduced in *Pax6*-*Gsx2* dcKO cortex (Fig. 8E).

We then examined how *Robo3* expression was affected by Gsx2 in its normal domain of expression in the GEs by using a *CAGG-Cre^ERTM^* allele with tamoxifen^E9.5^ to delete *Gsx2* throughout the entire embryo (Fig. 8F). This increased *Robo3* expression in the LGE, where Gsx2 is normally strongly expressed. This effect was opposite to that caused by *Gsx2* deletion in eGCs (Fig. 8G).

We concluded that although Gsx2 was one of the earliest genes expressed in eGCs, the expression of other eGC marker genes did not require its expression, suggesting that Pax6 deletion has parallel effects on multiple eGC marker genes. Moreover, the transcriptional responses of eGCs to Gsx2 loss were unlike those of normal Gsx2-expressing GE cells, which might reflect an intrinsic difference in the nature of the two cell types and/ or a difference in their extracellular environments.

### Pax6 was not required in RGPs for their production of cortical excitatory neurons

As described above (Fig. S1B-F), almost all Pax6 protein was lost across all *Emx1*-expressing cortex by E12.5 in *Pax6* cKOs generated by tamoxifen^E9.5^. This near-universal loss had a near-universal effect on the expression of some genes. For example, some genes with strong expression across control E13.5 cortex became undetectable in most *Pax6* cKO cortical regions by E13.5 (Fig. 9), indicating that most cKO cells altered their gene expression at least to some extent. Nevertheless, Pax6 loss from RGPs did not stop them from generating large numbers of cells that were competent to migrate into the CP (in agreement with previous studies: Caric et al., 1997; Tuoc et al., 2009) (Fig. S9A,B), where they differentiated into deep and superficial layer neurons (our scRNAseq data; Fig. 3A). Moreover, *Pax6* cKO CP contained Slc17a7 (Vglut1), a specific marker of glutamatergic neurons and synapses, distributed in a similar pattern to that in controls (Fig. S9C).

**Figure 9:**
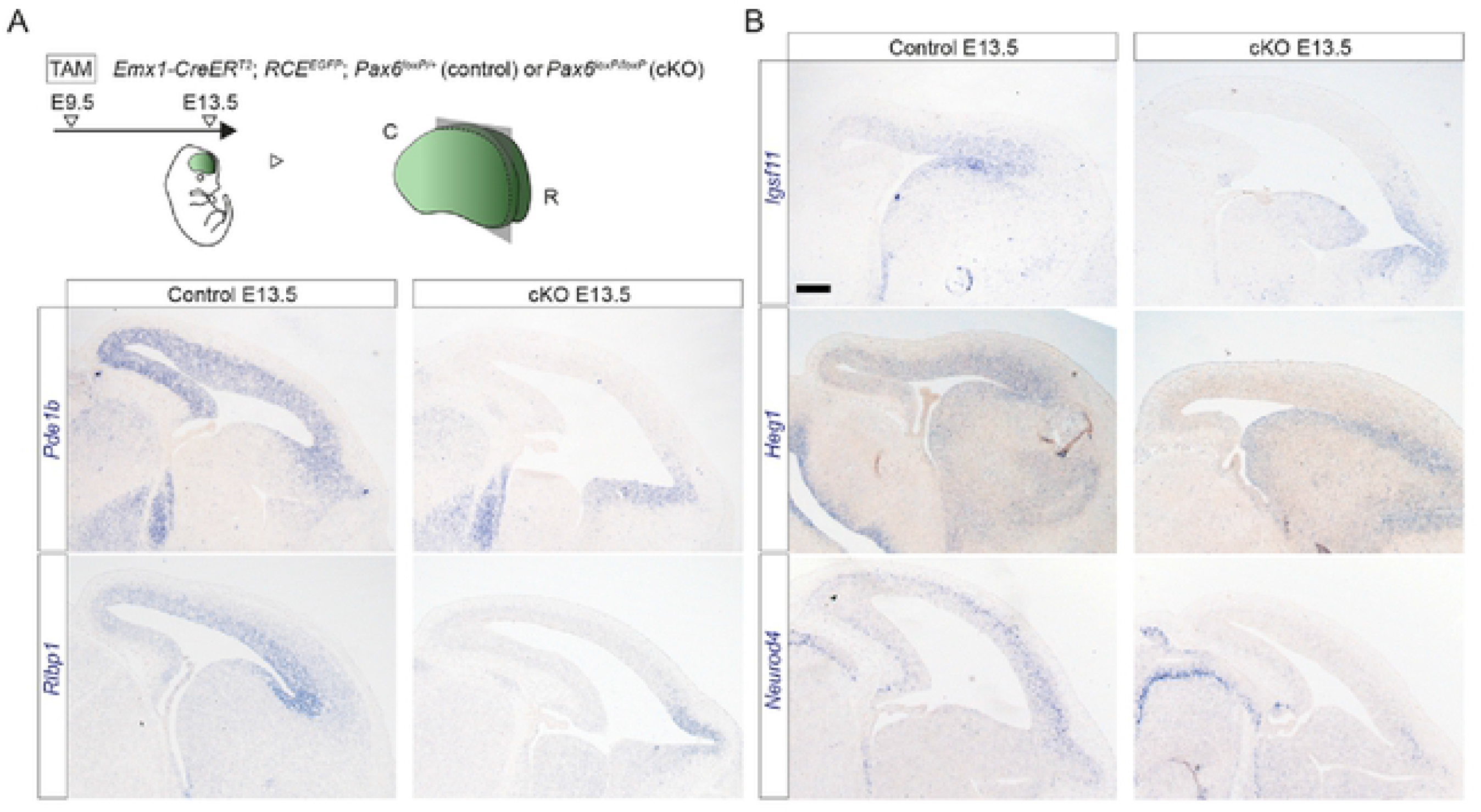
Widespread loss of gene expression in Pax6 cKOs. **(A)** Tamoxifen (TAM) was administered at E9.5 to generate control and *Pax6* cKO embryos; brains were sectioned sagittally at E13.5. **(B)** In situ hybridizations for *Pde1b*, *Rlbp1*, *Igsf11*, *Heg1* and *Neurod4* in E13.5 control and *Pax6* cKO cortex. Scale bar: 0.1mm.

Further evidence that Pax6 removal from RGPs did not prevent their generation of apparently normal cortical neurons came from whole-cell current-clamp recordings from GFP+ cells in layers 2/3 and 5 in primary somatosensory cortex (S1) (Fig. S9D-H). We detected no effects of genotype on the individual intrinsic functional properties of recorded cells (Table S5), nor were cells separated by genotype using unsupervised hierarchical agglomerative clustering based on the cells’ property profiles (Fig. S9G,H; Andjelic et al., 2009; Soldado-Magraner *et al*., 2019). Of the 73 GFP+ cells recorded in *Pax6*cKO cortex, one (in layer 2/3) showed properties compatible with those of fast spiking interneurons (Fig. S9I; Daw et al., 2007; Butt et al., 2008). It is possible that this represented a rare example of an eGC contributing to the cortical layers (see above; Fig. S7E). No such cells were found in controls (n=70 cells).

### Why *Pax6* deletion altered the fates of only some cortical cells: a hypothesis

We then turned to the question of why some cortical cells switched fate while others did not after Pax6 deletion from cortical progenitors. A parsimonious explanation was that Pax6 loss increased the *potential* for all RGPs to generate inappropriate cell lineages, but *triggering* this required additional, extracellular factors. Systematic cross-cortical variations in the types and levels of these factors might have been responsible for generating the spatiotemporal patterns of normal and abnormal specification seen after *Pax6* deletion. We set out to test this idea.

### Immigrating cortical interneurons enhanced the misspecification of *Pax6* cKO cortical cells

The striking similarity between the spatiotemporal characteristics of the wave of eGC production and the wave of subcortically-generated interneuron immigration (Fig. S10A), which was not disrupted by Pax6 removal (see above), suggested that the immigrating interneurons might have been one source of extracellular factors triggering abnormal specification among *Pax6* cKO cortical cells. To test this possibility, we removed subcortical tissue from one side of cultured coronal slices of E13.5 *Pax6* cKO (tamoxifen^E9.5^) telencephalon to prevent further interneuron influx and compared the production of Gsx2+ cells on the two sides after 48h in culture, using the GFP reporter to mark cells of cortical origin (Fig. 10A,B). The numbers of subcortically-generated interneurons (i.e. GFP-negative *Gad1*+ cells) were ∼4 times higher on the intact side (Fig. 10C,D), as anticipated from previous work using this approach (Anderson et al., 1997). Proportions of GFP+ Gsx2+ cells were several times higher on the intact side, with significant differences in the more lateral parts of cortex (Fig. 10E,F).

**Figure 10:**
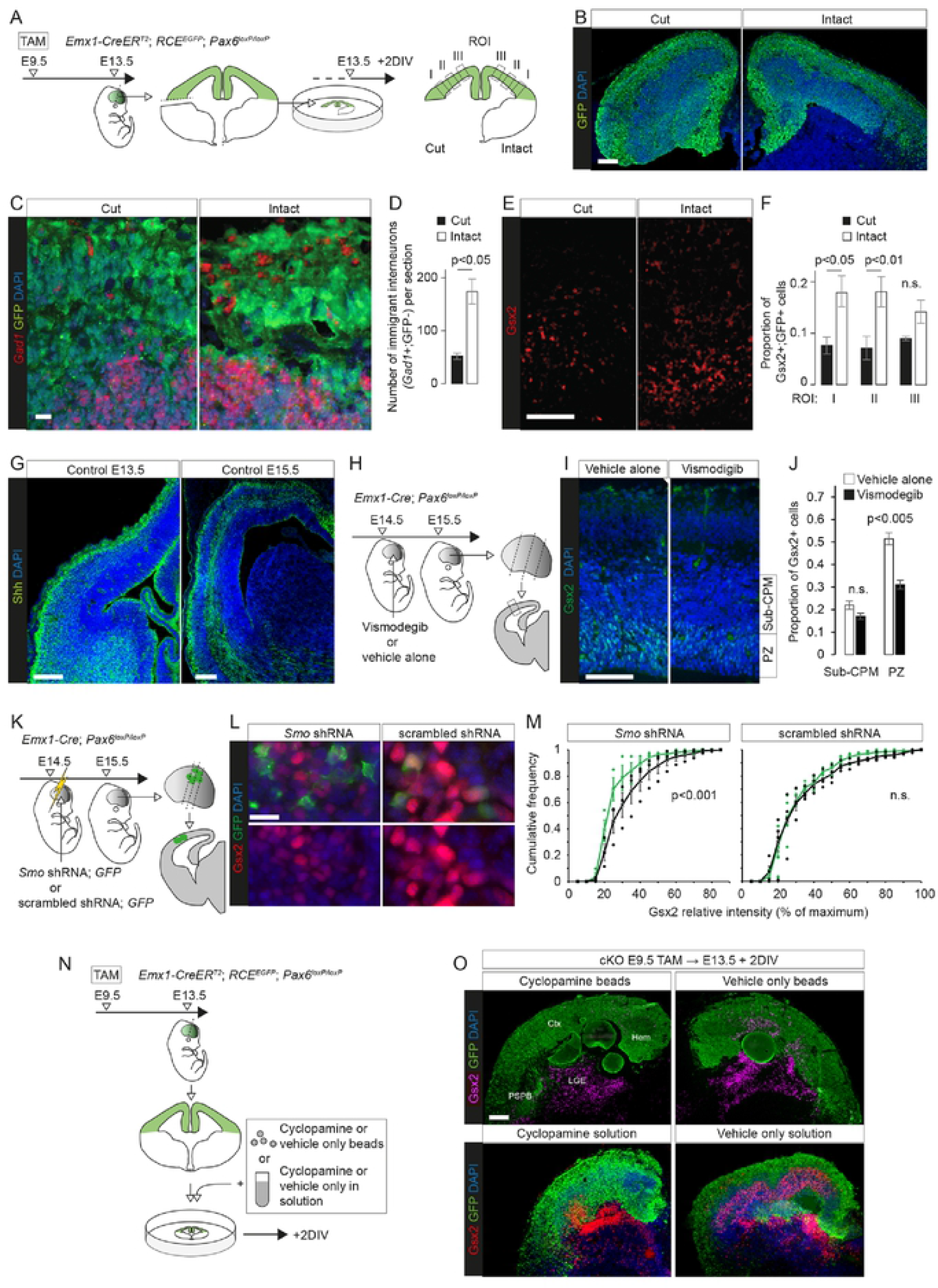
Extracellular signals promoted eGC production in *Pax6* cKOs cortex. **(A)** The experimental procedure for (B-F): tamoxifen (TAM) was given at E9.5 to generate *Pax6* cKOs, with Cre deleted cells expressing GFP; coronal slices were cultured on E13.5 with the ventral telencephalon removed on one side; after 2 days in vitro (DIV) sections from cultured slices were cut and processed. Gsx2+ GFP+ cells were counted in 3 regions of interest (ROI) on each side. **(B,C)** GFP immunoreactivity and in situ hybridizations for *Gad1* in sections prepared as in (A). Scale bars: 0.1mm and 0.01mm. **(D)** Average (±sem) numbers of immigrant *Gad1*+ interneurons (i.e. GFP non-expressing) per section were lower on the side lacking ventral telencephalon (n=3 independent cultures; Student’s paired t-test) (Source Data Table 5A). **(E)** Gsx2 immunoreactivity in sections prepared as in (A). Scale bar: 0.1mm. **(F)** Average (±sem) proportions of GFP+ cells that were Gsx2+ in each ROI in (A) (n=3 independent cultures; Student’s paired t-tests; n.s., not significant) (Source Data Table 5B). **(G)** Immunoreactvity for Shh in control telencephalic sections at E13.5 and E15.5 (see Fig. S10B for evidence of antibody specificity). Scale bar: 0.1mm. **(H)** The experimental procedure for (I,J): vismodegib or vehicle alone was injected into the ventricle of E14.5 *Pax6* cKO embryos made using *Emx1-Cre*; central regions of lateral cortex from coronal sections at three rostral-to-caudal levels were analysed at E15.5. **(I)** Gsx2 immunoreactivity in boxed region in (H). PZ: proliferative zone; sub-CPM: sub-cortical plate masses. Scale bar: 0.1mm. **(J)** Average (±sem) proportions of cells in the PZs and CPMs that were Gsx2+ (n=5 embryos from 3 litters given vehicle alone; n=6 embryos from 3 litters given vismodegib; Student’s t-tests) (Source Data Table 6A). **(K)** The experimental procedure for (L,M): constructs expressing *Smo* shRNA + GFP or scrambled shRNA + GFP were electroporated into the cortex of E14.5 *Pax6* cKO embryos made using *Emx1-Cre*; electroporated cells were analysed at E15.5 (as in Fig. S10D). **(L)** Gsx2 and GFP immunoreactivity in electroporated regions. Scale bar: 0.01mm. **(M)** Cumulative frequency distributions of the intensity of Gsx2 immunoreactivity in electroporated cells (GFP+; green) and surrounding randomly selected non-electroporated cells (GFP-; black) for the two constructs (see K and Fig. S10D) (n=3 embryos from 3 litters given *Smo* shRNA; n=4 embryos from 3 litters given scrambled shRNA; Kolmogorov–Smirnov tests) (Source Data Table 6B). **(N)** The experimental procedure for (O): tamoxifen (TAM) was given at E9.5 to generate *Pax6* cKOs and Cre deleted cells expressed GFP; coronal slices of telencephalon were cultured on E13.5; cyclopamine or vehicle alone were added either on beads or in solution (10µM); slices were cultured for 2 DIV. **(O)** Sections from cultured slices obtained as in (N) were immunoreacted for Gsx2 and GFP. Ctx: cortex; PSPB, pallial-subpallial boundary; LGE, lateral ganglionic eminence; Hem: cortical hem. Scale bar: 0.1mm.

This outcome suggested that the proportions of *Pax6* cKO cortical cells that deviated to develop as eGCs was influenced by extracellular factors.

### Misspecification of *Pax6* cKO cortical cells depended on their ability to respond to Shh

We then hypothesized that the signalling molecule, Shh, might be one factor contributing to the reprogramming of *Pax6*-deleted RGPs and their daughters. The embryonic cortex contains Shh from a variety of sources, including immigrating interneurons (Winkler et al., 2018; Komada et al., 2008; Baudoin et al., 2012) and cerebrospinal fluid (CSF) (Chau et al, 2015). Immunohistochemistry showed that Shh levels varied considerably with cortical location and that its distribution patterns were similar in control and *Pax6* cKO embryonic cortex of equivalent ages (Fig. 10G; S10C; evidence for antibody specificity is in Fig. S10B). Shh levels were higher laterally at E13.5 and increased across the cortex over the following two days (Fig. 10G).

To test the importance of endogenous Shh, we injected either an antagonist of the Shh receptor Smo (vismodegib; Morinello et al., 2015) into the lateral ventricle or electroporated a plasmid expressing both an shRNA against *Smo* and GFP into the cortex of *Emx1-Cre* induced *Pax6* cKO embryos and measured the effects on cortical Gsx2 expression (Fig. 10H-M). (We used *Emx1-Cre* rather than *Emx1-Cre^ERT2^* because we found it gave better survival rates following in utero surgery, while inducing a similar pattern of Gsx2 expression.) Vismodegib intraventricular injection significantly lowered by ∼40% the proportions of proliferative zone cells expressing Gsx2 compared to vehicle-only injection (Fig. 10H-J). Cells expressing *Smo* shRNA (GFP+) were on average significantly less immunoreactive for Gsx2 than a randomly selected sample of interspersed non-expressing (GFP-) cells, a difference that was lost when a control scrambled shRNA (GFP+) was used (Fig. 10K-M; for quantification method see Fig. S10D).

Further evidence came from adding the Shh pathway blocker cyclopamine, either in beads or in solution, to cultured *Pax6* cKO slices (Fig. 10N,O). This reduced their ectopic cortical expression of Gsx2. Interestingly, it had little if any effect on normal Gsx2 expression in the GEs, suggesting that their state of commitment was higher than that of the more labile eGC population.

Previous work had shown that the ability of embryonic telencephalic cells to express GE marker genes in response to the ventralizing morphogen Shh requires the transcription factor Foxg1, which we found was expressed by cortical RGPs and their daughters in both control and Pax6 cKO cortex (Fig. S3) (Danesin et al., 2009; Manuel et al., 2010). We postulated that Pax6 and Foxg1 have opposing actions (which might be direct, indirect or both) on specific aspects of cortical cells’ competence to respond to Shh, including their ability to activate ventral telencephalic marker genes, but not others (Fig. 11A). This idea was based on the following evidence from our RNAseq data and previous studies. First, we found that Pax6 removal caused little or no change in canonical readouts of Shh activity, namely *Ptch1* and *Gli1* mRNA expression levels (Goodrich et al, 1996; Marigo and Tabin, 1996; Lee et al, 1997; Wijgerde et al, 2002; Bai et al, 2002; Bai et al, 2004); only *Gli1* was significantly upregulated to a small extent (LFC=0.56) in caudal cortex at E13.5 (Table S1). Second, in *Pax6* cKO cortex, there were no abnormalities in the expression of mRNAs for Shh itself, the Shh receptor Smo or modulators of the Shh intracellular signal transduction pathways such as Kif7 and Sufu (Svard et al, 2006; Endoh-Yamagami et al. 2009; Liem et al., 2009) (Table S1). Third, previous work in *Foxg1^-/-^* telencephalon found that while cells failed to activate GE marker genes in response to Shh, they did activate *Ptch1* and *Gli1* normally (Manuel et al., 2010).

**Figure 11:**
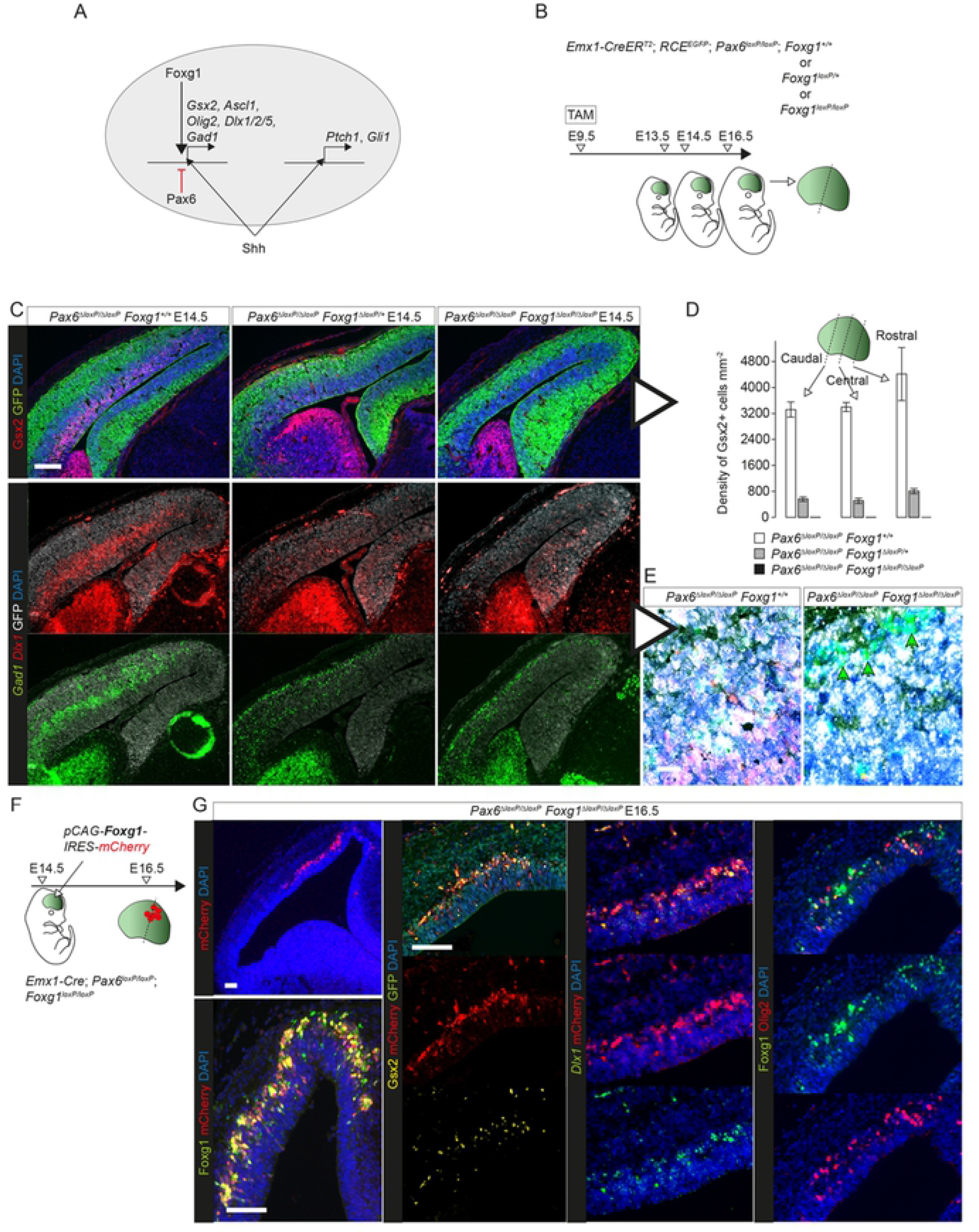
Foxg1 required cell autonomously for eGC production. **(A)** A hypothesis: Pax6 suppresses specifically Foxg1-permitted Shh-induced generation of eGCs without interfering with other effects of Shh pathway activation in these cells. **(B)** The experimental procedure for (C-E): tamoxifen (TAM) was administered at E9.5 to generate *Pax6* cKOs in which neither, one or both *Foxg1* allele(s) were also deleted (for alleles, see Fig. S11A); brains were analysed at E13.5, E14.5 or E16.5. **(C)** Immunoreactivity for Gsx2 and GFP and in situ hybridizations for *Gad1* and *Dlx1* in E14.5 cortex from *Pax6* cKO embryos in which neither, one or both *Foxg1* allele(s) were deleted by tamoxifen at E9.5. Scale bar: 0.1mm. **(D)** Densities of Gsx2+ cells in the lateral cortex of E14.5 embryos with the three genotypes in (C) (averages ± sem; n=3 embryos of each genotype, from 3 independent litters) (Source Data Table 6C). **(E)** High magnification images from (C): at least the majority of residual *Gad1*+ (green) cells in *Pax6 Foxg1* double KOs (arrows) were GFP-negative (i.e. not white) of subcortical origin (non-*Emx1*-lineage). Scale bar: 0.01mm. **(F)** The experimental procedure for (G): *Pax6 Foxg1* double cKO cortex made using *Emx1-Cre*, avoiding the need for tamoxifen; a construct expressing Foxg1 and mCherry was electroporated into the cortex on E14.5; coronal sections were analysed on E16.5. **(G)** Results of experiment in (F): expression and co-expression of Foxg1, Gsx2, Olig2, mCherry and GFP protein and *Dlx1* mRNA in coronal sections. Scale bars: 0.1mm.

To test the prediction that the upregulation of GE markers in *Pax6* cKOs would be reversed by Foxg1 removal, we used tamoxifen^E9.5^-induced *Emx1-Cre^ERT2^* to delete both copies of *Pax6* together with both, one or neither copies of *Foxg1* from embryonic cortical cells (Fig. 11B; Fig. S11A). Deletion of both copies of *Foxg1* resulted in the loss of *Foxg1* mRNA from almost all cortical cells by E13.5 (Fig. S11B,C); the few remaining undeleted cells formed small clones expressing both Foxg1 and Pax6 (arrows in Fig. S11C). Deletion of one copy of *Foxg1* appeared to lower its cortical mRNA and protein levels (Fig. S11C).

Deletion of both copies of *Pax6* together with one copy of *Foxg1* reduced the numbers of cortical cells expressing Gsx2, *Dlx1* and *Gad1* at E14.5 (Fig. 11C,D). Deletion of both copies of both *Pax6* and *Foxg1* abolished cortical expression of Gsx2 and *Dlx1* (Fig. 11C,D) and left only *Gad1*+ cells that were GFP-negative and presumably had originated subcortically (arrows in Fig. 11E). Deletion of one or both copies of *Foxg1* also reduced Ascl1 expression in *Pax6* cKOs, with a greater effect in lateral than in medial cortex (Fig. S11D). The reduction of the proportions of progenitor layer cells that were Eomes+ in E16.5 *Pax6* cKO cortex was reversed by co-deletion of one or both copies of *Foxg1* (Fig. S11E,F).

To confirm that these actions of Foxg1 were cell autonomous, we electroporated *Pax6^-/-^*; *Foxg1^-/-^* double conditional knock-out cortex with a plasmid construct that resulted in the expression of mCherry and the re-expression of Foxg1 by small groups of cells (Fig. 11F,G). Two days after electroporation, an average of 33.2% (±7.4 s.d.; n=3 embryos) of electroporated cells re-expressed Gsx2 (Source Data Table 6E) and many were re-expressing *Dlx1* and Olig2 (Fig. 11G). Electroporated cells were much less likely than their non-electroporated neighbours to express Eomes (Fig. S11G), in line with their re-direction to an eGC-like fate.

In summary, these findings all suggested that Pax6 limits the competence of cortical cells to respond to Shh in their local environment by preventing them from adopting Foxg1-dependent developmental trajectories towards GE-like fates.

### *Pax6* deletion increases cortical cells’ sensitivity to Shh pathway activation

We then compared the sensitivity of control and *Pax6* cKO cortical cells, in terms of their ability to express GE/ eGC marker genes in response to Shh pathway activation. We dissociated E13.5 control or *Pax6* cKO cortex (tamoxifen^E9.5^) carrying the GFP Cre-reporter, cultured the cells for 48h in the presence of Shh signalling agonist (SAG; Chen et al., 2002) or vehicle alone (Fig. 12A) and quantified numbers of GFP+ cells expressing Gsx2, Olig2 or *Dlx1* (Fig. 12B; quantification method in Fig. S12A). In common with previous studies, we used SAG concentrations in the nM range (Heine et al., 2011; Chen et al., 2002), which existing evidence suggested would likely have covered the levels of pathway activation experienced by telencephalic cells in vivo (Radonjic et al., 2016; Chau et al., 2015).

**Figure 12:**
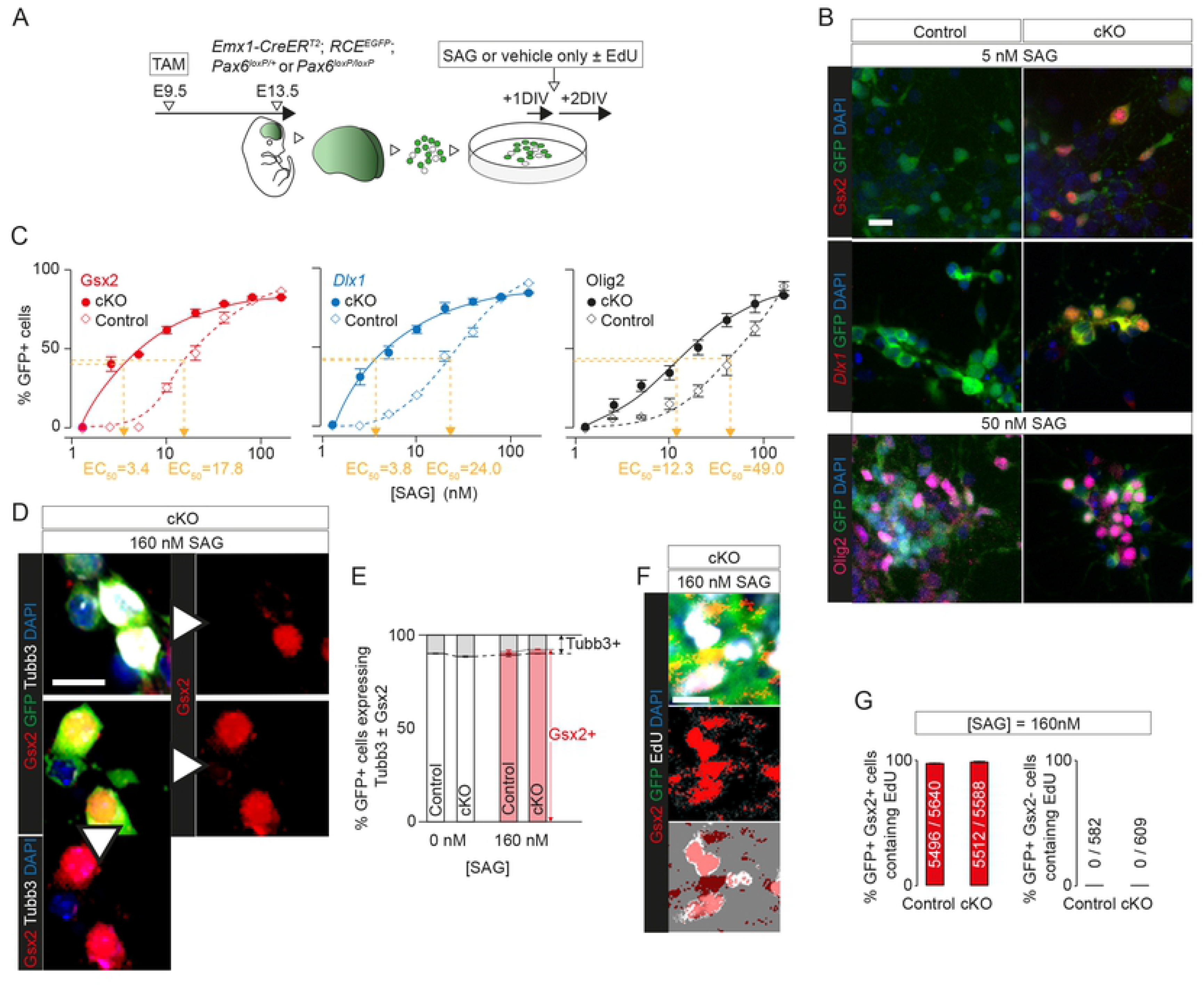
*Pax6* deletion affected concentration-response of cortical cells to Shh pathway activation. **(A)** The experimental procedure for (B-G): tamoxifen (TAM) was given at E9.5 to generate *Pax6* cKO and control embryos, with Cre activation revealed by GFP expression; E13.5 cortex was dissociated, cells were treated with SAG or vehicle alone, and with EdU in some cases, after 1 day in vitro (DIV), and were analysed after a further 2DIV. **(B)** Examples of labelling of E13.5 control and *Pax6* cKO cells grown in dissociated culture with 5nM or 50nM SAG. Labelling was for DAPI and GFP with Gsx2, *Dlx1* or Olig2. Scale bar: 0.01mm. **(C)** Graphs of concentration-responses to SAG (measured as percentages of GFP+ cells expressing Gsx2, *Dlx1* or Olig2). Data are averages (±sem; n=3 independent experiments each), with EC_50_s for each response curve. Two-way analyses of variance were conducted. For Gsx2: significant effects of genotype (f(1,32)=798.9, p<0.001) and SAG concentration (f(7,32)=1138, p<0.001) and significant interaction effect (f(7,32)=123.5, p<0.001). For *Dlx1*: significant effects of genotype (f(1,32)=763.6, p<0.001) and SAG concentration (f(7,32)=1011, p<0.001) and significant interaction effect (f(7,32)=91.90, p<0.001). For Olig2: significant effects of genotype (f(1,32)=177.4, p<0.001) and SAG concentration (f(7,32)=415.1, p<0.001) and significant interaction effect (f(1,32)=15.88, p<0.001) (Source Data Table 7A-C). **(D)** Examples of labelling of E13.5 control and *Pax6* cKO cells grown in dissociated culture with 160nM SAG. Labelling was for DAPI, GFP, Gsx2 and Tubb3. Examples include GFP+ cells that were: Gsx2+, Tubb3-; Gsx2+, Tubb3+; Gsx2-, Tubb3+. Scale bar: 0.01mm. **(E)** Average percentages (±sem; n=4 independent experiments each) of GFP+ control or *Pax6* cKO cells with or without 160nM SAG that expressed Tubb3, Gsx2 or both (Source Data Table 7D). **(F)** Examples of labelling of E13.5 control and *Pax6* cKO cells grown in dissociated culture with 160nM SAG. Labelling was for DAPI, GFP and EdU. Examples include GFP+ cells that were: Gsx2+, EdU+; Gsx2+, EdU-. Scale bar: 0.01mm. **(G)** Average percentages (±sem; n=4 independent experiments each) of GFP+ cells that contained EdU among the Gsx2+ and Gsx2-populations in cultures from control and *Pax6* cKO cortex treated with SAG. Total numbers, across all cultures, of GFP+ cells that contained EdU over total numbers of GFP+ cells are stated for each condition (Source Data Table 7E).

We found that no cells cultured with the lowest doses of SAG expressed Gsx2, *Dlx1* or Olig2, despite the fact that all three genes would have been expressed by significant numbers of cells in E13.5 *Pax6* cKO cortex. The likely explanation for this, in line with suggestions made above, was that they were in a labile state requiring continual activation of their Shh signalling pathways to maintain their aberrant identity, and the signals they were receiving in vivo would been have dissipated by dissociation.

We found that control cells responded in a concentration-dependent manner to addition of SAG, in agreement with previous work showing that non-physiological elevation of Shh signalling in normal embryonic cortex can activate the expression of ventral telencephalic marker genes (Gaiano et al., 1999; Corbin et al., 2000; Rallu et al., 2002; Quinn et al., 2009; Shikata et al., 2011; Yabut et al., 2015). However, *Pax6* cKO cells were significantly more sensitive to SAG than control cells (Fig. 12C). The concentration-response functions for Gsx2 and *Dlx1* were relatively similar, reflecting the close association between their expression patterns in vivo, but differed from those for Olig2, which showed a different pattern of activation in vivo (Fig. 6I). EC_50_s for Olig2 were ∼2-3 times higher, with cells less likely to express Olig2 than Gsx2 or *Dlx1* in response to low/ intermediate levels of SAG (Fig. 12C). Since our in vivo findings had shown that early Olig2 activation was more widespread than early Gsx2 and *Dlx1* activation in *Pax6* cKO cortex (Fig. 6A,B,I), this suggested that factors additional to Shh activation were required to explain the difference between the in vivo patterns of activation (see the next section).

The concentration-response functions for Gsx2 and *Dlx1* appeared to plateau with ∼85% of GFP+ cells expressing the markers, suggesting that ∼15% of E13.5 cortical cells were not competent to respond to Shh activation. Since this was similar to the percentage of differentiating glutamatergic neurons in E13.5 control or *Pax6* cKO cortex (Fig. 2D), we tested whether the incompetent cells were those that were most highly differentiated. In one set of experiments, we identified differentiating neurons by their expression of Tubb3 (Fig. 12D,E). We found that 9.9-11.5% of control and *Pax6* cKO GFP+ cells expressed Tubb3 whether SAG was added or not and that all GFP+ Gsx2-negative cells in SAG-treated cultures were Tubb3+, with only 2.0-2.3% of GFP+ cells expressing both Gsx2 and Tubb3. In another set of experiments, we identified cells that had divided in culture by adding the thymidine analogue EdU to the culture medium (Fig. 12A,F,G). We found that EdU was incorporated by most of the GFP+ cells that had activated Gsx2 in response to SAG, but by none of those that remained Gsx2-negative. We concluded that the cells that were the most highly differentiated were the least susceptible to the effects of SAG.

### Bmps contributed to regional differences in ectopic gene activation

We next considered whether Pax6 is also involved in regulating the responses of developing cortical cells to other morphogens. Previous research has shown that the embryonic dorsal telencephalic midline is a rich source of Bmps, including Bmps 4-7 (Furuta et al., 1997), and that Bmps can inhibit the expression of genes involved in the specification of GABAergic neurons (Gulacsi and Lillien, 2003). Our evidence indicated that many aspects of cortical Bmp signalling remained close to normal in *Pax6* cKO cortex. Our RNAseq data showed that Pax6 removal had no detectable effects on the expression of mRNAs for any of the Bmps and identified only two canonical Bmp signalling pathway genes with significantly altered expression levels in *Pax6*-deleted cortex (both only at E13.5: *Bmpr1b* LFC=-0.76 rostrally and -0.64 caudally; *Smad3* LFC=-0.35 rostrally and -0.28 caudally) (Table S1).

Second, phospho-Smad1/5/9 immunoreactivity, whose levels correlate positively with Bmp activity and, therefore, tend to be higher in medial than in lateral embryonic cortex (Cheng et al., 2006; Hu et al., 2008; Doan et al., 2012; Lehtinen et al., 2011), showed a similar pattern in control and *Pax6* cKO embryos (Fig. S13A).

We tested the effects of Bmp4 on the expression of *Gsx2* and *Prdm13*, whose spatial expression patterns were altered in different ways by *Pax6* deletion. We added increasing doses of Bmp4 to cultured E13.5 control and *Pax6* cKO cortical slices and measured gene expression levels with qRT-PCR and visualised expression patterns in sections (Fig. 13A). We found that addition of Bmp4 to *Pax6* cKO cortex suppressed *Gsx2* mRNA levels and caused loss of Gsx2 immunoreactivity in sections (Fig. 13B,C).

**Figure 13:**
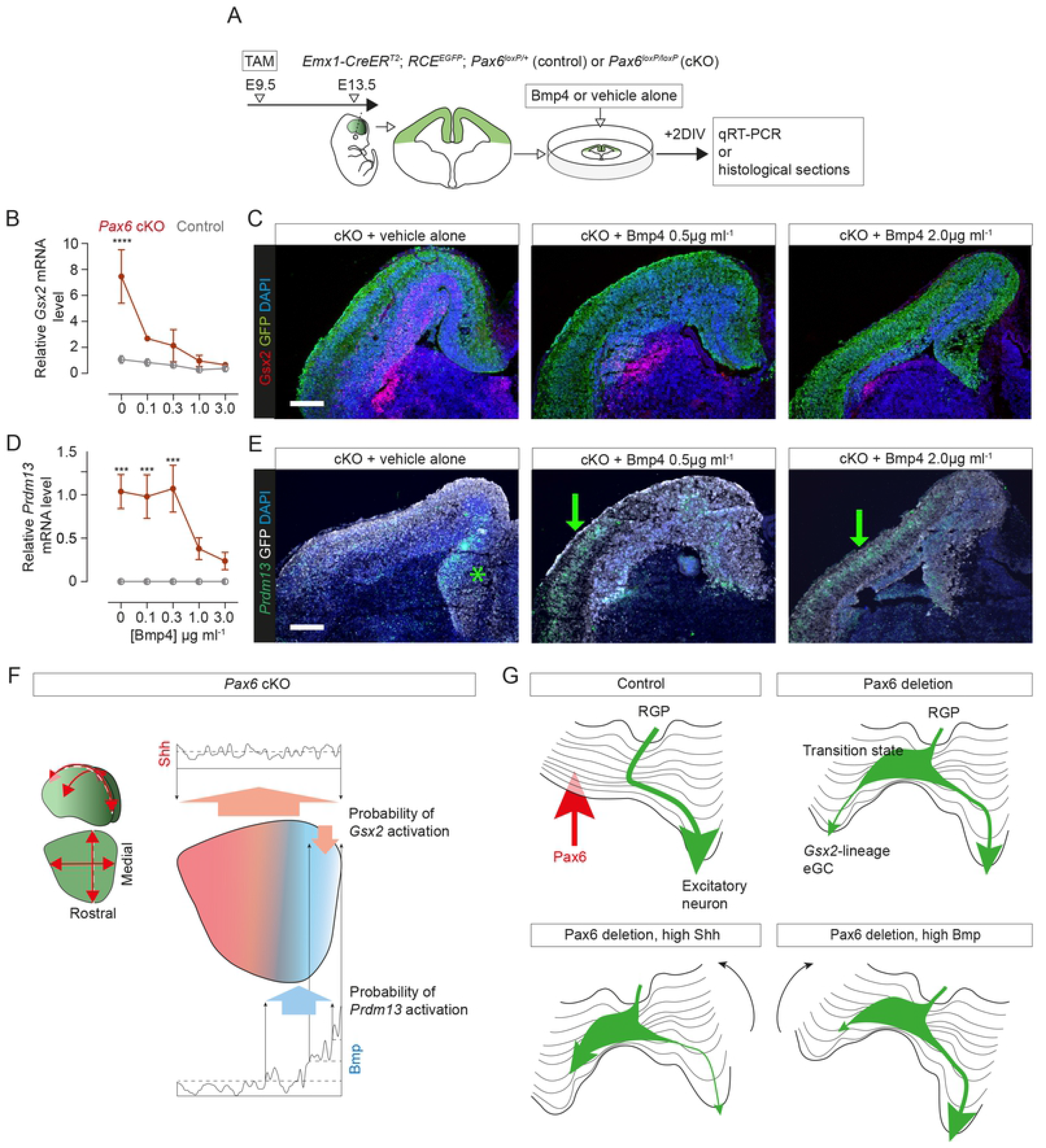
Morphogen regulation of ectopic gene expression following Pax6 loss. **(A)** The experimental procedure for (B-E): tamoxifen (TAM) was administered at E9.5 to delete either one (control) or both (cKO) *Pax6* allele(s), with Cre deleted cells expressing GFP; coronal slices were cultured on E13.5 with Bmp4 or vehicle alone for 2 days in vitro (DIV); slices were analysed using qRT-PCR or sectioned. **(B)** Concentration-response measured using qRT-PCR: *Gsx2* levels (averages ±sem; values were relative to the average level in control cortex treated with 0 Bmp4) in control and *Pax6* cKO slices with increasing concentrations of Bmp4 (n=3 independent cultures at each concentration). Two-way ANOVA showed significant effects of genotype (p<0.001), of Bmp4 concentration (p<0.005) and an interaction effect (p<0.01). Differences between genotypes at each Bmp4 concentration were tested with Bonferroni’s method for comparison of means (****, p<0.001) (Source Data Table 8). **(C)** Immunoreactivity for Gsx2 and GFP in telencephalic slices from *Pax6* cKOs cultured with vehicle alone or Bmp4. Scale bar: 0.1mm. **(D)** Concentration-response measured using qRT-PCR: *Prdm13* levels (averages ±sem; values are relative to the average level in *Pax6* cKO cortex treated with 0 Bmp4) in control and *Pax6* cKO slices with increasing concentrations of Bmp4 (n=3 independent cultures at each concentration). Two-way ANOVA showed significant effects of genotype (p<0.005), of Bmp4 concentration (p<0.05) and an interaction effect (p<0.05). Differences between genotypes at each Bmp4 concentration were tested with Bonferroni’s method for comparison of means (***, p<0.005) (Source Data Table 8). **(E)** In situ hybridizations for *Prdm13* and immunoreactivity for GFP in telencephalic slices from *Pax6* cKOs cultured with vehicle alone or Bmp4. Green arrows indicate *Prdm13* expression in lateral cortex. Scale bar: 0.1mm. **(F)** A hypothesis of how Shh and Bmp4 might combine to generate the observed spatial patterns of *Gsx2* and *Prdm13* expression in the embryonic cortex after Pax6 deletion. Deletion might increase the probability of *Gsx2* being activated in cells exposed to physiological levels of Shh above a threshold (broken line). In medial cortex, exposure the levels of Bmp above a threshold (central broken line) might reduce the probability of Gsx2 activation. Cells exposed to intermediate levels of Bmp (between upper and lower broken lines) might have an increased probability of expressing *Prdm13*. **(G)** Waddington’s epigenetic landscape analogy, used to illustrate our main conclusions. A saddle-node bifurcation illustrates Pax6’s normal action, closing a valley on the left (RGP: radial glial progenitor). Pax6 deletion opens this valley, creating a subcritical pitchfork bifurcation where cells emerging from the transition state can enter either of two valleys (eGC: ectopic GABAergic cell). Increasing exposure to Shh tilts the landscape to the left making it more likely that the cell will enter the open valley on the left; increasing exposure to Bmp has the opposite effect.

The effects of Bmp4 on *Prdm13* expression were more complex (Fig. 13D,E). Whereas addition of Bmp4 at concentrations <1µg mL^-1^ had no detectable effect on overall levels of *Prdm13* measured with qRT-PCR, and higher concentrations suppressed overall expression (Fig. 13D), in situ hybridizations in sections revealed that addition of Bmp4 actually increased *Prdm13* expression in lateral cortex (Fig. 13E, green arrows). A possible reason why this was not reflected in the *overall* levels of *Prdm13* mRNA was that it appeared to be offset by decreased expression in medial cortex (Fig. 13E, asterisk). It seemed possible that *Prdm13* responded biphasically to Bmp activation in *Pax6* cKO cortex, such that: i) *Prdm13* was activated in the range of Bmp activation levels that existed endogenously in medial cortex or were achieved in lateral cortex after exogenous application of Bmp4; ii) *Prdm13* expression was suppressed at the relatively higher Bmp activation levels that were achieved medially when endogenous Bmp activation was supplemented by exogenous Bmp4. Our findings indicated that cells in both medial and lateral *Pax6* cKO cortex were competent to express *Prdm13* and whether they did so depended on them receiving the requisite signals.

Figure 13F outlines how Shh and Bmp4 might combine to generate the spatial expression patterns of *Gsx2* and *Prdm13* in *Pax6* cKOs (Figs. 5,6). In this model, *Gsx2* activation in cells exposed to supra-threshold levels of Shh is counteracted in medial cortex by relatively high levels of Bmp signalling, preventing Gsx2 expression in this region. *Prdm13* is activated by intermediate levels of Bmp signalling but suppressed by the highest levels, which might explain the lowering of *Prdm13* expression very close to the dorsal midline (Fig. 5F,G).

We found that Bmp4 did not suppress the expression of Olig2 in *Pax6* cKO cortex (Fig. S13B). The selective suppression of Gsx2 but not Olig2 by Bmps suggests a possible explanation for results described above, which showed that Pax6 deletion induced more widespread activation of Olig2 than of Gsx2 in vivo (Fig. 6A,B,I) even though Shh activation had a greater effect on the activation of the latter (Fig. 12C). Bmps might be the additional factors required to reconcile these findings.

## Conclusion

Figure 13G illustrates our main findings using Waddington’s epigenetic landscape in which the developmental trajectory of a cell is represented as a ball rolling downhill through valleys created by the actions of that cell’s genes. Pax6’s actions ensure that, under physiological conditions, cortex-born cells progress unerringly towards their excitatory neuronal fate. If Pax6 is deleted, alternative trajectories become available (at what are known as pitchfork bifurcations: Huang et al., 2007; Moris et al., 2016). Morphogens such as Shh and Bmps, whose concentrations vary dynamically and regionally, have the effect of tilting each cell’s landscape to a variable degree, thereby influencing which alternative is likely to be adopted (Ferrell, 2012). Pitchfork bifurcations are associated with unstable and reversible transition states in which the cell shows a mixed identity between the original and the destination states (Moris et al., 2016). We envisage that the aPs we identified by scRNAseq in *Pax6* cKO cortex are examples of cells in this state.

## DISCUSSION

### Pax6 affects the competence of cortical cells to respond to signalling molecules

Regionally expressed transcription factors such as Pax6 can contribute to the development of cellular diversity in several ways: (i) by regulating cells’ competence to respond to signalling molecules around them; (ii) by controlling their production of intercellular signalling molecules; (iii) by acting internally to influence cellular development without affecting intercellular signalling. Our present study has highlighted the overriding importance of Pax6 in the first of these mechanisms during cortical neurogenesis, although it is probably involved to some extent in all three.

Pax6 is expressed in many cell types in structures including the eye, brain, spinal cord and pancreas (Walther and Gruss, 1991; Turque et al., 1994). Its regulation of cellular competence during cerebral cortical neurogenesis is mirrored by some of its actions in other tissues. For example: Pax6 is involved in establishing the competence of different sets of murine thalamic cells to produce either glutamatergic or GABAergic neurons (Robertshaw et al., 2013); PAX6 restricts the competence of neuroepithelium derived from human embryonic stem cells to express markers of GABAergic progenitors in response to SHH application (Chi et al., 2017); Pax6 regulates the competence of cells to respond to inductive signals during vertebrate and invertebrate eye development (Collinson et al., 2000; Luan et al., 2014). It is important to stress, however, that the nature of Pax6’s effects vary considerably in different contexts: for example, whereas it limits cells’ competence to express genes such as *Gsx2* during cerebral corticogenesis, it has an opposite effect on *Gsx2* expression in diencephalic cells (Caballero et al., 2014; Quintana-Urzainqui et al., 2018). This and other context-dependent differences likely arise at least in part from differences in its combinatorial expression with other transcription factors, such as Foxg1 in the telencephalon (Quintana-Urzainqui et al., 2018) or Irx3 in the diencephalon (Robertshaw et al., 2013).

Regarding the second possibility listed above -that Pax6 regulates the production of intercellular signalling molecules that affect how other cells develop - our RNAseq data identified very few changes in the expression of genes encoding signalling molecules. These did, however, include a change in *Ffg15* expression, which was upregulated in rostral cortex following *Pax6* deletion. This is potentially interesting because experimentally-induced upregulation of *Ffg15* in embryonic cortex can facilitate the activation of ventral telencephalic genes (Yabut et al., 2020). Although previous work on the embryonic cortex of mouse chimeras containing mixtures of wild-type and *Pax6*-null cells found no evidence that wild-type cells were affected by abnormal signalling from *Pax6*-null cells (Quinn et al., 2007), it remains possible that altered signal production by *Pax6*-deleted cells affects other *Pax6*-deleted cells, due to their altered competence.

Regarding the third possibility listed above, Pax6’s regulation of genes not directly involved in intercellular signalling almost certainly contributes to some of its cellular actions. For example, we have argued previously that Pax6 limits cortical progenitor cell cycle rates through mechanisms that include direct repression of *Cdk6* (Mi et al., 2013). Nevertheless, such explanations are probably incomplete. For example, Shh is a mitogen (Kalyani et al., 1998; Wechsler-Reya and Scott, 1999; Groves et al., 2020) and the highly proliferative nature of many eGCs identified in the present study might be caused in part by cortical cells’ having enhanced responses to Shh that include effects on their proliferation.

Our overall conclusion is that, while Pax6’s dominant role during cortical neurogenesis is to limit the developmental potential of cortical cells to respond to signalling molecules that are carrying out other functions around them, it is very likely to act in other ways too.

### Pax6 raises cortical cellular thresholds for responses to morphogens such as Shh

In developing multicellular organisms, the reproducible production of distinct specialized cell types in specific locations involves cells acquiring information about their relative positions through interactions with their extracellular environment. For example, one iconic model proposes that cells in a contiguous population, or field, acquire positional information from their levels of exposure to substances distributed in concentration gradients across the field (Wolpert, 1969). It is thought that such mechanisms specify the two major domains of the rostral neural plate, a ventral domain producing the basal ganglia and a dorsal domain producing the cerebral cortex. The morphogen Shh is one example of a substance whose concentration varies dorsoventrally as these two domains emerge, with its high ventral levels contributing to the specification of ventral fates (Marti et al., 1995; Matise and Wang, 2011). As the rostral neural tube expands and complexifies, however, so do the distributions of morphogens that affect its subsequent development. For example, the closure of the neural tube to create the nervous system’s ventricular system gives numerous morphogens, including Shh, widespread access to developing telencephalic cells via the cerebrospinal fluid (Chau et al., 2015) and interneurons migrating into the cortex produce numerous morphogens, including Shh (Komada et al., 2008; Baudoin et al., 2012; Voronova et al., 2017; Winkler et al., 2018). Previous work has suggested that Shh levels in embryonic cortex are high enough that they could, potentially, induce the ectopic expression of ventral marker genes (Yabut et al, 2020). These observations bring into sharp focus the importance of mechanisms regulating not only the distributions of morphogenetic substances but also the responses of cells to them.

A classic proposition is that cells have intrinsic thresholds determining how aspects of their development are influenced by surrounding morphogens (Wolpert, 1969). Our work suggests that Pax6 raises some of these intrinsic thresholds in embryonic cortical cells, preventing them from activating erroneous programs of gene expression in response to physiological levels of Shh, Bmps and possibly other morphogens around them (Dahmane et al., 2001; Viti et al., 2003; Komada et al, 2008; Xu et al., 2010; Baudoin et al., 2012; Winkler et al., 2018; Yabut and Pleasure, 2018; Yabut et al, 2020; Zhang et al., 2020; Delmotte et al., 2020). The reasons for thinking in terms of threshold adjustment is that Pax6 does not produce an absolute block under all circumstances to the possibility of morphogen-induced ectopic gene expression. Previous work has shown that non-physiological elevation of Shh signalling in Pax6-expressing embryonic cortex can activate the ectopic expression of ventral telencephalic marker genes (Gaiano et al., 1999; Corbin et al., 2000; Rallu et al., 2002; Quinn et al., 2009; Shikata et al., 2011; Yabut et al., 2015). Our in vitro data showed that Shh can induce Pax6-expressing cortical cells to express ventral telencephalic markers in a concentration-dependent manner, but *Pax6* cKO cells respond with much greater sensitivity.

Pax6 acts selectively on the various cellular process affected by morphogen signalling. It does not prevent morphogens such as Shh and Bmps having important physiological functions in processes including cortical cell proliferation, migration and neuronal morphogenesis (Komada et al, 2008; Baudoin et al., 2012; Saxena et al, 2018; Yabut and Pleasure, 2018; Hou et al., 2021). This selectivity might be a consequence of Pax6 acting around the point where the intracellular signal transduction pathways target the genome; for example, it might reduce the ability of molecules acting late in these pathways, such as Gli and Smad transcription factors (Matise and Wang, 2011; Hegarty et al., 2013; Yabut and Pleasure, 2018; Groves et al., 2020), to bind some of their potential genomic target sites. At least some such effects might involve indirect actions, via effects of Pax6 on the expression of other transcription factors. For example, Ascl1 is a possible intermediate. We found that its upregulation in *Pax6* cKO cortex is widespread and is associated with multiple lineages. Previous work has shown (i) that forced expression of Ascl1 in embryonic cortex activates the ectopic expression of ventral markers such as *Dlx1* and *Gad1* in some cortical cells (Fode et al., 2000), (ii) that Ascl1 promotes chromatin accessibility during neurogenesis and (iii) that *Olig2* is one of Ascl1’s direct targets (Raposo et al., 2015).

A remarkable feature of the mechanisms that regulate the development of multicellular organisms is the extent to which the same morphogens are reused in multiple diverse mechanisms as the embryo grows. It seems highly likely that adjustments to how cells at different locations and ages respond to the same morphogens would have been instrumental in allowing the diversification of morphogen function during evolution. The evolution of mechanisms limiting cellular competence, such as those described here, would have allowed a morphogen to acquire new functions by minimizing the risk of the beneficial consequences being offset by its pre-existing potential to induce changes that would be undesirable in the new context.

### Pax6’s effects on Shh signalling might control the timing of cortical excitatory neuron specification

Previous studies have shown that Shh can repress *Pax6* expression in diverse tissues including the telencephalon (Macdonald et al., 1995; Ericson et al., 1997; Rallu et al., 2002; Danjo et al., 2011; Robertshaw et al., 2013; Yabut et al., 2020). The fact that Shh levels are highest in ventral telencephalon from the earliest stages of its development is likely to be a major factor establishing the ventral-low versus dorsal-high difference in *Pax6*’s expression levels (Echelard et al., 1993; Shimamura et al., 1995; Gunhaga et al., 2000; Rallu et al., 2002b). Our findings indicate that, as dorsal telencephalic development progresses, its high levels of Pax6 repress the potential actions of morphogens including Shh itself, thereby opening a time-window for the unhindered production of a normal complement of cortical excitatory neurons.

When this time-window opens, Pax6 is normally expressed by a large proportion of cortical cells, with highest levels in RGPs (Caric et al., 1997; Englund et al., 2005; Wong et al., 2015). For example, our scRNAseq data showed that more than 70% of cells express *Pax6* at E13.5. As the cortex ages, a progressively smaller proportion of cells express *Pax6* (∼50% at E14.5 in our scRNAseq data) as non-RGP populations, including non-RGP progenitors such as IPs, expand. This decline coincides with the normal cortical activation from ∼E16.5 onwards of *Gsx2* and *Olig2* in a small proportion of SVZ cells (Zhang et al., 2020). These cells are thought to be tri-potential IPs that generate interneurons for the olfactory bulb, cortical oligodendrocytes and astrocytes. Their generation requires Shh and they and their daughters express many of the same genes as those activated earlier in Pax6-deleted cortex. It seems plausible that, in normal corticogenesis, the waning influence of Pax6 in late progenitor populations allows the time-window for the focussed production of cortical excitatory neurons to close.

### Pax6 loss generates cortical cells with varying degrees of similarity to normal cell types

Previous studies have shown that even if cortical progenitors lose the ability to produce functional Pax6, whether due to constitutive or conditional mutation, they can still generate many cells that migrate into the CP to form layers containing neurons with excitatory morphologies and connections resembling those in normal cortex (Caric et al., 1997; Tuoc et al., 2009). In the present study we confirmed and extended these previous conclusions: the transcriptional profiles of *Pax6*-deleted cells that were differentiating into excitatory neurons and the electrophysiological properties of *Pax6*-deleted CP neurons were indistinguishable from those of control cells. Overall, we found no evidence that Pax6 is required to instruct the specification of cortical excitatory neurons from cortical progenitors.

Our analyses indicated that some populations of cells that are present in normal cortex expanded following Pax6 loss. This applied to cells in the dorsomedial part of the embryonic cortex. The expansion of these populations might have been due, at least in part, to an increased sensitivity to Bmps, which are present at higher levels dorsomedially and are involved in the specification of dorsomedial cell identities during normal development (Furuta et al., 1997; Gulacsi and Lillien, 2003; Cheng et al., 2006; Caronia et al., 2010; Doan et al., 2012).

Previous reports have suggested that *Pax6*-deleted cortical progenitors generate cells with a dorsal LGE (dLGE) identity that go on to produce immature olfactory bulb interneurons, as do dLGE cells in normal telencephalon (Stenman et al., 2003; Kroll and O’Leary, 2005; Guo et al., 2019). While some evidence supports this interpretation, we suggest that it should be treated with caution. Most tellingly, we found that many of these cells do not respond to Gsx2 loss in the same way as normal dLGE cells. This difference might reflect intrinsic differences with normal dLGE cells and/ or differences in their extracellular environments. Similarly, intrinsic and/ or environmental factors might contribute to the accumulation of abnormal cell types beneath the CP and their subsequent death. Overall, it seems unwise at present to assume that these abnormal cell types are necessarily representative of cell types present in normal telencephalon.

## METHODS

### Animals

All experimental procedures involving mice were regulated by the University of Edinburgh Animal Welfare and Ethical Review Body in accordance with the UK Animals (Scientific Procedures) Act 1986. All the alleles used and their genotyping have been described before: *Btg2-GFP* (Haubensak et al., 2004); *CAGG-Cre^ERTM^* (Hayashi and McMahon, 2002); *Emx1-Cre* (Gorski et al., 2002), *Emx1-Cre^ERT2^* (Kessaris et al., 2006), *Foxg1^loxP^* (Miyoshi and Fishell, 2012), *Gsx2-Cre* (Kessaris et al., 2006), *Pax6^loxP^* (Simpson et al., 2009), *Pax6^Sey^* (Hill et al., 1991), *Gsx2^loxP^* (Waclaw et al., 2009); *RCE:loxP* (*R26R CAG-boosted enhanced green fluorescence protein (EGFP)* Cre reporter) (Miyoshi et al., 2010); *Shh^-^* (Chiang et al., 1996). Activation of CreERT2 or CreERTM was achieved by giving pregnant females 10mg of tamoxifen by oral gavage.

### Quantitative real-time polymerase chain reaction (qRT-PCR)

Total RNA was extracted using either the RNeasy Plus Mini kit (Qiagen) for samples taken directly from cortex or the RNeasy Micro kit (Qiagen) for cultured cortical slices and cDNA was synthesized with a Superscript reverse transcriptase reaction (Thermo Fisher Scientific).

*Pax6*: qRT-PCR was performed using a DNA Engine Opticon Continuous Fluorescence Detector (MJ Research) and a Quantitect SYBR Green PCR kit (Qiagen). We used the following primer pairs. *Pax6*: 5’-TATTACGAGACTGGCTCCAT-3’ and 5’-TTGATGACACACTGGGTATG-3’; *Gapdh*: 5’-GGGTGTGAACCACGAGAAAT-3’ and 5’-CCTTCCACAATGCCAAAGTT-3’. We calculated the relative abundances of *Pax6* and *Gapdh* transcripts for each sample. For each biological replicate, we ran three technical replicates.

*Crabp1, Gsx2, Olig2, Prdm13*: qRT-PCR was performed using the Applied Biosystems StepOnePlus RT-PCR machine (Thermo Fisher Scientific) and a TaqMan Gene Expression Assay (Thermo Fisher Scientific) containing a TaqMan probe with a fluorescein amidite dye label on the 5’ end and a minor groove binder and non-fluorescent quencher on the 3’ end. TaqMan probes used in this experiment, designed and supplied by Thermo Fisher Scientific, were as follows. *Crabp1*: Mm00442775_g1; *Gsx2*: Mm00446650_m1; *Olig2*: Mm01210556_m1; *Prdm13*: Mm01220811_m1. For each sample, we ran three technical replicates with no-template control. Target gene expression was calculated as fold change relative to gene expression in the vehicle alone condition.

### Western blots

Proteins extracted from homogenized and lysed cortical tissue were resolved on denaturing gels. Primary antibodies against Pax6 (1:500; rabbit; Covance) and β-actin (1:2000; rabbit; Abcam) were detected with Alexa-coupled secondary antibodies and blots were quantified using the LI-COR scanning system (LI-COR Biosciences). The intensity of each Pax6 band was normalized to that of its corresponding β-actin band.

### Immunostaining and in situ hybridization on tissue sections

Pregnant mice were killed by cervical dislocation and embryonic brains were dissected into 4% paraformaldehyde (PFA); postnatal mice were perfused transcardially with 4% PFA; cultured cortical slices were placed in 4% PFA. Samples were fixed overnight at 4°C and cryosections were cut usually at 10μm or at 2μm in some cases.

### Immunostaining

Primary antibodies used in this study were as follows. Anti-Ascl1 (mouse; BD Bioscience); anti-Bromodeoxyuridine (BrdU) (mouse; clone B44, BD Bioscience); anti-Caspase 3 (rabbit; Millipore); anti-Crabp1 (rabbit; Cell Signalling); anti-Eomes (rabbit; Abcam); anti-Foxg1 (mouse; kindly provided by Steven Pollard, University of Edinburgh, UK; Bulstrode et al., 2017); anti-Green fluorescent protein (GFP) (rabbit or goat; Abcam); anti-Gsx2 (rabbit; Merck); anti-Histone H3 (phospho S10) (rabbit; Abcam); anti-mCherry (chicken; Abcam); anti-Olig2 (rabbit; Millipore); anti-Pax6 (mouse; clone AD2.38, described in Engelkamp et al., 1999); anti-pSmad1/5/9 (rabbit; Cell Signalling); anti-Shh (rabbit; kindly provided by Genentech Inc.); anti-Slc17a7 (formerly Vglut1) (rabbit; Synaptic Systems); anti-Sox9 (rabbit; Millipore); anti-Turbo GFP (mouse; Origene).

For fluorescence immunostaining, sections were incubated with secondary antibodies (1:200) coupled with Alexa Fluor 488 (Abcam) or 568 (Abcam) or Cy3 (Abcam) and then incubated with diamidino-2-phenylindole (DAPI; 1:1000; Thermo Fisher Scientific) before being mounted with Vectashield HardSet (Vector Laboratories) or ProLon Gold Antifade Mountant (Thermo Fisher Scientific).

For colorimetric immunostaining, sections were incubated with biotinylated secondary antibodies followed by Avidin Biotin Complex (ABC kit, Vector laboratories) and then diaminobenzidene (DAB, Vector Laboratories). Sections were mounted in DPX (Sigma).

### In situ hybridization

Riboprobes used in this study were as follows. *Ascl1* and *Gad1* (kindly provided by Francois Guillemot, Francis Crick Institute, UK); *Dlx1* (kindly provided by Yorick Gitton, INSERM Institut de la Vision, France); *Foxg1* (kindly provided by Vassiliki Fotaki, University of Edinburgh, UK); *Gsx2* (kindly provided by Kenneth Campbell, Cincinnati Children’s hospital, USA); *Gsx1* and *Neurog2* (kindly provided by Thomas Theil, University of Edinburgh, UK); *Prdm13* (kindly provided by Tomomi Shimogori, Riken Centre for Brain Science, Japan). Templates for *Robo3*, *Rlpb1*, *Pde1b*, *Igsf11*, *Heg1* and *Neurod4* riboprobes were synthesised from mouse embryonic brain cDNA using the following PCR primers (convention: gene, forward primer, reverse primer). *Heg1*, 5’-ACTTCCAAATGTCCCCATACAC-3’, 5’-CCAGCCCAATCTATTAAAGTGC-3’; *Igsf11*, 5’-TCAGTGCCCTCTCTTCCG-3’, 5’-CAGGCCACTTCACACACG-3’; *Neurod4*, 5’-TGGAATGCTCGGAACCTTAA-3’, 5’-TACAGGAACATCATAGCGGG-3’; *Pde1B*, 5’-GCTGACTGATGTGGCAGAAA-3’, 5’-AGAATCCCAATGGCTCCTCT-3’; *Rlbp1*, 5’-TTCCTCCTGCGCTTCATC-3’, 5’-TTGGGATGAGGTGCCACT-3’; *Robo3*, 5’-GCTGTCCTCCGTGATGATTT-3’, 5’-AAATTGTGGTGGGACGTGAA-3’.

Riboprobes were labelled with digoxigenin (DIG) or dinitrophenol (DNP). DIG labelled probes were synthesised using a DIG RNA labelling mix (Roche). DNP labelled probed were synthesised using a similar process but with the DIG labelling mix replaced by a 20x NTP stock solution (20mM each ATP, GTP, CTP, 13mM UTP, Thermo Fisher Scientific) and a 20x DNP-11UTP stock solution (7mM; Perkin Elmer). For fluorescence in situ hybridisation, probes were detected using an anti-DIG peroxidase (Roche) or anti-DNP peroxidase (Tyramide Signal Amplification (TSA) Plus DNP (HRP) kit, Akoya Bioscience) followed by TSA-plus cyanine 3 or TSA-plus fluorescein (Akoya Bioscience). When required, amplification was achieved using TSA Biotin (Akoya Bioscience). For fluorescence double in situ hybridization, the probes were detected sequentially, and the slides were incubated in 10 mM HCl before detection of the second probe. When fluorescence in situ hybridisation was followed by immunofluorescence, a microwave antigen retrieval step (20 minutes in 10mM sodium citrate) was required at the end of the in situ hybridisation protocol prior to proceeding with immunostaining.

### Bulk RNAseq analysis of *Pax6* cKO, *Pax6 Gsx2* double cKO and control cortex

Pregnant mice were killed by cervical dislocation following isoflurane overdose and embryos were removed. Quality control prior to RNAseq involved the use of immunohistochemistry on one cerebral hemisphere from every *Pax6* cKO or *Pax6 Gsx2* double cKO (dcKO) embryo and from all control littermates to check for efficient deletion of Pax6 or Pax6 and Gsx2. The other hemisphere from brains that passed this quality control was processed for RNAseq as follows. The EGFP reporter was used to guide the dissection to ensure only EGFP+ cortex was included. The rostral and caudal halves of each cortex were separated and frozen on dry ice. Cortical pieces from three E13.5 embryos or four E12.5 embryos were pooled to produce each sample. Total RNA was extracted from each pool using an RNeasy+ Micro kit (Qiagen). Poly-A mRNA was purified and TruSeq RNA-Seq libraries (Illumina) were prepared and sequenced (100 base paired-end; Illumina, HiSeq v3).

Sequencing gave an average of 108 million reads per sample. Following adapter trimming and standard quality checks, reads were aligned using STAR v. 2.4.0a to GRCm38.p3 (mm10) genome build obtained from Ensembl release 77. Reads were counted using featureCounts v. 1.4.5-p from the Subread package and only fragments with both reads properly aligned to exon regions were counted.

DESeq2 (Love et al., 2014) and edgeR (Robinson et al., 2010) were used to identify genes that were differentially expressed (DE) between conditions. When using DESeq2, package functions were used with default parameters and DE genes were defined as those with adjusted p-value ≤ 0.05 after Benjamini-Hochberg correction for multiple testing. When using edgeR, sample dispersion was estimated using default parameters and pairwise Exact Tests were used to compare expression under different conditions: DE genes were defined as those with false discovery rate (FDR) ≤ 0.05.

We included all genes identified by either of these packages for further consideration as DE genes (most DE genes were identified by both packages).

### Single cell RNAseq (scRNAseq) analysis of *Pax6* cKO and control cortex

Pregnant mice were killed by cervical dislocation following isoflurane overdose and embryos were removed. Embryos were placed in ice-cold Earle’s Balanced Salt Solution (EBSS) and examined under a fluorescence stereomicroscope for the presence of EGFP. EGFP-positive brains were used for the scRNAseq experiment; EGFP-negative brains from littermates were used as negative controls for EGFP signal calibration in subsequent fluorescence-activated cell sorting (FACS). The rostral half of the cortex from one hemisphere of each GFP-positive brain was isolated and used for tissue dissociation; the other hemisphere was fixed with 4% PFA and used in immunohistochemistry to check for efficient deletion of Pax6. The tails from the embryos were collected for genotyping to confirm *Pax6^flox^* copy number.

Tissue was dissociated in 30 U/mL papain reagent containing 0.4% DNase (Worthington) for 30 mins at 37C. Samples were gently triturated using glass pipettes (BrainBits LLC) and sieved through a 40μm cell strainer (pluriSelect) to remove cell clumps. Dissociated cells were resuspended in basic sorting medium containing 2% heat-inactivated fetal bovine serum. Trypan blue was added to sample aliquots to allow quantification of cell density and viability. EGFP-positive cortical cells were selected using the FACSAria II (BD Biosciences). Cells were stained with DAPI to select against cell debris, cell doublets and dead cells. The system was then calibrated with EGFP-negative cells, before selecting approximately 20,000 EGFP-positive cells for each sample. FACSDiva 8.0.1 software (BD Biosciences) was used to process all flow cytometry data.

Three embryos from a pregnant mouse were used to prepare each E13.5 library. For E14.5 libraries, *Pax6* cKO embryos from three pregnant mice and control embryos from two pregnant mice were pooled together. All libraries from E13.5 and E14.5 were produced from male embryos; none of the cells across all four libraries expressed *Xist*.

The 10x Genomics Chromium Controller and Single Cell 3’ Reagent Kits (v.2 for E13.5; v.3 for E14.5) were used for library construction, as per manufacturer’s instructions. Cell density and viability were quantified using an automated cell counter as part of the quality control before loading onto the controller chip. Each sample was its own library. For each library, 3000-4000 cells were targeted for capture. Each library was constructed with a specific sample barcode and standard Illumina paired-end constructs. Two libraries from each embryonic age were sequenced simultaneously using the Illumina NovaSeq 6000 S1 flow cell. Raw sequencing reads were processed through the CellRanger pipeline (v.2 at E13.5; v.3 at E14.5; 10x Genomics) to produce a gene-cell count matrix using the mm10 mouse genome as reference. Approximately 60,000-100,000 reads per cell were generated to achieve optimal sequencing depth.

All data were subjected to SoupX (Young and Behjati, 2020) to remove ambient RNA captured in droplets. To filter out cells giving low-quality data, cut-offs on unique molecular identifier (UMI), gene count and mitochondrial gene percentage were applied. Cells with gene counts > 3 standard deviations from the mean were excluded: depending on sequencing depth, cells expressing <∼1000 and >∼5500 genes were typically excluded. Cells expressing extremely high numbers of mitochondrial genes (>5% at E13.5; >10% at E14.5) were also excluded.

Clustering analysis was performed using the Seurat v.3.0 bioinformatics pipeline (https://github.com/satijalab/seurat). For each dataset, a Seurat object was created. UMI counts were natural log-normalized using log1p with scale factor of 10,000 using the NormalizeData function. Individual gene expression on Uniform Manifold Approximation and Projection (UMAP) was constructed with natural log-normalized expression using the FeaturePlot function. Expression values for each gene across all cells was standardized using z-score transformation implemented in the ScaleData function. A cell cycle phase was assigned to each cell using the Seurat pipeline. The datasets were not regressed with cell cycle scores as this might have hindered identification of the physiological relevance of cell cycle in the temporal progression of cortical neurogenesis. For integrated analysis across the control and *Pax6* cKO datasets, normalization and feature selection for highly variable genes in each library were performed independently. Based on the selected features, the control and *Pax6* cKO datasets were passed through the FindIntegrationAnchors function to identify the cross-dataset anchors for data integration. The resulting integration anchors were then implemented in the IntegratedData function, producing an integrated Seurat object. Next, the most variable genes in the integrated dataset were identified and used for principal component analysis. The most statistically significant principal components were used as inputs for non-linear dimensionality reduction using UMAP implemented in the FindCluster function at a resolution between 0.8 and 1.6 to determine cellular clusters based on k-nearest neighbours and shared nearest neighbour (SNN) graphs. Top cluster marker genes were identified and differential expression analysis (DEA) across two different cellular clusters was carried out using Model-based Analysis of Single-cell Transcriptomics (MAST) implemented in the FindAllMarkers function.

Cell lineage or pseudotime inference was carried out using the Slingshot algorithm (https://github.com/kstreet13/slingshot) (Street et al., 2018). A matrix with reduced dimensionality and cell clustering assignments was taken as input by Slingshot. With the radial glial progenitor (RGP) as the initial cell state (or root node), lineage trajectories and branch points were inferred by connecting the cluster medoids with a minimum spanning tree. Patterns of gene expression along pseudotime were computed using the tradeSeq algorithm (htts://statomics.gitbub.io/tradeSeq/index.html) (Van den Berge et al., 2020). Gene expression along pseudotime was normalized using log1p.

RNA velocity estimation was generated using the Velocyto (https://github.com/velocyto-team) and scVelo (https://github.com/theislab/scvelo) software packages. Count matrices with cells that had passed quality control were used as inputs for Velocyto. All velocity embeddings were estimated using the stochastic model. For visualization purposes, the RNA velocity embeddings were visualized using the UMAP coordinates produced from the previous cell clustering analysis in Seurat.

### BrdU and Ethynyldeoxyuridine (EdU) labelling

Pregnant females were given a single intraperitoneal dose of BrdU or EdU (10mg mL^-1^; Thermo Fisher Scientific). Subsequent immunohistochemistry was as described above. Cells labelled with EdU were visualised using a Click-iT EdU Alexa Fluor 647 Imaging Kit (Thermo Fisher Scientific).

### Whole-cell electrophysiology

Postnatal mice were anesthetized with isoflurane and sacrificed by decapitation, the brain was quickly removed into artificial cerebrospinal fluid (aCSF) (80 mM NaCl, 2.5 mM KCl, 1.25 mM NaH_2_PO_4_, 25 mM NaHCO_3_, 10 mM glucose, 90 mM sucrose, 0.5 mM CaCl_2_, 4.5 mM MgSO_4_) and 400 µm coronal slices were prepared in ice-cold aCSF solution using a vibrating microtome (VT1200S, Leica, Germany). Slices were transferred to aCSF solution for 30 minutes at 35 °C and current clamp whole-cell recordings were performed using a Multiclamp 700A amplifier (Molecular Devices, Palo Alto, USA). Signals were filtered online at 10 kHz and digitized at 40 kHz and traces stored on the computer using Signal 2 software (Cambridge Electronic Design, UK). Borosilicate glass electrodes (Harvard Apparatus, UK) with a resistance of 3.5-7.5 MΩ (Digitimer Research Instruments, UK) were filled with the following solution: 135 mM KMeSO_4_, 8 mM NaCl, 10 mM HEPES, 0.5 mM EGTA, 0.5 mM Na-GTP, and 4 mM Mg-ATP, 0.5 mM EGTA, pH 7.3, osmolarity adjusted to 285 mOsm. The chamber was perfused at a rate of 3-4mL/min with carbonated (95% O2, 5% CO2) external solution containing: 130 mM NaCl, 2.5 mM KCl, 1.25 mM NaH_2_PO_4_, 25 mM NaHCO_3_, 10 mM glucose, 2.5 mM CaCl_2_, 1.5 mM MgSO_4_ at 32-35 °C.

Access resistance was monitored online throughout the recording and if it dropped >20% the recording was discarded and no further analysis was carried out. In current clamp small incremental voltage steps (10/25 pA) were applied for 500 ms to measure cell intrinsic properties. Membrane potential was held at -70mV. Feature extraction was done blind to genotype using custom scripts written in Signal 2. Some raw .csf recordings of cortical cells were lost and are no longer available.

Action potential (AP) features were extracted from single APs at rheobase. AP threshold was defined as the time at membrane potential at which the slope first exceeded 30 V/s. AP amplitude was defined as height from AP threshold to peak. Afterhyperpolarization (AHP) amplitude (mV) was defined as potential difference between AP threshold and AHP trough. AP latency was defined as the time between start of current step and first AP peak. Properties of trains of APs were determined in response to a current step double rheobase. First-second adaptation ratio and first-last adaptation ratio were calculated by dividing instantaneous frequency of the second/last AP by the first AP instantaneous frequency (this measures the interval between spikes). Spikelet amplitude (mV) was defined as the difference between peak voltage and steady-state voltage during the current step. Input resistance and membrane time constant (tau) were measured using the average of 20 membrane potential responses to negative current steps (-10 or -25 pA) lasting 500ms. Membrane time constant (tau) was calculated by fitting a single exponential (10%-90%) to the membrane potential response curve. Capacitance was then calculated using tau/input resistance = capacitance. To test the effects of tetrodotoxin (TTX) on spikelets, we used spikelet-inducing current steps (square pulses 0.5s long; 60 sweeps over 60s) while the chamber was perfused with external solution either containing TTX (300nM) or without TTX, to ensure there was no spikelet wane with persistent stimulation. To show TTX blockade of spikelets, which occurred over 30 sweeps, we compared the average amplitudes of spikelets over the first and last 10 sweeps.

Data were analysed statistically and by unsupervised hierarchical clustering. For pairwise comparisons, N’s were animals. Data was analysed in R (packages: cluster, ggpubr, dendextend, Nbclust, PairedData and tidyverse). Pre-analysis routines were carried out to inspect data normality by plotting residual plots (qqplots). If data was not normally distributed, a non-parametric test was used. Data for clustering was scaled using base R scale function. For unsupervised clustering, the agglomerative Ward’s linkage method was used after calculating Euclidean distances to minimize the variance within clusters (Ward, 1963). Data features used for clustering: AHP amplitude, AP halfwidth, AP threshold, input resistance, capacitance, number of spikes in response to 500ms current step at double rheobase current, first-spike latency, first to second AP adaptation ratio and first to last AP adaptation ratio. Silhouette coefficients were calculated using Nbclust R package. Signal analysis scripts available at Marcos Tiago, pax6_codes (2019-2021) online GitHub repository (https://github.com/TiagoMarcos).

### In utero injection and electroporation

Pregnant mice were maintained under inhaled isoflurane anaesthesia for the duration of the procedure. The uterine horns were exposed. In some experiments, 1-2 μl of Vismodegib (5mM solution in DMSO; Stratech) or DMSO alone was injected into the lateral ventricle of each embryo’s brain with a glass micropipette. In other experiments, plasmids were injected into the lateral ventricle of each embryo’s brain at 1 to 4 mg mL^-1^, the embryo in the uterus was placed between tweezer-type electrodes (CUY650; Nepagene) and an electroporator (CUY21E; Nepagene) was used to deliver 6 pulses (30 V, 50 ms each, 950 ms apart). In both cases, the uterine horns were replaced, the abdominal wall was sutured and animals recovered. Processing was as described above.

Plasmids used in this study were: CAG-FoxG1-IRES-mCherry (kindly provided by Goishi Myioshi, Tokyo Women’s Medical University, Japan); Scrambled shRNA control in pGFP-V-RS shRNA Vector (TR30013, Origene); Smoothened shRNA in pGFP-V-RS shRNA Vector (TG510788, Origene).

### Organotypic slice cultures

Pregnant mice were killed by cervical dislocation following isoflurane overdose and embryos were removed. Embryonic brains were embedded in 3.5% (w/v) low melting point agarose (Lonza) in phosphate buffered saline. Coronal sections were cut at a thickness of 300μm using a vibratome. The slices were transferred onto a polycarbonate culture membrane (Whatman) floating on minimum essential medium (MEM; Gibco) supplemented with fetal bovine serum (FBS) and incubated at 37°C, 5% CO_2_. After an hour of incubation, the FBS -supplemented medium was replaced with serum-free neurobasal medium (Gibco) supplemented with B27 (1:50 dilution; Gibco) and N2 supplement (1:100 dilution; Gibco).

In some experiments, after an hour of incubation, the ventral telencephalon of one of the hemispheres was removed from the section while the other hemisphere was left intact. In other experiments, after an hour of incubation, pharmacological treatments were carried out. For Bmp4 treatment, Bmp4 (R&D Systems) or vehicle alone was added to the culture medium to achieve a range of concentrations while maintaining the concentration of all other components constant.

Cyclopamine treatment was done in one of two ways. Either Affi-gel agarose beads (Bio-Rad) were soaked overnight in cyclopamine (4 mg mL^-1^; Toronto Research Chemicals) or vehicle alone and transferred onto the cortex; or cyclopamine (10µM) or vehicle alone were added in solution to the culture medium. After culture at 37°C, 5% CO_2_, telencephalic slices were processed for qRT-PCR, immunohistochemistry or in situ hybridization as described above.

### Dissociated cell culture

Pregnant mice were killed by cervical dislocation and embryos were removed. Tissue samples were taken from each embryo for genotyping to identify whether they were *Pax6* cKO or control (Simpson et al., 2009). GFP+ cortices were dissociated using a papain dissociation kit (Worthington) and trituration using glass pipettes (BrainBits LLC). Cells were cultured on poly-L-ornithine and laminin-coated coverslips in 24-well plates at 150,000 cells per well in serum-free 2i Medium (Merck). After 24h, Smoothened Agonist (SAG; Abcam) dissolved in DMSO was added to make final SAG concentrations of 1.25nM, 2.5nM, 5nM, 10nM, 20nM, 40nM, 80nM or 160nM. An equivalent number of cultures of each genotype had DMSO only added to give each of the concentrations present at each of the SAG concentrations (0.025%, 0.05%, 0.1%, 0.2%, 0.4%, 0.8%, 1.6% and 3.2%). Some cultures had both 160nm SAG and 10mM EdU added. Cells were fixed with 4% PFA 48h later.

Antibodies for immunocytochemistry included anti-Tubb3 (1:500; chicken; Novus Biologicals); the others were as described above. DIG labelled *Dlx1* probes were synthesized and used for fluorescence in situ hybridization as described above. EdU was detected using a Click-iT EdU Alexa Fluor 647 Imaging Kit (Thermo Fisher Scientific). Cells were counterstained with DAPI (1:1000) and coverslips were mounted with Vectashield HardSet (Vector Laboratories).

All cultures were imaged at x20 and imported to Fiji software (https://imagej.net/software/fiji/) for cell counting. Each coverslip was divided into 25 areas (Fig. S10C) and counting was done using the Cell Counter plug-in in 5 areas selected at random using a random number generator.

### Microscopy

Imaging was carried out using Leica brightfield or epifluorescence microscopes or a Nikon A1R confocal microscope. Microscope images were acquired using the Leica Application Suite X (LAS X) or Nikon NIS-Elements software.

## ACKNOWEDGEMENTS

We thank Ugo Borello, Matt Colligan, Petrina Georgala, Anisha Kubasik-Thayil, Da Mi, Soham Mitra, Elena Purlyte, Thomas Theil, Daphne Triantafyllou and staff in Bioresearch and Veterinary Services and Edinburgh Genomics for their contributions. This research was funded by the Medical Research Council UK (Mr/J003662/1; Mr/N012291/1), the Biotechnology and Biological Sciences Research Council UK (Bb/N006542/1), the Muir Maxwell Epilepsy Centre, Simons Initiative for the Developing Brain (SFARI – 529085), Principal’s Career Development and Edinburgh Global Research Scholarships (K.B.T.) and a Scholarship from the Ministry of Higher Education, Malaysia (M.F.A.R.)

## Supplementary Figures

**Figure S1:**
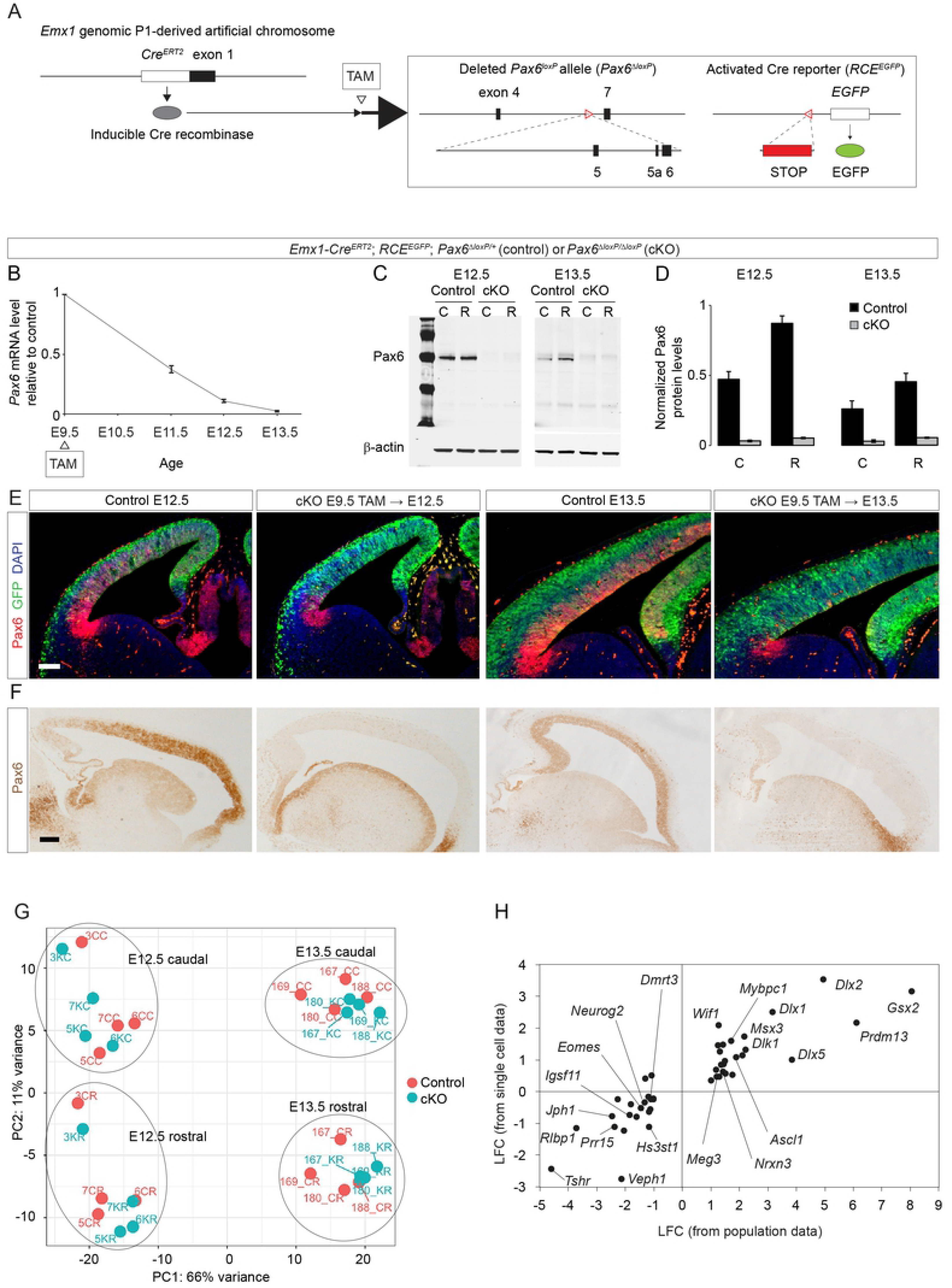
Validation of Pax6 removal and RNAseq data. **(A)** Frequently used alleles: *Emx1-Cre^ERT2^* producing tamoxifen (TAM)-inducible Cre recombinase (Kessaris et al., 2006); *Pax6^loxP^*, from which paired domain-encoding exons were removed by Cre recombinase (*Pax6^ΔloxP^*), rendering it non-functional (Simpson et al., 2009); *RCE^EGFP^*, a Cre reporter producing R26R CAG-boosted enhanced green fluorescence protein (EGFP) (Miyoshi et al., 2010). Mice with deletions in both copies of *Pax6* were designated conditional knock-outs (*Pax6* cKOs); those with a deletion in just one copy served as controls. **(B)** Quantitative RT-PCR measurements of *Pax6* mRNA levels, normalised to those of *Gapdh*, after tamoxifen administration at E9.5 were used to calculate average (±sem) ratios between levels in *Pax6* cKO and control littermates (n=3 embryos from 3 litters at each age). **(C)** Western blots showing Pax6 protein expression in the rostral (R) and caudal (C) cortex of control and *Pax6* cKO littermates at E12.5 and E13.5 after tamoxifen administration at E9.5. **(D)** Quantification of Western blots at E12.5 and E13.5 after tamoxifen administration at E9.5. Pax6 protein levels were measured relative to β-actin levels. Average levels (±sem) were calculated (n=3 independent repeats in each region at each age; in each case, levels in *Pax6* cKOs and controls differed with p<0.01 in Student’s t-tests). (Note that Pax6 protein levels in control rostral cortex were almost double those in caudal control cortex at each age, in agreement with previous observations: Manuel et al., 2007). **(E)** Expression of green fluorescent protein (GFP) and Pax6 protein in coronal sections through the cortex of control and *Pax6* cKO embryos at E12.5 and E13.5 after tamoxifen administration at E9.5. GFP was activated by most cortical cells and Pax6 protein was lost from most cortical radial glial progenitors across almost the entire cortex, excluding a narrow ventral pallial domain where *Emx1-Cre^ERT2^* was not expressed. Scale bar: 0.1mm. **(F)** Expression of Pax6 protein in sagittal sections through the cortex of control and *Pax6* cKO embryos at E12.5 and E13.5 after tamoxifen administration at E9.5. Scale bar: 0.1mm. **(G)** Principal component analysis on RNAseq data from control caudal (CC) and control rostral (CR) and *Pax6* cKO caudal (KC) and *Pax6* cKO rostral (KR) cortex at E12.5 and E13.5 showing major clustering based on region and age. **(H)** Significant LFCs with values ≥ 1 or ≤ -1 obtained from bulk RNAseq on samples of E13.5 rostral cortex plotted against significant LFCs calculated from scRNAseq data (by carrying out differential expression analysis on average gene expression levels obtained from random subsets of rostral E13.5 *Pax6* cKO and control cells). By way of example, genes are named for some datapoints.

**Figure S2:**
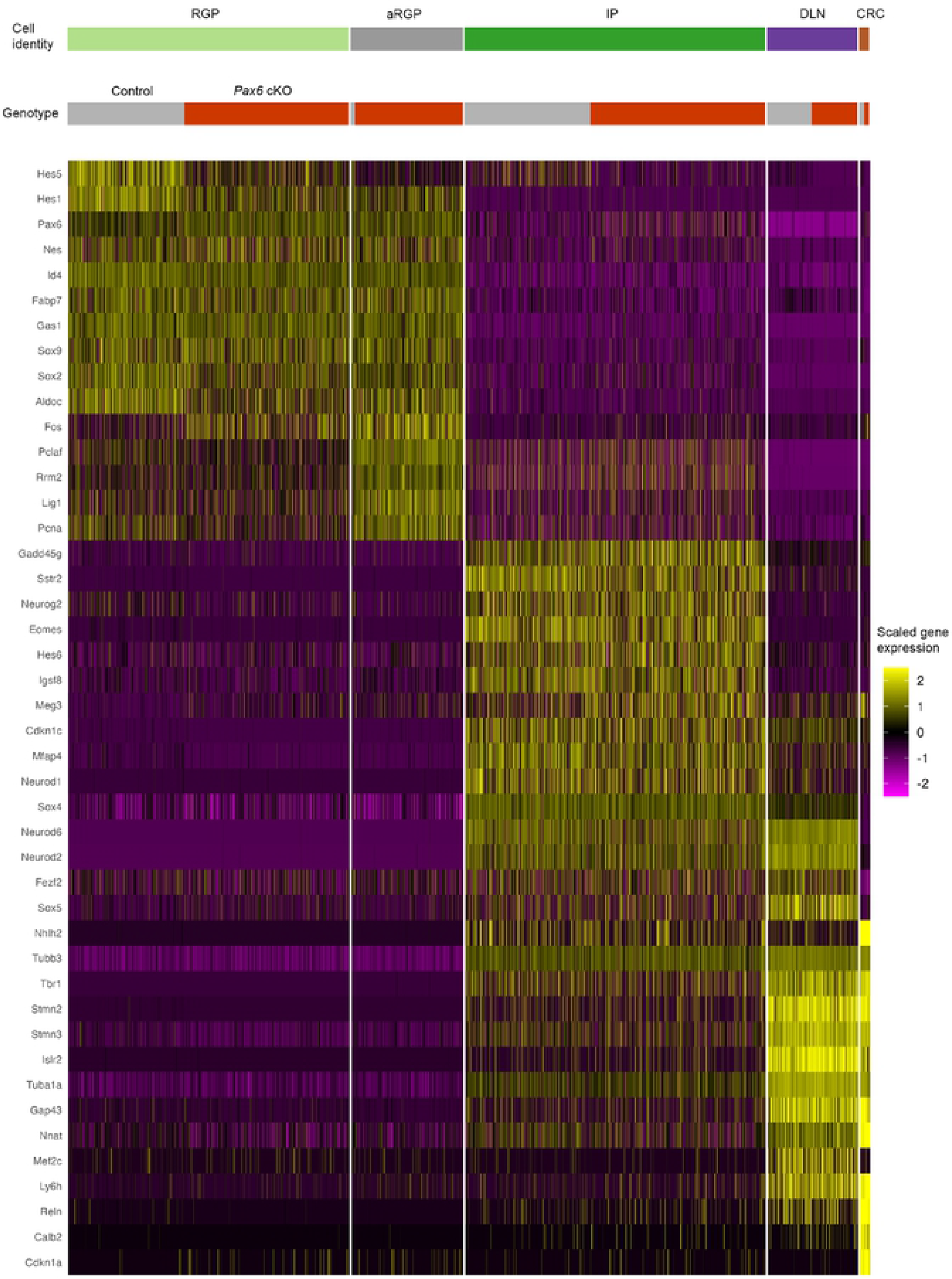
Expression profiles of major cell types in *Pax6* cKO and control cortex at E13.5. Heat map of gene expression in RGPs (radial glial progenitors), aRGPs (atypical RGPs), IPs (intermediate progenitors), DLNs (deep layer neurons) and CRCs (Cajal-Retzius cells) in E13.5 control and *Pax6* cKO cortex.

**Figure S3:**
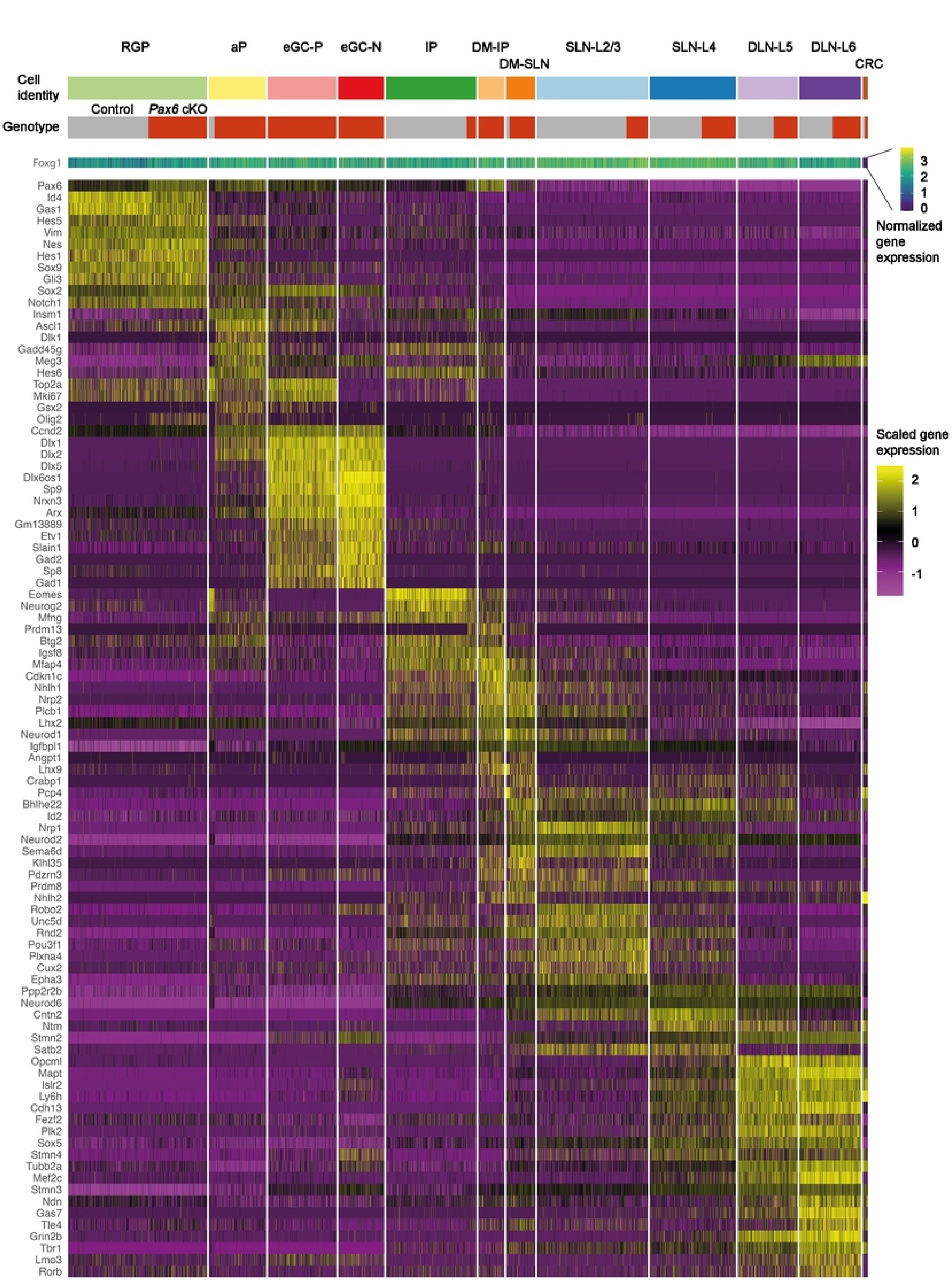
Expression profiles of major cell types in *Pax6* cKO and control cortex at E14.5. Heat map of gene expression in RGPs (radial glial progenitors), aPs (atypical progenitors), eGC-Ps and eCG-Ns (proliferating or non-proliferating ectopic GABAergic cells), IPs (intermediate progenitors), DM-IPs and DM-SLNs (intermediate progenitors or superficial layer neurons in dorsomedial cortex), SLN-L2/3s and SLN-L4s (layer 2/3 or layer 4 superficial layer neurons), DLN-L5s and DLN-L6s (layer 5 or layer 6 deep layer neurons) and CRCs (Cajal-Retzius cells) in E14.5 control and *Pax6* cKO cortex. Note the increased levels of *Pax6* mRNA in *Pax6* cKO cells, which was anticipated because the loss of Pax6 protein would have removed the negative feedback constraining the transcription of residual *Pax6* coding sequence (Manuel et al., 2007).

**Figure S4:**
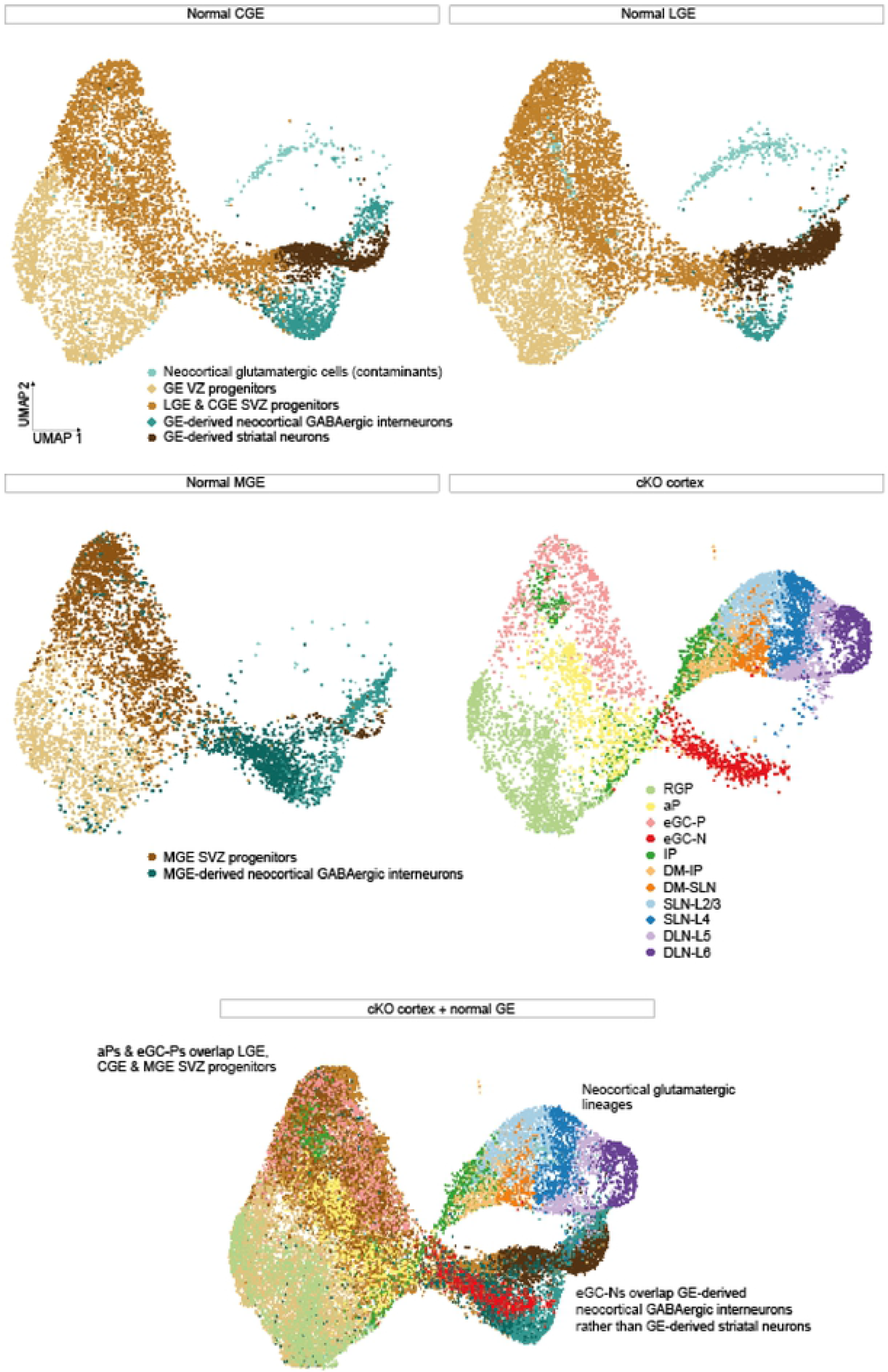
Integrated scRNAseq data from E14.5 *Pax6* cKO cortex and normal ganglionic eminences. UMAP plots of scRNAseq data from normal E14.5 ventral telencephalon (Mayer et al., 2018) integrated with our scRNAseq data from E14.5 *Pax6* cKO cortex. Abbreviations for cortex in Fig. 2A. Additional abbreviations: GE, ganglionic eminence; LGE, lateral ganglionic eminence; CGE, caudal ganglionic eminence; MGE, medial ganglionic eminence; VZ, ventricular zone; SVZ, subventricular zone.

**Figure S5:**
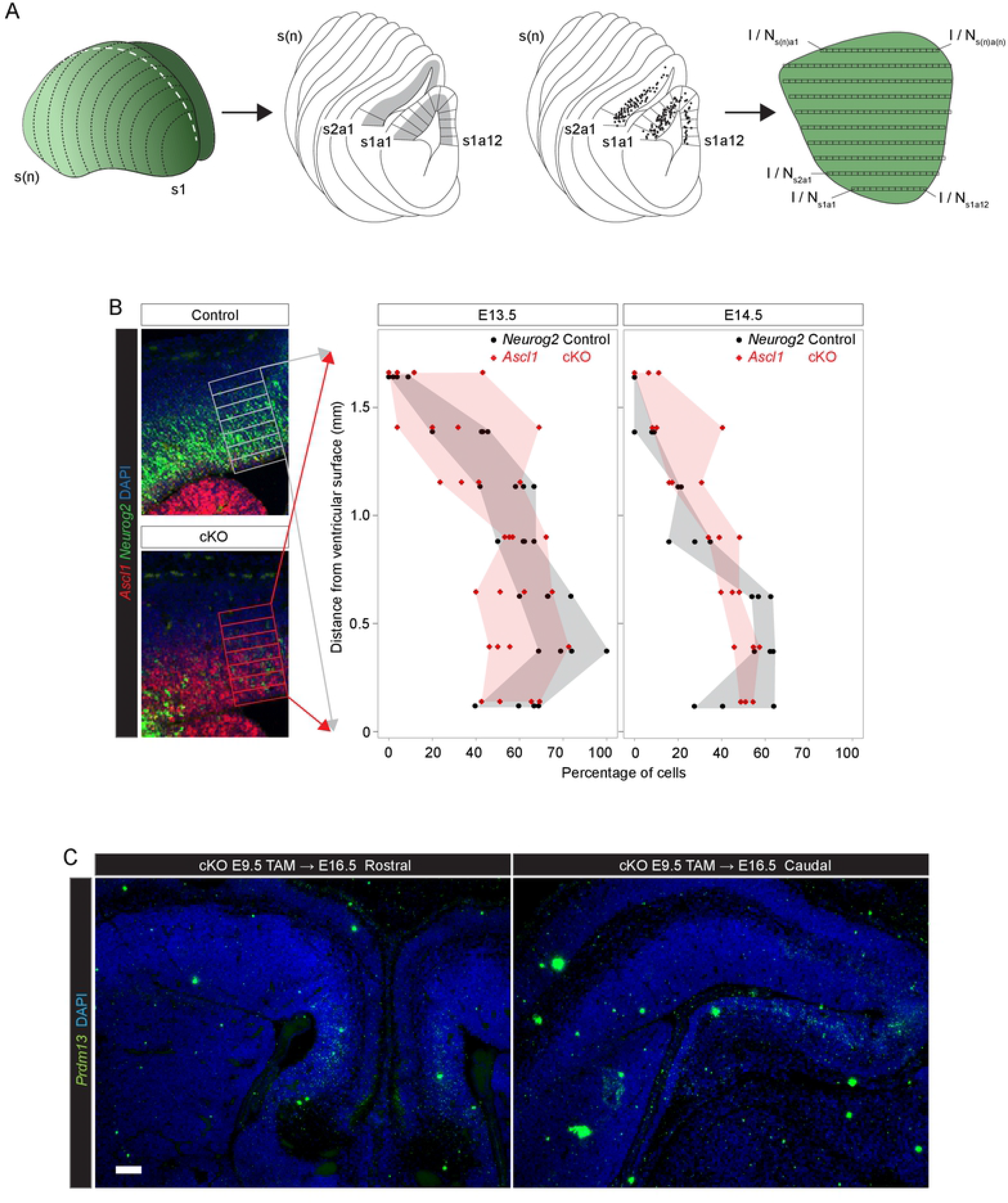
A*s*cl1, *Neurog2* and *Prdm13* expression in control and *Pax6* cKO cortex. **(A)** Method for obtaining a surface-view reconstruction of labelling across the cortex from a series of equally-spaced coronal sections (s1 to s(n)). Relative intensity of label (I) or labelled cell number (N) were measured in areas (a1 to a(n)) with average width (through cortical depth) of 0.1 mm in each section. Values were then laid out on a flattened representation of the cortical surface and maps generated by interpolation. **(B)** Quantification showing the similarity in the distributions with depth of the proportions of cells expressing *Neurog2* mRNA in control lateral cortex and *Ascl1* mRNA in *Pax6* cKO lateral cortex at E13.5 and E14.5. Data were obtained using 25µm × 100µm bins, as shown, from the lateral cortex in centrally located coronal sections through the brains of 4 *Pax6* cKOs and 4 controls at E13.5 and 3 *Pax6* cKOs and 3 controls at E13.5. Data-points are for individual animals and shaded areas show the range of values with depth (Source Data Table 1). **(C)** In situ hybridization for *Prdm13* in rostral and caudal *Pax6* cKO cortex at E16.5 after tamoxifen administration at E9.5. Scale bar: 0.1mm.

**Figure S6:**
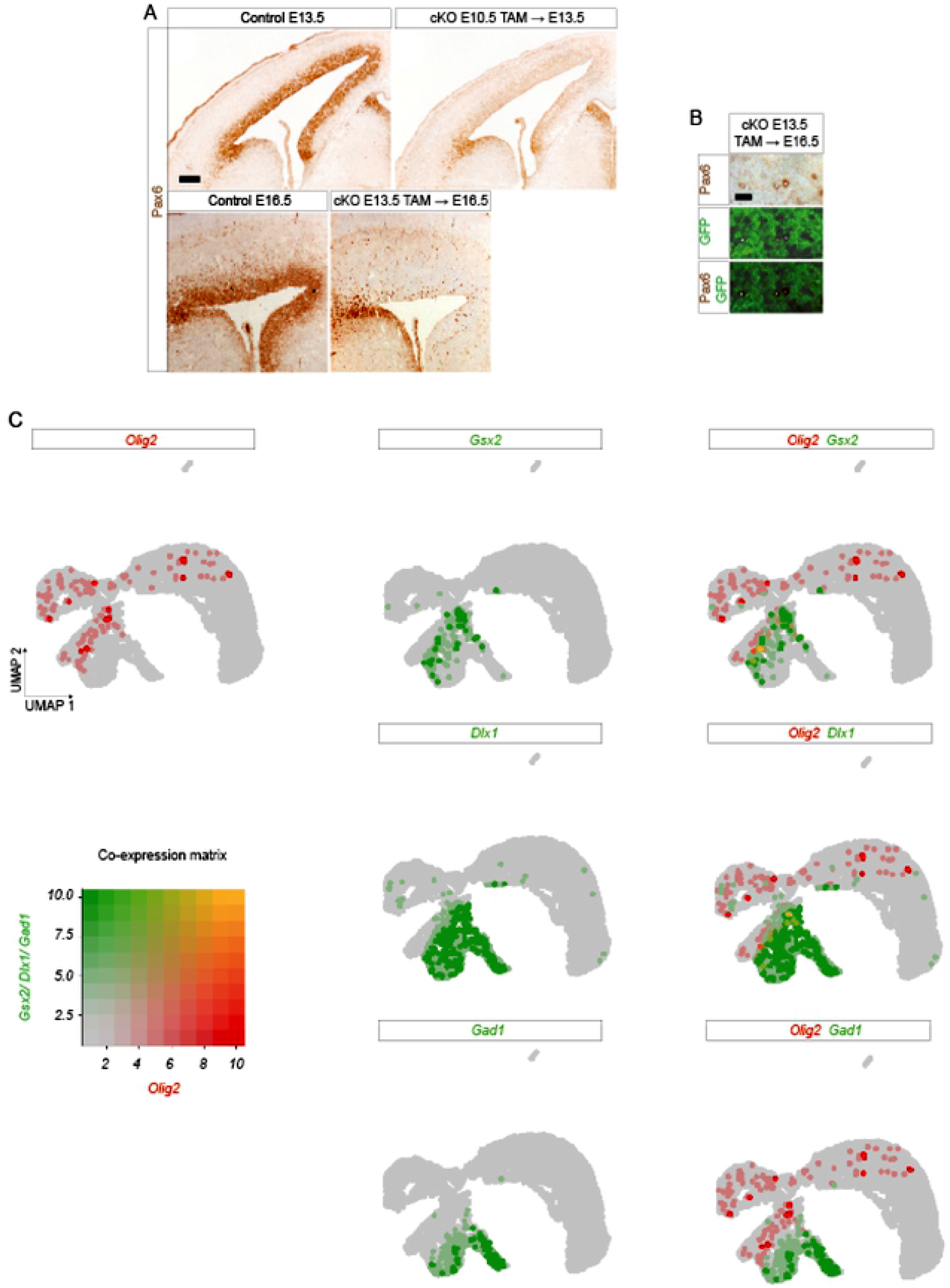
Pax6 deletion and Olig2 expression in *Pax6* cKO cortex. **(A)** Expression of Pax6 protein in coronal sections through the cortex of control and *Pax6* cKO embryos at E13.5 after tamoxifen administration at E10.5 and at E16.5 after tamoxifen administration at E13.5. Scale bar: 0.1mm. **(B)** Expression of Pax6 protein and GFP in sections through the *Pax6* cKO cortex at E16.5 after tamoxifen administration at E13.5. The vast majority of cortical RGPs were Pax6-negative. Many of the few remaining Pax6+ cells were GFP-negative (examples marked with white dots). Scale bar: 0.01mm. **(C)** UMAP plots from *Pax6* cKO cortex at E14.5 showing relative expression and co-expression levels of *Olig2*, *Gsx2*, *Dlx1* and *Gad1* in each cell (on a 0-10 scale, 10 was the highest expression level of the gene in question).

**Figure S7:**
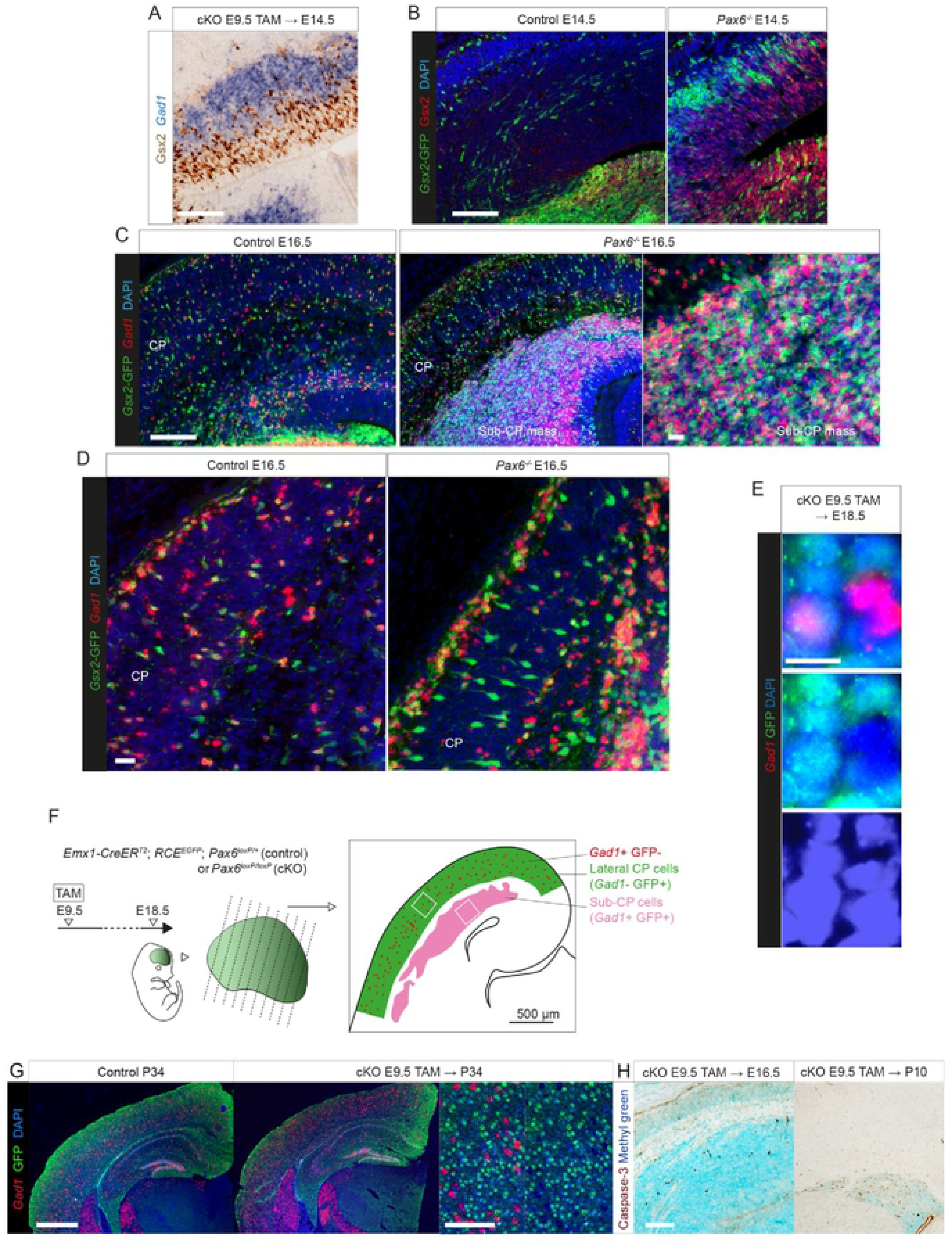
Analyses of eGC fates. **(A)** Colorimetric immunoreactivity for Gsx2 and in situ hybridization for *Gad1* in E14.5 *Pax6* cKO lateral cortex after tamoxifen^E9.5^. Scale bar: 0.1mm. **(B)** Fluorescence immunoreactivity for Gsx2 and GFP, the latter marking *Gsx2*-lineage cells, in E14.5 control and *Pax6^-/-^* telencephalon. Scale bar: 0.1mm. **(C,D)** Fluorescence immunoreactivity for GFP+ (*Gsx2*-lineage) and in situ hybridization for *Gad1*+ cells in E16.5 control and *Pax6^-/-^* lateral cortex. Scale bars: 0.1mm and 0.01mm. **(E)** Three *Gad1*+ cells in the CP of an E18.5 *Pax6* cKO (tamoxifen^E9.5^) carrying the *RCE^EGFP^* Cre-reporter. One was *Emx1*-lineage (GFP+). Scale bar: 0.01mm. **(F)** Experimental design for analysis of the effects of Pax6 deletion on cortical cell numbers (Fig. 7H). The *Emx1-Cre^ERT2^* allele with tamoxifen^E9.5^ was used to delete *Pax6*; embryos carried a GFP reporter of Cre activity; embryos were collected at E18.5; regularly-spaced coronal sections were double-stained for GFP protein and *Gad1* mRNA. We measured the surface area of the cortical plate (CP) of the lateral cortex and the surface area of the sub-CP masses of one hemisphere in each section and interpolated to estimate the volumes of these structures in each embryo. We used random sampling (e.g. white boxes) to measure the average densities of cells in each region. We then calculated their total numbers in each region in each embryo. Data from 4 littermate pairs from separate mothers were used to produce Fig. 7H. **(G)** Fluorescence immunoreactivity for GFP (*Emx1*-lineage) and in situ hybridization for *Gad1*+ cells in P34 control and *Pax6* cKO cortex after tamoxifen at E9.5. High magnification images: same region of CP with and without *Gad1* staining. Scale bar: 1mm and 0.1mm. **(H)** Immunoreactivity for Caspase-3 in CP and in sub-CP masses in *Pax6* cKO cortex at E16.5 and at P10 after tamoxifen at E9.5. Scale bar: 0.1mm.

**Figure S8:**
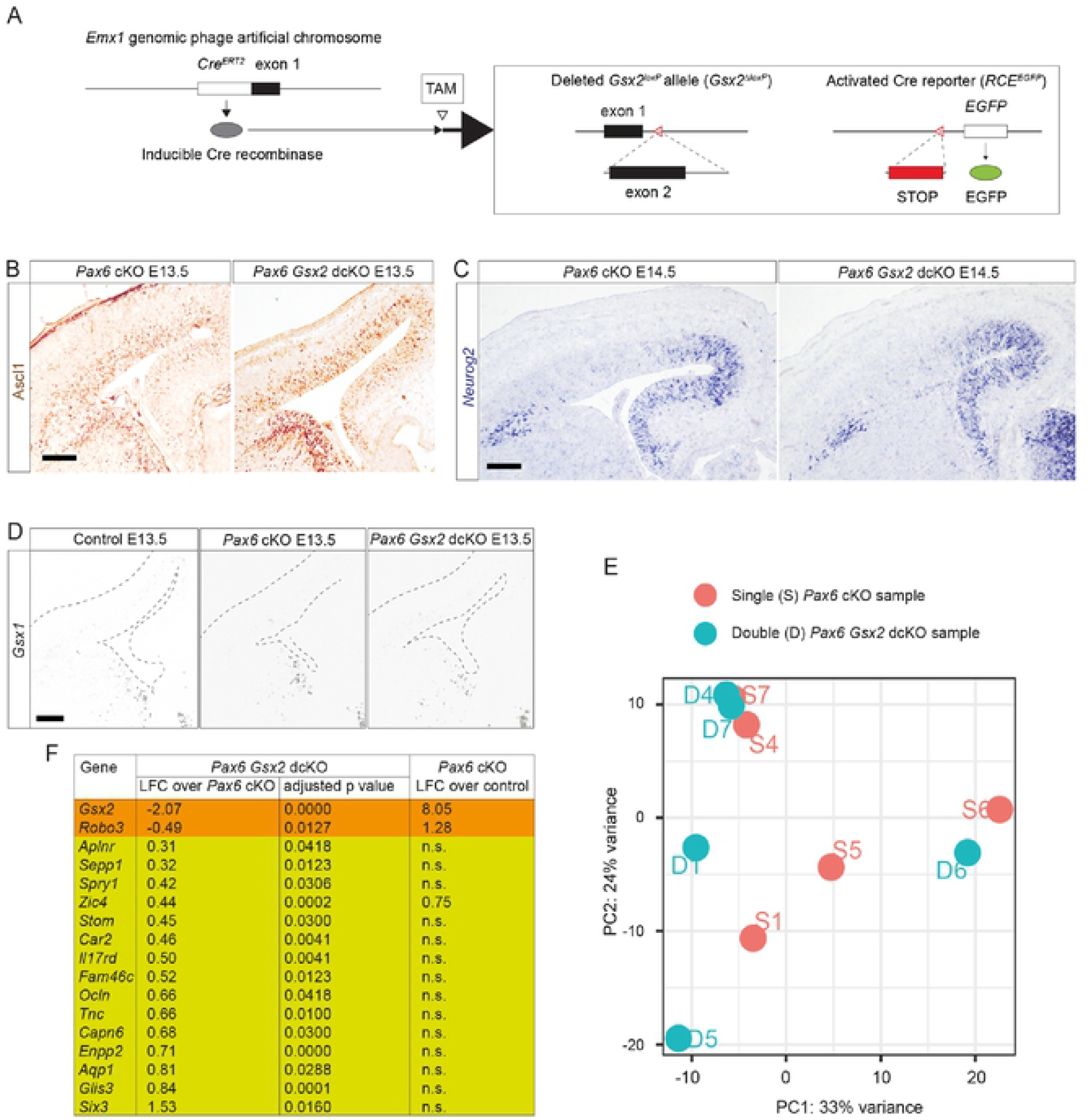
Comparing the effects of losing Pax6 alone versus Pax6 and ectopic Gsx2. **(A)** Alleles used to delete conditionally *Gsx2*: *Emx1-Cre^ERT2^* producing tamoxifen (TAM)-inducible Cre recombinase (Kessaris et al., 2006); *Gsx2^loxP^* (Waclaw et al., 2009); *RCE^EGFP^* (Miyoshi et al., 2010). **(B,C)** Immunoreactivity for Ascl1 and in situ hybridization for *Neurog2* in *Pax6* cKO and *Pax6 Gsx2* dcKO cortex at E13.5 and E14.5. Scale bar: 0.1mm. **(D)** In situ hybridization for *Gsx1* in control, *Pax6* cKO and *Pax6 Gsx2* dcKO cortex at E13.5. Scale bar: 0.1mm. **(E)** Principal component analysis on RNAseq data from *Pax6* cKO (n=5 embryos) and *Pax6 Gsx2* dcKO (n=5 embryos) cortex at E13.5. Data were from the rostral half of the cortex, where the proportion of cells activating *Gsx2* after *Pax6* deletion was highest. **(F)** A list of all genes that were significantly downregulated or upregulated (adjusted p<0.05) in *Pax6 Gsx2* dcKO cortex compared to *Pax6* cKO cortex and their log_2_ fold changes (LFCs) in *Pax6* cKO compared to control cortex (n.s., not significant).

**Figure S9:**
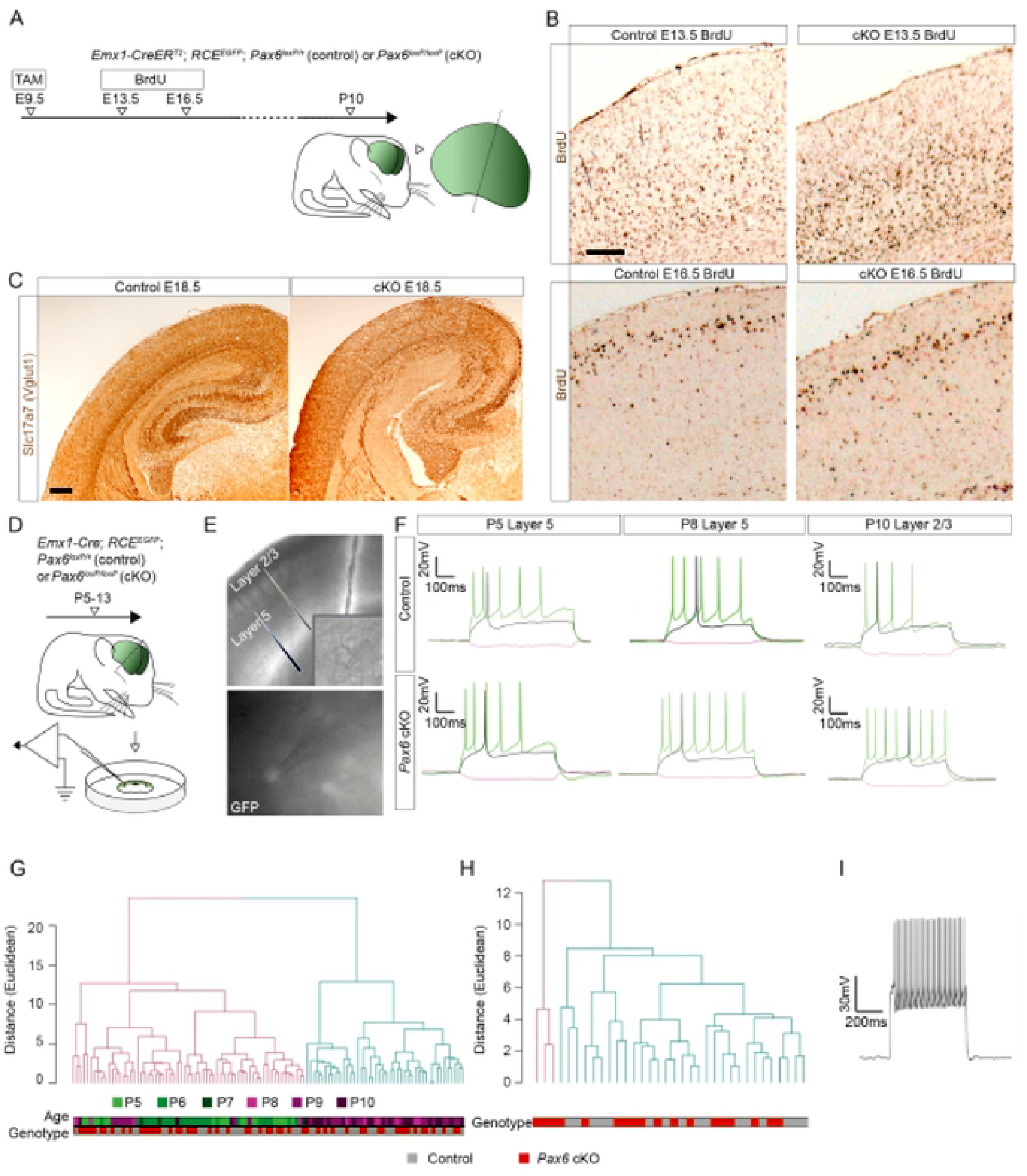
Cortical plate development from *Pax6* cKO RGPs. **(A)** The experimental procedure for (B): tamoxifen (TAM) was administered at E9.5 to generate control and *Pax6* cKO embryos; 5-Bromo-2’-deoxyuridine (BrdU) was injected on either E13.5 or E16.5; brains were sectioned coronally and analysed at P10. **(B)** BrdU-labelled cells in P10 lateral cortical plate; experimental procedure in (A). Scale bar: 0.1mm. **(C)** Immunoreactivity for Slc17a7 (formerly Vglut1) in E18.5 control and *Pax6* cKO cortex after tamoxifen at E9.5. Scale bar: 0.1mm. **(D)** The experimental procedure for (E-I): slices from P5-13 *Emx1*-Cre; *RCE^EGFP^*; *Pax6^loxP/+^* (control) or *Pax6^loxP/loxP^* (*Pax6* cKO) mice were used for electrophysiology. **(E)** GFP+ cells in layers 2/3 and 5 of somatosensory cortex area 1 (S1) were targeted for whole cell current clamp recordings (electrodes are visible in inset in upper panel and targeting a GFP+ cell in lower panel). **(F)** Examples of membrane voltage responses to progressive current injections for control and *Pax6*cKO cortex (500ms square steps; hyperpolarizing step: -25 pA; depolarizing steps: rheobase and double rheobase). **(G)** Unsupervised hierarchical clustering analysis for S1 layer 5. Features used for clustering are listed in Table S5. Purple tones represent older pups (P8-P10). Green tones represent younger pups (P5-P7). Genotype bar indicates control cells (grey; n=54) and *Pax6*cKO cells (red; n=55). Cells from both genotypes were spread across the clusters with no segregation of *Pax6*cKO cells. Silhouette coefficient analysis suggested the optimal number of clusters was 2 (silhouette coefficient =0.26, k=2), which separated cells mainly by age (Source Data Table 4C). **(H)** Unsupervised hierarchical clustering analysis for S1 layers 2/3. Features used for clustering are listed in Table S5. Silhouette coefficient analysis suggested the optimal number of clusters was 2 (silhouette coefficient = 0.44, k=2). Clustering split the cells into two main branches, with one containing three cells all from *Pax6*cKO mice, but this separation might have occurred by chance (Barnard unconditional two-tailed test, p=0.09) (Source Data Table 4C). **(I)** S1 layer 2/3 GFP+ fast spiking cell; double rheobase current injection response.

**Figure S10:**
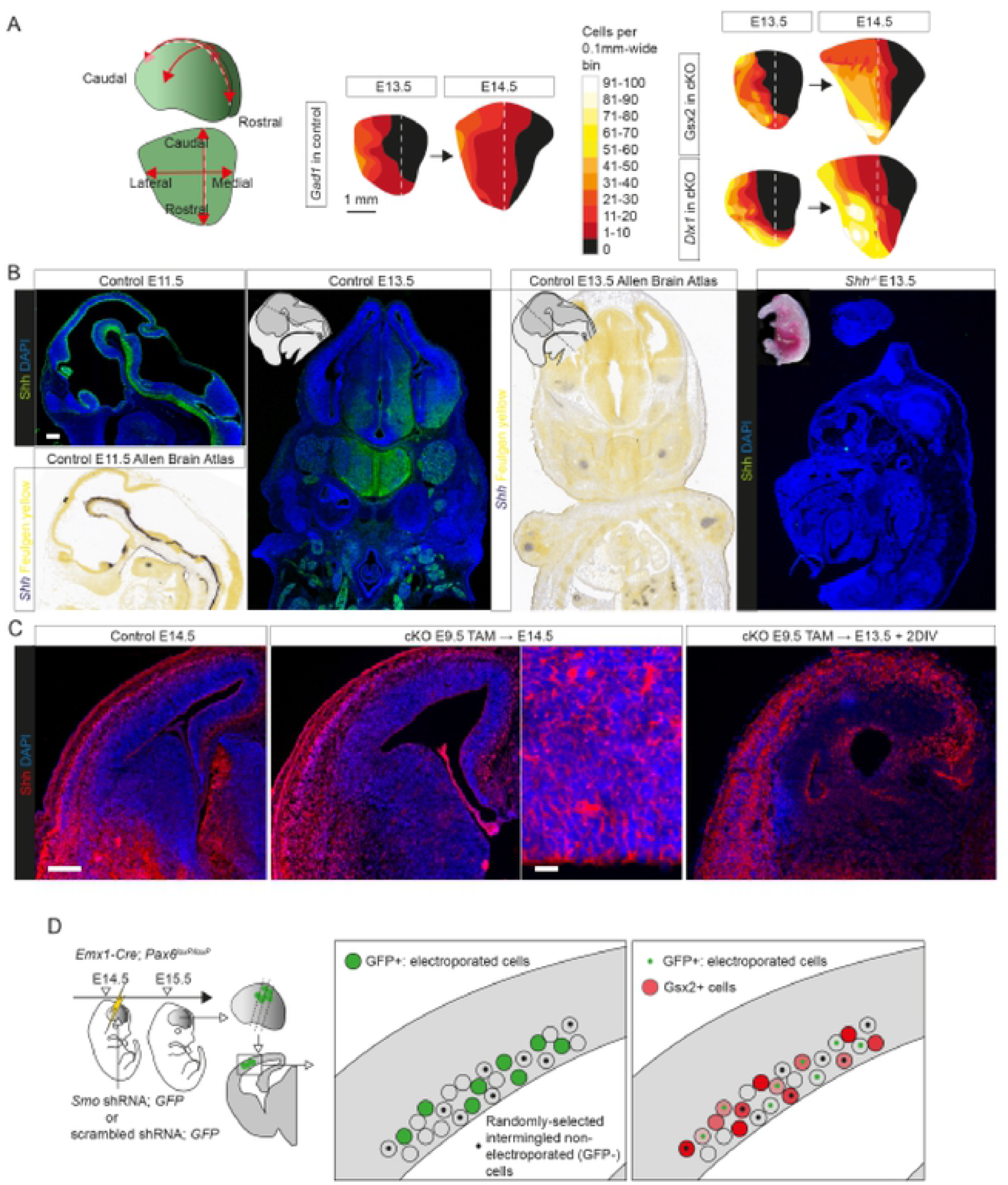
Expression and effects of Shh on eGC production. **(A)** Flattened surface views of the cortex showing the densities of *Gad1*+ cells at E13.5 and E14.5 in controls: *Gad1*+ cells spread in increasing numbers across the cortex from lateral to medial with a similar spatiotemporal pattern to the spread of *Gsx2* and *Dlx1* activation after tamoxifen at E9.5 (maps for Gsx2 and *Dlx1* reproduced from Fig. 6E). **(B)** Validation of the Shh antibody: comparison of staining patterns, in both neural and non-neural tissues (Bitgood and McMahon, 1995), with in situ hybridization patterns from the Allen Brain Atlas in E11.5 and E13.5 controls and lack of staining in E13.5 *Shh^-/-^* mutants (kindly provided by Laura Lettice and Bob Hill, Edinburgh University). Scale bar: 0.1mm. **(C)** Immunoreactivity for Shh in control telencephalon at E14.5 and in *Pax6* cKO telencephalon after tamoxifen (TAM) at E9.5 at E14.5 and after 2 days in vitro (DIV) from E13.5 (Fig. 10A). Scale bars: 0.1mm and 0.01mm. **(D)** Following the experimental procedure reproduced from Fig. 10K: constructs expressing *Smo* shRNA + GFP or scrambled shRNA + GFP were electroporated into the cortex of E14.5 *Pax6* cKO embryos made using *Emx1-Cre*; electroporated cells were analysed at E15.5. Blind to Gsx2 expression, we identified 80-100 GFP+ cells and a random selection of 80-100 intermingled GFP-cells in each of 3 embryos from 3 litters given *Smo* shRNA and each of 4 embryos from 3 litters given scrambled shRNA. The intensity of Gsx2 immunoreactivity was then measured in all of these cells and frequency distributions of intensities in electroporated versus non-electroporated cells were compared in brains that received *Smo* shRNA and in brains that received scrambled shRNA (results in Fig. 10M).

**Figure S11:**
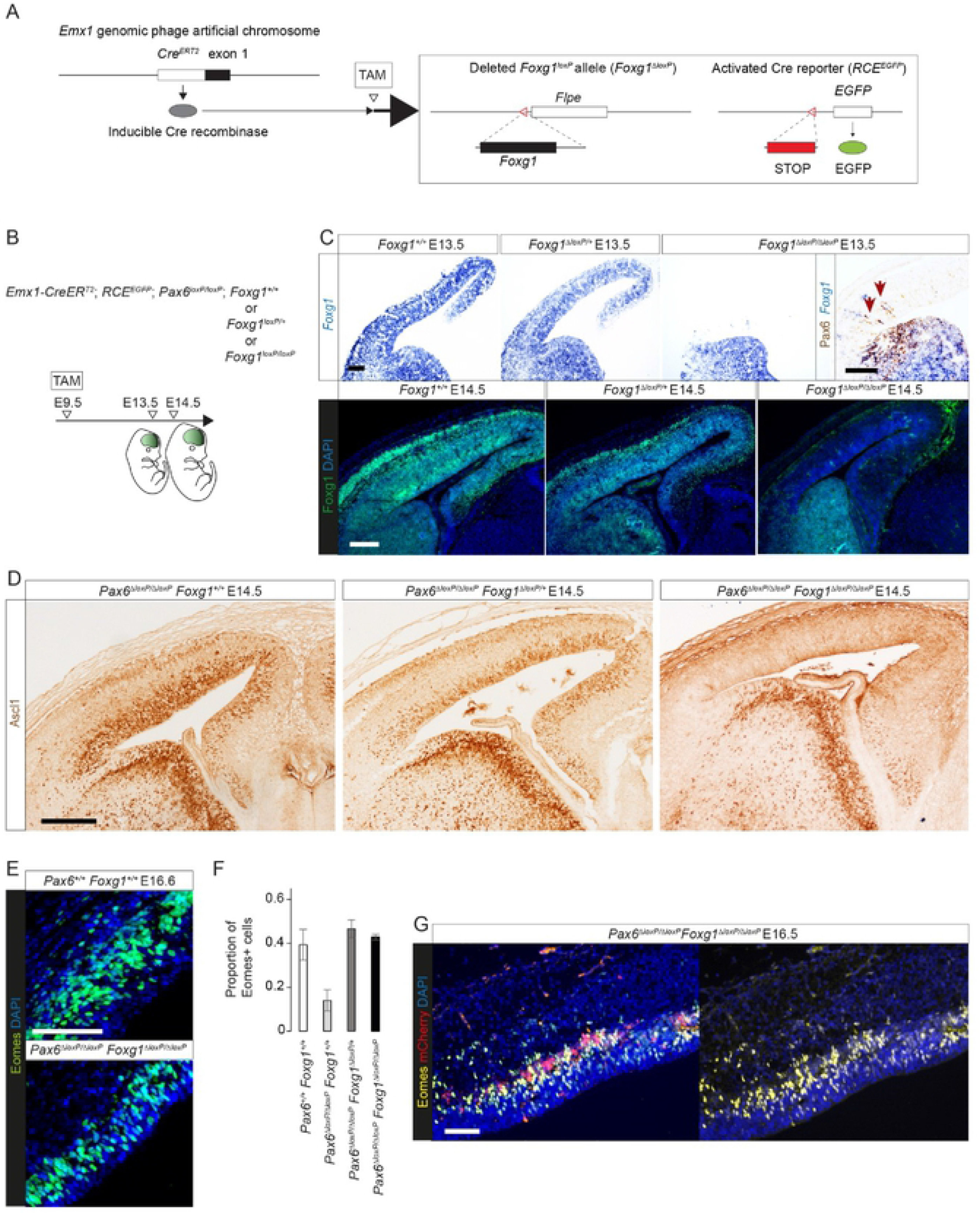
F*o*xg1 deletion and co-deletion with *Pax6*. **(A)** Deletion of *Foxg1*, here with *Emx1-Cre^ERT2^* and tamoxifen (TAM), removes its coding region (Miyoshi and Fishell, 2012) and activates the Cre reporter. **(B)** Tamoxifen (TAM) was administered at E9.5 to generate *Pax6* cKOs in which both, one or neither *Foxg1* allele(s) were also deleted; brains were analysed at E13.5 and E14.5. The *Pax6^loxP^* allele was shown in Fig. S1A. **(C)** In situ hybridizations for *Foxg1* and immunohistochemistry for Foxg1 and Pax6 following deletion of both, one or neither *Foxg1* allele(s) at E13.5 and E14.5. Arrows: a few cells remained undeleted and formed small clones expressing both Foxg1 and Pax6. Scale bars: 0.1mm. **(D)** Immunoreactivity for Ascl1 in E14.5 cortex from *Pax6* cKO embryos in which neither, one or both *Foxg1* allele(s) were deleted by tamoxifen at E9.5. Scale bars: 0.1mm. **(E)** Immunoreactivity for Eomes in E16.5 lateral cortex from control embryos and *Pax6 Foxg1* double KOs. Scale bar: 0.1mm. **(F)** Proportions of cells in the ventricular and subventricular zones of E16.5 lateral cortex expressing Eomes in control embryos and embryos of the three genotypes in (B) (averages ± sd of n=3; *Pax6* single cKO average was significantly lower than all others, p<0.05 in all comparisons; Student’s t-tests) (Source Data Table 6D). **(G)** Results of experiment in Fig. 11F: co-expression of Eomes and mCherry in a coronal section. Scale bar: 0.1mm.

**Figure S12:**
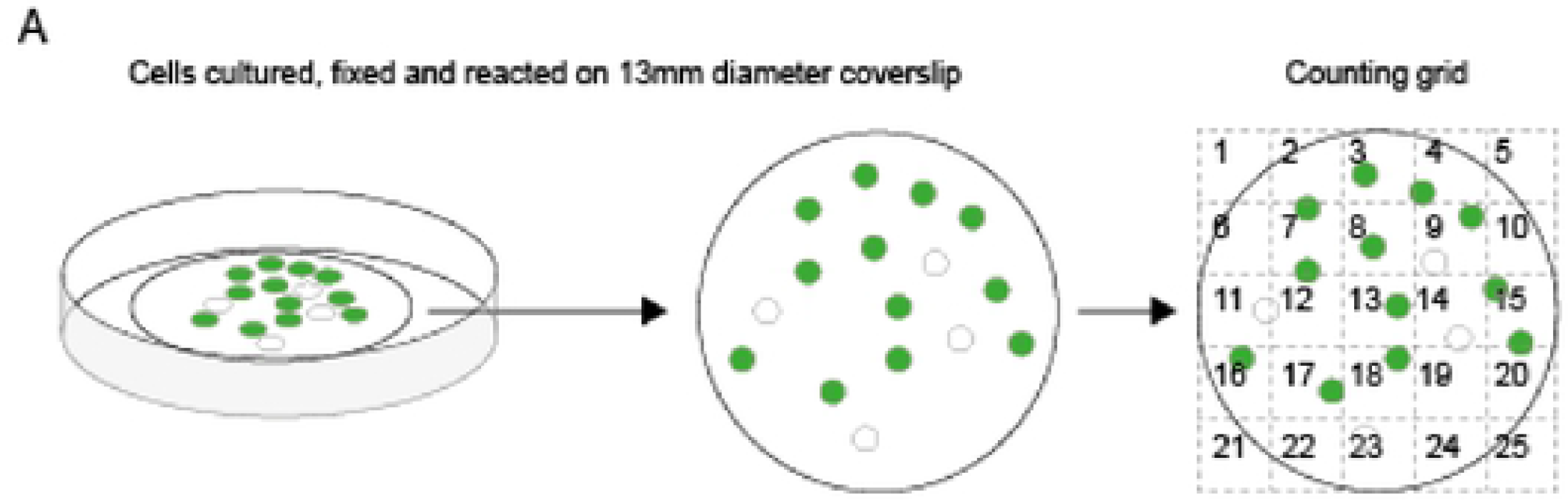
Quantification method for dissociated cultures. **(A)** Method for quantification of the effects of SAG on numbers of GFP+ cells expressing various markers: cells were cultured on coverslips and, after fixation and reaction, counting grids were used to sample 5 randomly selected areas from each coverslip. Several independent biological repeats were used for each condition (i.e. each concentration of SAG or vehicle alone, on *Pax6* cKO or control cells) (Fig. 12).

**Figure S13:**
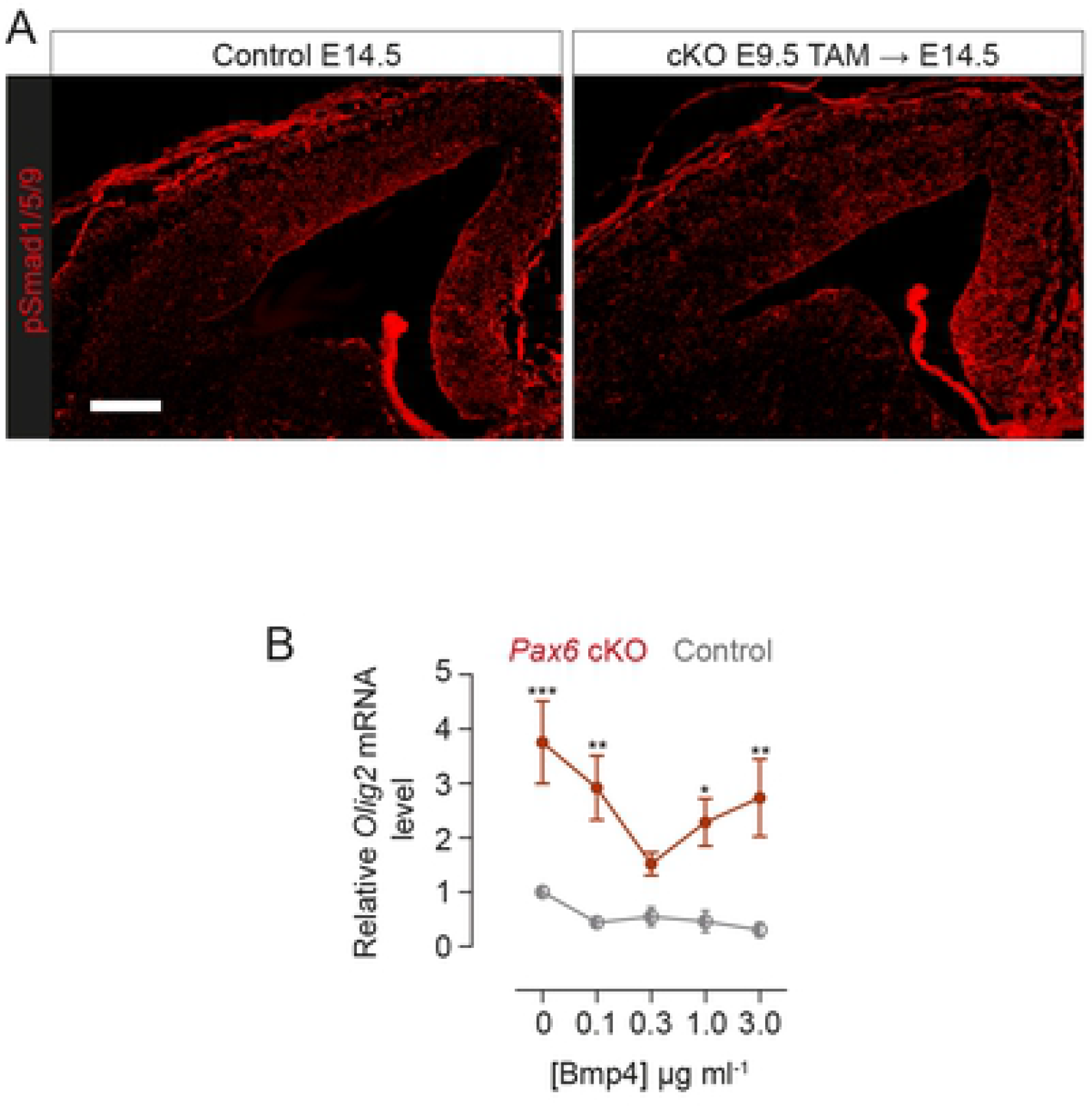
Phospho-Smad1/5/9 expression and Bmp4 effects on Olig2 expression in control and *Pax6* cKO cortex. **(A)** Immunoreactivity for phospho-Smad1/5/9 in E14.5 control and *Pax6* cKO cortex after tamoxifen^E9.5^. Scale bar: 0.1mm. **(B)** Concentration-response measured using qRT-PCR: *Olig2* levels (averages ±sem; values are relative to the average level in control cortex treated with 0 Bmp4) in control and *Pax6* cKO slices with increasing concentrations of Bmp4 (n=3 independent cultures at each concentration). Two-way ANOVA showed significant effects of genotype on *Olig2* (p<0.005), but no significant effect of Bmp4 concentration and no significant interaction effect. Differences between genotypes at each Bmp4 concentration were tested with Bonferroni’s method for comparison of means (*, p<0.05; **, p<0.01; ***, p<0.005) (Source Data Table 8).

## Supplementary Tables

Table S1: Results of bulk RNAseq: log_2_ fold changes of genes that were significantly differentially expressed between control and *Pax6* cKO cortex (adjusted p, or padj, <0.05) rostrally and caudally at E12.5 and E13.5.

Table S2: Significantly enriched gene ontology (GO) terms obtained by passing through DAVID v6.8 all functionally annotated genes with a LFC in expression ≥ 1 (for upregulated genes) or ≤ -1 (for downregulated genes) in at least one of the four combinations of age and region studied.

Table S3: Citations providing evidence on the normal sites of expression of genes that were upregulated in *Pax6* cKO RNAseq datasets.

Table S4: Genes whose expression levels were the most different between RGPs and aRGPs (average LFCs >0.3 or <-0.3) and GO terms obtained by passing this list through DAVID.

Table S5: Intrinsic properties of primary somatosensory cortex (S1) layer 5 cells at P5-7 and P8-P10 and layer 2/3 cells at P10-13. Values are means ± 95% confidence intervals.

**Source Data** Tables 1-8: Data referred to in the text and Figure legends.

## REFERENCES

Alcamo, E. A., L. Chirivella, M. Dautzenberg, G. Dobreva, I. Fariñas, R. Grosschedl and S. K. McConnell (2008). “Satb2 regulates callosal projection neuron identity in the developing cerebral cortex.” Neuron 57(3): 364–377.

Anderson, S. A., D. D. Eisenstat, L. Shi and J. L. Rubenstein (1997). “Interneuron migration from basal forebrain to neocortex: dependence on Dlx genes.” Science 278(5337): 474–476.

Anderson, S. A., C. E. Kaznowski, C. Horn, J. L. Rubenstein and S. K. McConnell (2002). “Distinct origins of neocortical projection neurons and interneurons in vivo.” Cereb Cortex 12(7): 702–709.

Anderson, S. A., O. Marín, C. Horn, K. Jennings and J. L. Rubenstein (2001). “Distinct cortical migrations from the medial and lateral ganglionic eminences.” Development 128(3): 353–363.

Andjelic, S., T. Gallopin, B. Cauli, E. L. Hill, L. Roux, S. Badr, E. Hu, G. Tamás and B. Lambolez (2009). “Glutamatergic nonpyramidal neurons from neocortical layer VI and their comparison with pyramidal and spiny stellate neurons.” J Neurophysiol 101(2): 641–654.

Angevine, J. B., Jr. and R. L. Sidman (1961). “Autoradiographic study of cell migration during histogenesis of cerebral cortex in the mouse.” Nature 192: 766–768.

Bai, C. B., W. Auerbach, J. S. Lee, D. Stephen and A. L. Joyner (2002). “Gli2, but not Gli1, is required for initial Shh signaling and ectopic activation of the Shh pathway.” Development 129(20): 4753–4761.

Bai, C. B., D. Stephen and A. L. Joyner (2004). “All mouse ventral spinal cord patterning by hedgehog is Gli dependent and involves an activator function of Gli3.” Dev Cell 6(1): 103–115.

Barber, M., T. Di Meglio, W. D. Andrews, L. R. Hernández-Miranda, F. Murakami, A. Chédotal and J. G. Parnavelas (2009). “The role of Robo3 in the development of cortical interneurons.” Cereb Cortex 19 Suppl 1(Suppl 1): i22-31.

Baudoin, J. P., L. Viou, P. S. Launay, C. Luccardini, S. Espeso Gil, V. Kiyasova, T. Irinopoulou, C. Alvarez, J. P. Rio, T. Boudier, J. P. Lechaire, N. Kessaris, N. Spassky and C. Métin (2012). “Tangentially migrating neurons assemble a primary cilium that promotes their reorientation to the cortical plate.” Neuron 76(6): 1108–1122.

Bean, B. P. (2007). “The action potential in mammalian central neurons.” Nat Rev Neurosci 8(6): 451–465.

Bergen, V., M. Lange, S. Peidli, F. A. Wolf and F. J. Theis (2020). “Generalizing RNA velocity to transient cell states through dynamical modeling.” Nat Biotechnol 38(12): 1408–1414.

Bitgood, M. J. and A. P. McMahon (1995). “Hedgehog and Bmp genes are coexpressed at many diverse sites of cell-cell interaction in the mouse embryo.” Dev Biol 172(1): 126–138.

Britanova, O., C. de Juan Romero, A. Cheung, K. Y. Kwan, M. Schwark, A. Gyorgy, T. Vogel, S. Akopov, M. Mitkovski, D. Agoston, N. Sestan, Z. Molnár and V. Tarabykin (2008). “Satb2 is a postmitotic determinant for upper-layer neuron specification in the neocortex.” Neuron 57(3): 378–392.

Bulchand, S., L. Subramanian and S. Tole (2003). “Dynamic spatiotemporal expression of LIM genes and cofactors in the embryonic and postnatal cerebral cortex.” Dev Dyn 226(3): 460–469.

Bulfone, A., S. M. Smiga, K. Shimamura, A. Peterson, L. Puelles and J. L. Rubenstein (1995). “T-brain-1: a homolog of Brachyury whose expression defines molecularly distinct domains within the cerebral cortex.” Neuron 15(1): 63–78.

Bulstrode, H., E. Johnstone, M. A. Marques-Torrejon, K. M. Ferguson, R. B. Bressan, C. Blin, V. Grant, S. Gogolok, E. Gangoso, S. Gagrica, C. Ender, V. Fotaki, D. Sproul, P. Bertone and S. M. Pollard (2017). “Elevated FOXG1 and SOX2 in glioblastoma enforces neural stem cell identity through transcriptional control of cell cycle and epigenetic regulators.” Genes Dev 31(8): 757–773.

Butt, S. J., V. H. Sousa, M. V. Fuccillo, J. Hjerling-Leffler, G. Miyoshi, S. Kimura and G. Fishell (2008). “The requirement of Nkx2-1 in the temporal specification of cortical interneuron subtypes.” Neuron 59(5): 722–732.

Caballero, I. M., M. N. Manuel, M. Molinek, I. Quintana-Urzainqui, D. Mi, T. Shimogori and D. J. Price (2014). “Cell-autonomous repression of Shh by transcription factor Pax6 regulates diencephalic patterning by controlling the central diencephalic organizer.” Cell Rep 8(5): 1405–1418.

Callaerts, P., G. Halder and W. J. Gehring (1997). “PAX-6 in development and evolution.” Annu Rev Neurosci 20: 483–532.

Carić, D., D. Gooday, R. E. Hill, S. K. McConnell and D. J. Price (1997). “Determination of the migratory capacity of embryonic cortical cells lacking the transcription factor Pax-6.” Development 124(24): 5087–5096.

Caronia, G., J. Wilcoxon, P. Feldman and E. A. Grove (2010). “Bone morphogenetic protein signaling in the developing telencephalon controls formation of the hippocampal dentate gyrus and modifies fear-related behavior.” J Neurosci 30(18): 6291–6301.

Castro, D. S., B. Martynoga, C. Parras, V. Ramesh, E. Pacary, C. Johnston, D. Drechsel, M. Lebel-Potter, L. G. Garcia, C. Hunt, D. Dolle, A. Bithell, L. Ettwiller, N. Buckley and F. Guillemot (2011). “A novel function of the proneural factor Ascl1 in progenitor proliferation identified by genome-wide characterization of its targets.” Genes Dev 25(9): 930–945.

Castro, D. S., D. Skowronska-Krawczyk, O. Armant, I. J. Donaldson, C. Parras, C. Hunt, J. A. Critchley, L. Nguyen, A. Gossler, B. Göttgens, J. M. Matter and F. Guillemot (2006). “Proneural bHLH and Brn proteins coregulate a neurogenic program through cooperative binding to a conserved DNA motif.” Dev Cell 11(6): 831–844.

Caviness, V. S., Jr. (1982). “Neocortical histogenesis in normal and reeler mice: a developmental study based upon [3H]thymidine autoradiography.” Brain Res 256(3): 293–302.

Chau, K. F., M. W. Springel, K. G. Broadbelt, H. Y. Park, S. Topal, M. P. Lun, H. Mullan, T. Maynard, H. Steen, A. S. LaMantia and M. K. Lehtinen (2015). “Progressive Differentiation and Instructive Capacities of Amniotic Fluid and Cerebrospinal Fluid Proteomes following Neural Tube Closure.” Dev Cell 35(6): 789–802.

Chen, J. K., J. Taipale, K. E. Young, T. Maiti and P. A. Beachy (2002). “Small molecule modulation of Smoothened activity.” Proc Natl Acad Sci U S A 99(22): 14071–14076.

Cheng, X., C. M. Hsu, D. S. Currle, J. S. Hu, A. J. Barkovich and E. S. Monuki (2006). “Central roles of the roof plate in telencephalic development and holoprosencephaly.” J Neurosci 26(29): 7640–7649.

Chi, L., B. Fan, D. Feng, Z. Chen, Z. Liu, Y. Hui, X. Xu, L. Ma, Y. Fang, Q. Zhang, G. Jin, L. Liu, F. Guan and X. Zhang (2017). “The Dorsoventral Patterning of Human Forebrain Follows an Activation/Transformation Model.” Cereb Cortex 27(5): 2941–2954.

Chiang, C., Y. Litingtung, E. Lee, K. E. Young, J. L. Corden, H. Westphal and P. A. Beachy (1996). “Cyclopia and defective axial patterning in mice lacking Sonic hedgehog gene function.” Nature 383(6599): 407–413.

Collinson, J. M., R. E. Hill and J. D. West (2000). “Different roles for Pax6 in the optic vesicle and facial epithelium mediate early morphogenesis of the murine eye.” Development 127(5): 945–956.

Connors, B. W. and M. J. Gutnick (1990). “Intrinsic firing patterns of diverse neocortical neurons.” Trends Neurosci 13(3): 99–104.

Corbin, J. G., N. Gaiano, R. P. Machold, A. Langston and G. Fishell (2000). “The Gsh2 homeodomain gene controls multiple aspects of telencephalic development.” Development 127(23): 5007–5020.

Dahmane, N., P. Sánchez, Y. Gitton, V. Palma, T. Sun, M. Beyna, H. Weiner and A. Ruiz i Altaba (2001). “The Sonic Hedgehog-Gli pathway regulates dorsal brain growth and tumorigenesis.” Development 128(24): 5201–5212.

Danesin, C., J. N. Peres, M. Johansson, V. Snowden, A. Cording, N. Papalopulu and C. Houart (2009). “Integration of telencephalic Wnt and hedgehog signaling center activities by Foxg1.” Dev Cell 16(4): 576–587.

Danjo, T., M. Eiraku, K. Muguruma, K. Watanabe, M. Kawada, Y. Yanagawa, J. L. Rubenstein and Y. Sasai (2011). “Subregional specification of embryonic stem cell-derived ventral telencephalic tissues by timed and combinatory treatment with extrinsic signals.” J Neurosci 31(5): 1919–1933.

Daw, M. I., M. C. Ashby and J. T. Isaac (2007). “Coordinated developmental recruitment of latent fast spiking interneurons in layer IV barrel cortex.” Nat Neurosci 10(4): 453–461.

del Rio, J. A. and E. Soriano (1989). “Immunocytochemical detection of 5’-bromodeoxyuridine incorporation in the central nervous system of the mouse.” Brain Res Dev Brain Res 49(2): 311–317.

Delmotte, Q., M. Hamze, I. Medina, E. Buhler, J. Zhang, Y. H. Belgacem and C. Porcher (2020). “Smoothened receptor signaling regulates the developmental shift of GABA polarity in rat somatosensory cortex.” J Cell Sci 133(20).

Destexhe, A., M. Rudolph and D. Paré (2003). “The high-conductance state of neocortical neurons in vivo.” Nat Rev Neurosci 4(9): 739–751.

Doan, L. T., A. L. Javier, N. M. Furr, K. L. Nguyen, K. W. Cho and E. S. Monuki (2012). “A Bmp reporter with ultrasensitive characteristics reveals that high Bmp signaling is not required for cortical hem fate.” PLoS One 7(9): e44009.

Echelard, Y., D. J. Epstein, B. St-Jacques, L. Shen, J. Mohler, J. A. McMahon and A. P. McMahon (1993). “Sonic hedgehog, a member of a family of putative signaling molecules, is implicated in the regulation of CNS polarity.” Cell 75(7): 1417–1430.

Elliott, A. A. and J. R. Elliott (1993). “Characterization of TTX-sensitive and TTX-resistant sodium currents in small cells from adult rat dorsal root ganglia.” J Physiol 463: 39–56.

Endoh-Yamagami, S., M. Evangelista, D. Wilson, X. Wen, J. W. Theunissen, K. Phamluong, M. Davis, S. J. Scales, M. J. Solloway, F. J. de Sauvage and A. S. Peterson (2009). “The mammalian Cos2 homolog Kif7 plays an essential role in modulating Hh signal transduction during development.” Curr Biol 19(15): 1320–1326.

Engelkamp, D., P. Rashbass, A. Seawright and V. van Heyningen (1999). “Role of Pax6 in development of the cerebellar system.” Development 126(16): 3585–3596.

Englund, C., A. Fink, C. Lau, D. Pham, R. A. Daza, A. Bulfone, T. Kowalczyk and R. F. Hevner (2005). “Pax6, Tbr2, and Tbr1 are expressed sequentially by radial glia, intermediate progenitor cells, and postmitotic neurons in developing neocortex.” J Neurosci 25(1): 247–251.

Ericson, J., P. Rashbass, A. Schedl, S. Brenner-Morton, A. Kawakami, V. van Heyningen, T. M. Jessell and J. Briscoe (1997). “Pax6 controls progenitor cell identity and neuronal fate in response to graded Shh signaling.” Cell 90(1): 169–180.

Ferrell, J. E., Jr. (2012). “Bistability, bifurcations, and Waddington’s epigenetic landscape.” Curr Biol 22(11): R458–466.

Flames, N., R. Pla, D. M. Gelman, J. L. Rubenstein, L. Puelles and O. Marín (2007). “Delineation of multiple subpallial progenitor domains by the combinatorial expression of transcriptional codes.” J Neurosci 27(36): 9682–9695.

Fode, C., Q. Ma, S. Casarosa, S. L. Ang, D. J. Anderson and F. Guillemot (2000). “A role for neural determination genes in specifying the dorsoventral identity of telencephalic neurons.” Genes Dev 14(1): 67–80.

Furuta, Y., D. W. Piston and B. L. Hogan (1997). “Bone morphogenetic proteins (BMPs) as regulators of dorsal forebrain development.” Development 124(11): 2203–2212.

Gaiano, N., J. D. Kohtz, D. H. Turnbull and G. Fishell (1999). “A method for rapid gain-of-function studies in the mouse embryonic nervous system.” Nat Neurosci 2(9): 812–819.

Gao, P., M. P. Postiglione, T. G. Krieger, L. Hernandez, C. Wang, Z. Han, C. Streicher, E. Papusheva, R. Insolera, K. Chugh, O. Kodish, K. Huang, B. D. Simons, L. Luo, S. Hippenmeyer and S. H. Shi (2014). “Deterministic progenitor behavior and unitary production of neurons in the neocortex.” Cell 159(4): 775–788.

Georgala, P. A., M. Manuel and D. J. Price (2011). “The generation of superficial cortical layers is regulated by levels of the transcription factor Pax6.” Cereb Cortex 21(1): 81–94.

Gillies, K. and D. J. Price (1993). “The fates of cells in the developing cerebral cortex of normal and methylazoxymethanol acetate-lesioned mice.” Eur J Neurosci 5(1): 73–84.

Goodrich, L. V., R. L. Johnson, L. Milenkovic, J. A. McMahon and M. P. Scott (1996). “Conservation of the hedgehog/patched signaling pathway from flies to mice: induction of a mouse patched gene by Hedgehog.” Genes Dev 10(3): 301–312.

Gorski, J. A., T. Talley, M. Qiu, L. Puelles, J. L. Rubenstein and K. R. Jones (2002). “Cortical excitatory neurons and glia, but not GABAergic neurons, are produced in the Emx1-expressing lineage.” J Neurosci 22(15): 6309–6314.

Groves, I., M. Placzek and A. G. Fletcher (2020). “Of mitogens and morphogens: modelling Sonic Hedgehog mechanisms in vertebrate development.” Philos Trans R Soc Lond B Biol Sci 375(1809): 20190660.

Gulacsi, A. and L. Lillien (2003). “Sonic hedgehog and bone morphogenetic protein regulate interneuron development from dorsal telencephalic progenitors in vitro.” J Neurosci 23(30): 9862–9872.

Gunhaga, L., T. M. Jessell and T. Edlund (2000). “Sonic hedgehog signaling at gastrula stages specifies ventral telencephalic cells in the chick embryo.” Development 127(15): 3283–3293.

Guo, T., G. Liu, H. Du, Y. Wen, S. Wei, Z. Li, G. Tao, Z. Shang, X. Song, Z. Zhang, Z. Xu, Y. You, B. Chen, J. L. Rubenstein and Z. Yang (2019). “Dlx1/2 are Central and Essential Components in the Transcriptional Code for Generating Olfactory Bulb Interneurons.” Cereb Cortex 29(11): 4831–4849.

Haubensak, W., A. Attardo, W. Denk and W. B. Huttner (2004). “Neurons arise in the basal neuroepithelium of the early mammalian telencephalon: a major site of neurogenesis.” Proc Natl Acad Sci U S A 101(9): 3196–3201.

Hayashi, S. and A. P. McMahon (2002). “Efficient recombination in diverse tissues by a tamoxifen-inducible form of Cre: a tool for temporally regulated gene activation/inactivation in the mouse.” Dev Biol 244(2): 305–318.

Hegarty, S. V., G. W. O’Keeffe and A. M. Sullivan (2013). “BMP-Smad 1/5/8 signalling in the development of the nervous system.” Prog Neurobiol 109: 28–41.

Heine, V. M., A. Griveau, C. Chapin, P. L. Ballard, J. K. Chen and D. H. Rowitch (2011). “A small-molecule smoothened agonist prevents glucocorticoid-induced neonatal cerebellar injury.” Sci Transl Med 3(105): 105ra104.

Heins, N., P. Malatesta, F. Cecconi, M. Nakafuku, K. L. Tucker, M. A. Hack, P. Chapouton, Y. A. Barde and M. Götz (2002). “Glial cells generate neurons: the role of the transcription factor Pax6.” Nat Neurosci 5(4): 308–315.

Herdegen, T. and J. D. Leah (1998). “Inducible and constitutive transcription factors in the mammalian nervous system: control of gene expression by Jun, Fos and Krox, and CREB/ATF proteins.” Brain Res Brain Res Rev 28(3): 370–490.

Hevner, R. F., L. Shi, N. Justice, Y. Hsueh, M. Sheng, S. Smiga, A. Bulfone, A. M. Goffinet, A. T. Campagnoni and J. L. Rubenstein (2001). “Tbr1 regulates differentiation of the preplate and layer 6.” Neuron 29(2): 353–366.

Hill, R. E., J. Favor, B. L. Hogan, C. C. Ton, G. F. Saunders, I. M. Hanson, J. Prosser, T. Jordan, N. D. Hastie and V. van Heyningen (1991). “Mouse small eye results from mutations in a paired-like homeobox-containing gene.” Nature 354(6354): 522–525.

Holm, P. C., M. T. Mader, N. Haubst, A. Wizenmann, M. Sigvardsson and M. Götz (2007). “Loss-and gain-of-function analyses reveal targets of Pax6 in the developing mouse telencephalon.” Mol Cell Neurosci 34(1): 99–119.

Hou, S., W. L. Ho, L. Wang, B. Kuo, J. Y. Park and Y. G. Han (2021). “Biphasic Roles of Hedgehog Signaling in the Production and Self-Renewal of Outer Radial Glia in the Ferret Cerebral Cortex.” Cereb Cortex 31(10): 4730–4741.

Hu, J. S., L. T. Doan, D. S. Currle, M. Paff, J. Y. Rheem, R. Schreyer, B. Robert and E. S. Monuki (2008). “Border formation in a Bmp gradient reduced to single dissociated cells.” Proc Natl Acad Sci U S A 105(9): 3398–3403.

Huang da, W., B. T. Sherman and R. A. Lempicki (2009). “Systematic and integrative analysis of large gene lists using DAVID bioinformatics resources.” Nat Protoc 4(1): 44–57.

Huang da, W., B. T. Sherman, X. Zheng, J. Yang, T. Imamichi, R. Stephens and R. A. Lempicki (2009). “Extracting biological meaning from large gene lists with DAVID.” Curr Protoc Bioinformatics Chapter 13: Unit 13.11.

Huang, S., Y. P. Guo, G. May and T. Enver (2007). “Bifurcation dynamics in lineage-commitment in bipotent progenitor cells.” Dev Biol 305(2): 695–713.

Iacopetti, P., M. Michelini, I. Stuckmann, B. Oback, E. Aaku-Saraste and W. B. Huttner (1999). “Expression of the antiproliferative gene TIS21 at the onset of neurogenesis identifies single neuroepithelial cells that switch from proliferative to neuron-generating division.” Proc Natl Acad Sci U S A 96(8): 4639–4644.

Kalyani, A. J., D. Piper, T. Mujtaba, M. T. Lucero and M. S. Rao (1998). “Spinal cord neuronal precursors generate multiple neuronal phenotypes in culture.” J Neurosci 18(19): 7856–7868.

Kaplan, E. S., K. A. Ramos-Laguna, A. B. Mihalas, R. A. M. Daza and R. F. Hevner (2017). “Neocortical Sox9+ radial glia generate glutamatergic neurons for all layers, but lack discernible evidence of early laminar fate restriction.” Neural Dev 12(1): 14.

Karin, M., Z. Liu and E. Zandi (1997). “AP-1 function and regulation.” Curr Opin Cell Biol 9(2): 240–246.

Kessaris, N., M. Fogarty, P. Iannarelli, M. Grist, M. Wegner and W. D. Richardson (2006). “Competing waves of oligodendrocytes in the forebrain and postnatal elimination of an embryonic lineage.” Nat Neurosci 9(2): 173–179.

Kikkawa, T., T. Obayashi, M. Takahashi, U. Fukuzaki-Dohi, K. Numayama-Tsuruta and N. Osumi (2013). “Dmrta1 regulates proneural gene expression downstream of Pax6 in the mammalian telencephalon.” Genes Cells 18(8): 636–649.

Komada, M., H. Saitsu, M. Kinboshi, T. Miura, K. Shiota and M. Ishibashi (2008). “Hedgehog signaling is involved in development of the neocortex.” Development 135(16): 2717–2727.

Kovach, C., R. Dixit, S. Li, P. Mattar, G. Wilkinson, G. E. Elsen, D. M. Kurrasch, R. F. Hevner and C. Schuurmans (2013). “Neurog2 simultaneously activates and represses alternative gene expression programs in the developing neocortex.” Cereb Cortex 23(8): 1884–1900.

Kowalczyk, M. S., I. Tirosh, D. Heckl, T. N. Rao, A. Dixit, B. J. Haas, R. K. Schneider, A. J. Wagers, B. L. Ebert and A. Regev (2015). “Single-cell RNA-seq reveals changes in cell cycle and differentiation programs upon aging of hematopoietic stem cells.” Genome Res 25(12): 1860–1872.

Kroll, T. T. and D. D. O’Leary (2005). “Ventralized dorsal telencephalic progenitors in Pax6 mutant mice generate GABA interneurons of a lateral ganglionic eminence fate.” Proc Natl Acad Sci U S A 102(20): 7374–7379.

La Manno, G., R. Soldatov, A. Zeisel, E. Braun, H. Hochgerner, V. Petukhov, K. Lidschreiber, M. E. Kastriti, P. Lönnerberg, A. Furlan, J. Fan, L. E. Borm, Z. Liu, D. van Bruggen, J. Guo, X. He, R. Barker, E. Sundström, G. Castelo-Branco, P. Cramer, I. Adameyko, S. Linnarsson and P. V. Kharchenko (2018). “RNA velocity of single cells.” Nature 560(7719): 494–498.

Lai, T., D. Jabaudon, B. J. Molyneaux, E. Azim, P. Arlotta, J. R. Menezes and J. D. Macklis (2008). “SOX5 controls the sequential generation of distinct corticofugal neuron subtypes.” Neuron 57(2): 232–247.

Le, T. N., G. Du, M. Fonseca, Q. P. Zhou, J. T. Wigle and D. D. Eisenstat (2007). “Dlx homeobox genes promote cortical interneuron migration from the basal forebrain by direct repression of the semaphorin receptor neuropilin-2.” J Biol Chem 282(26): 19071–19081.

Le, T. N., Q. P. Zhou, I. Cobos, S. Zhang, J. Zagozewski, S. Japoni, J. Vriend, T. Parkinson, G. Du, J. L. Rubenstein and D. D. Eisenstat (2017). “GABAergic Interneuron Differentiation in the Basal Forebrain Is Mediated through Direct Regulation of Glutamic Acid Decarboxylase Isoforms by Dlx Homeobox Transcription Factors.” J Neurosci 37(36): 8816–8829.

Lee, J., K. A. Platt, P. Censullo and A. Ruiz i Altaba (1997). “Gli1 is a target of Sonic hedgehog that induces ventral neural tube development.” Development 124(13): 2537–2552.

Lehtinen, M. K., M. W. Zappaterra, X. Chen, Y. J. Yang, A. D. Hill, M. Lun, T. Maynard, D. Gonzalez, S. Kim, P. Ye, A. J. D’Ercole, E. T. Wong, A. S. LaMantia and C. A. Walsh (2011). “The cerebrospinal fluid provides a proliferative niche for neural progenitor cells.” Neuron 69(5): 893–905.

Lendahl, U., L. B. Zimmerman and R. D. McKay (1990). “CNS stem cells express a new class of intermediate filament protein.” Cell 60(4): 585–595.

Leone, D. P., K. Srinivasan, B. Chen, E. Alcamo and S. K. McConnell (2008). “The determination of projection neuron identity in the developing cerebral cortex.” Curr Opin Neurobiol 18(1): 28–35.

Liem, K. F., Jr., M. He, P. J. Ocbina and K. V. Anderson (2009). “Mouse Kif7/Costal2 is a cilia-associated protein that regulates Sonic hedgehog signaling.” Proc Natl Acad Sci U S A 106(32): 13377–13382.

Lim, L., D. Mi, A. Llorca and O. Marín (2018). “Development and Functional Diversification of Cortical Interneurons.” Neuron 100(2): 294–313.

Lin, Y. L., S. D. Persaud, J. Nhieu and L. N. Wei (2017). “Cellular Retinoic Acid-Binding Protein 1 Modulates Stem Cell Proliferation to Affect Learning and Memory in Male Mice.” Endocrinology 158(9): 3004–3014.

Long, J. E., C. Swan, W. S. Liang, I. Cobos, G. B. Potter and J. L. Rubenstein (2009). “Dlx1&2 and Mash1 transcription factors control striatal patterning and differentiation through parallel and overlapping pathways.” J Comp Neurol 512(4): 556–572.

LoTurco, J. J., M. G. Blanton and A. R. Kriegstein (1991). “Initial expression and endogenous activation of NMDA channels in early neocortical development.” J Neurosci 11(3): 792–799.

Love, M. I., W. Huber and S. Anders (2014). “Moderated estimation of fold change and dispersion for RNA-seq data with DESeq2.” Genome Biol 15(12): 550.

Luan, Q., Q. Chen and M. Friedrich (2014). “The Pax6 genes eyeless and twin of eyeless are required for global patterning of the ocular segment in the Tribolium embryo.” Dev Biol 394(2): 367–381.

Luhmann, H. J., R. A. Reiprich, I. Hanganu and W. Kilb (2000). “Cellular physiology of the neonatal rat cerebral cortex: intrinsic membrane properties, sodium and calcium currents.” J Neurosci Res 62(4): 574–584.

Macdonald, R., K. A. Barth, Q. Xu, N. Holder, I. Mikkola and S. W. Wilson (1995). “Midline signalling is required for Pax gene regulation and patterning of the eyes.” Development 121(10): 3267–3278.

Manuel, M., P. A. Georgala, C. B. Carr, S. Chanas, D. A. Kleinjan, B. Martynoga, J. O. Mason, M. Molinek, J. Pinson, T. Pratt, J. C. Quinn, T. I. Simpson, D. A. Tyas, V. van Heyningen, J. D. West and D. J. Price (2007). “Controlled overexpression of Pax6 in vivo negatively autoregulates the Pax6 locus, causing cell-autonomous defects of late cortical progenitor proliferation with little effect on cortical arealization.” Development 134(3): 545–555.

Manuel, M., B. Martynoga, T. Yu, J. D. West, J. O. Mason and D. J. Price (2010). “The transcription factor Foxg1 regulates the competence of telencephalic cells to adopt subpallial fates in mice.” Development 137(3): 487–497.

Marigo, V. and C. J. Tabin (1996). “Regulation of patched by sonic hedgehog in the developing neural tube.” Proc Natl Acad Sci U S A 93(18): 9346–9351.

Markram, H., M. Toledo-Rodriguez, Y. Wang, A. Gupta, G. Silberberg and C. Wu (2004). “Interneurons of the neocortical inhibitory system.” Nat Rev Neurosci 5(10): 793–807.

Martí, E., D. A. Bumcrot, R. Takada and A. P. McMahon (1995). “Requirement of 19K form of Sonic hedgehog for induction of distinct ventral cell types in CNS explants.” Nature 375(6529): 322–325.

Mastick, G. S., N. M. Davis, G. L. Andrew and S. S. Easter, Jr. (1997). “Pax-6 functions in boundary formation and axon guidance in the embryonic mouse forebrain.” Development 124(10): 1985–1997.

Matise, M. P. and H. Wang (2011). “Sonic hedgehog signaling in the developing CNS where it has been and where it is going.” Curr Top Dev Biol 97: 75–117.

Mayer, C., C. Hafemeister, R. C. Bandler, R. Machold, R. Batista Brito, X. Jaglin, K. Allaway, A. Butler, G. Fishell and R. Satija (2018). “Developmental diversification of cortical inhibitory interneurons.” Nature 555(7697): 457–462.

Mi, D., C. B. Carr, P. A. Georgala, Y. T. Huang, M. N. Manuel, E. Jeanes, E. Niisato, S. N. Sansom, F. J. Livesey, T. Theil, K. Hasenpusch-Theil, T. I. Simpson, J. O. Mason and D. J. Price (2013). “Pax6 exerts regional control of cortical progenitor proliferation via direct repression of Cdk6 and hypophosphorylation of pRb.” Neuron 78(2): 269–284.

Mi, D., M. Manuel, Y. T. Huang, J. O. Mason and D. J. Price (2018). “Pax6 Lengthens G1 Phase and Decreases Oscillating Cdk6 Levels in Murine Embryonic Cortical Progenitors.” Front Cell Neurosci 12: 419.

Mihalas, A. B. and R. F. Hevner (2018). “Clonal analysis reveals laminar fate multipotency and daughter cell apoptosis of mouse cortical intermediate progenitors.” Development 145(17).

Miyoshi, G., S. J. Butt, H. Takebayashi and G. Fishell (2007). “Physiologically distinct temporal cohorts of cortical interneurons arise from telencephalic Olig2-expressing precursors.” J Neurosci 27(29): 7786–7798.

Miyoshi, G. and G. Fishell (2012). “Dynamic FoxG1 expression coordinates the integration of multipolar pyramidal neuron precursors into the cortical plate.” Neuron 74(6): 1045–1058.

Miyoshi, G., J. Hjerling-Leffler, T. Karayannis, V. H. Sousa, S. J. Butt, J. Battiste, J. E. Johnson, R. P. Machold and G. Fishell (2010). “Genetic fate mapping reveals that the caudal ganglionic eminence produces a large and diverse population of superficial cortical interneurons.” J Neurosci 30(5): 1582–1594.

Morinello, E., M. Pignatello, L. Villabruna, P. Goelzer and H. Bürgin (2014). “Embryofetal development study of vismodegib, a hedgehog pathway inhibitor, in rats.” Birth Defects Res B Dev Reprod Toxicol 101(2): 135–143.

Moris, N., C. Pina and A. M. Arias (2016). “Transition states and cell fate decisions in epigenetic landscapes.” Nat Rev Genet 17(11): 693–703.

Nishida, K., M. Hoshino, Y. Kawaguchi and F. Murakami (2010). “Ptf1a directly controls expression of immunoglobulin superfamily molecules Nephrin and Neph3 in the developing central nervous system.” J Biol Chem 285(1): 373–380.

Noctor, S. C., V. Martínez-Cerdeño, L. Ivic and A. R. Kriegstein (2004). “Cortical neurons arise in symmetric and asymmetric division zones and migrate through specific phases.” Nat Neurosci 7(2): 136–144.

Ortega, J. A., N. V. Radonjić and N. Zecevic (2013). “Sonic hedgehog promotes generation and maintenance of human forebrain Olig2 progenitors.” Front Cell Neurosci 7: 254.

Pan, Z. Z. (2012). “Transcriptional control of Gad2.” Transcription 3(2): 68–72.

Perkins, T. J. and P. S. Swain (2009). “Strategies for cellular decision-making.” Mol Syst Biol 5: 326.

Petryniak, M. A., G. B. Potter, D. H. Rowitch and J. L. Rubenstein (2007). “Dlx1 and Dlx2 control neuronal versus oligodendroglial cell fate acquisition in the developing forebrain.” Neuron 55(3): 417–433.

Pinto, L., D. Drechsel, M. T. Schmid, J. Ninkovic, M. Irmler, M. S. Brill, L. Restani, L. Gianfranceschi, C. Cerri, S. N. Weber, V. Tarabykin, K. Baer, F. Guillemot, J. Beckers, N. Zecevic, C. Dehay, M. Caleo, H. Schorle and M. Götz (2009). “AP2gamma regulates basal progenitor fate in a region-and layer-specific manner in the developing cortex.” Nat Neurosci 12(10): 1229–1237.

Pla, R., A. Stanco, M. A. Howard, A. N. Rubin, D. Vogt, N. Mortimer, I. Cobos, G. B. Potter, S. Lindtner, J. D. Price, A. S. Nord, A. Visel, C. E. Schreiner, S. C. Baraban, D. H. Rowitch and J. L. R. Rubenstein (2018). “Dlx1 and Dlx2 Promote Interneuron GABA Synthesis, Synaptogenesis, and Dendritogenesis.” Cereb Cortex 28(11): 3797–3815.

Quinn, J. C., M. Molinek, B. S. Martynoga, P. A. Zaki, A. Faedo, A. Bulfone, R. F. Hevner, J. D. West and D. J. Price (2007). “Pax6 controls cerebral cortical cell number by regulating exit from the cell cycle and specifies cortical cell identity by a cell autonomous mechanism.” Dev Biol 302(1): 50–65.

Quinn, J. C., M. Molinek, J. O. Mason and D. J. Price (2009). “Gli3 is required autonomously for dorsal telencephalic cells to adopt appropriate fates during embryonic forebrain development.” Dev Biol 327(1): 204–215.

Quintana-Urzainqui, I., Z. Kozić, S. Mitra, T. Tian, M. Manuel, J. O. Mason and D. J. Price (2018). “Tissue-Specific Actions of Pax6 on Proliferation and Differentiation Balance in Developing Forebrain Are Foxg1 Dependent.” iScience 10: 171–191.

Radler-Pohl, A., S. Gebel, C. Sachsenmaier, H. König, M. Krämer, T. Oehler, M. Streile, H. Ponta, U. Rapp, H. J. Rahmsdorf and et al. (1993). “The activation and activity control of AP-1 (fos/jun).” Ann N Y Acad Sci 684: 127–148.

Radonjić, N. V., F. Memi, J. A. Ortega, N. Glidden, H. Zhan and N. Zecevic (2016). “The Role of Sonic Hedgehog in the Specification of Human Cortical Progenitors In Vitro.” Cereb Cortex 26(1): 131–143.

Raivich, G. and A. Behrens (2006). “Role of the AP-1 transcription factor c-Jun in developing, adult and injured brain.” Prog Neurobiol 78(6): 347–363.

Rallu, M., R. Machold, N. Gaiano, J. G. Corbin, A. P. McMahon and G. Fishell (2002). “Dorsoventral patterning is established in the telencephalon of mutants lacking both Gli3 and Hedgehog signaling.” Development 129(21): 4963–4974.

Rallu, M., J. G. Corbin and G. Fishell (2002b). “Parsing the prosencephalon.” Nat Rev Neurosci 3(12): 943–951.

Raposo, A., F. F. Vasconcelos, D. Drechsel, C. Marie, C. Johnston, D. Dolle, A. Bithell, S. Gillotin, D. L. C. van den Berg, L. Ettwiller, P. Flicek, G. E. Crawford, C. M. Parras, B. Berninger, N. J. Buckley, F. Guillemot and D. S. Castro (2015). “Ascl1 Coordinately Regulates Gene Expression and the Chromatin Landscape during Neurogenesis.” Cell Rep 10(9): 1544–1556.

Robertshaw, E., K. Matsumoto, A. Lumsden and C. Kiecker (2013). “Irx3 and Pax6 establish differential competence for Shh-mediated induction of GABAergic and glutamatergic neurons of the thalamus.” Proc Natl Acad Sci U S A 110(41): E3919–3926.

Robinson, M. D., D. J. McCarthy and G. K. Smyth (2010). “edgeR: a Bioconductor package for differential expression analysis of digital gene expression data.” Bioinformatics 26(1): 139–140.

Sansom, S. N., D. S. Griffiths, A. Faedo, D. J. Kleinjan, Y. Ruan, J. Smith, V. van Heyningen, J. L. Rubenstein and F. J. Livesey (2009). “The level of the transcription factor Pax6 is essential for controlling the balance between neural stem cell self-renewal and neurogenesis.” PLoS Genet 5(6): e1000511.

Saxena, M., N. Agnihotri and J. Sen (2018). “Perturbation of canonical and non-canonical BMP signaling affects migration, polarity and dendritogenesis of mouse cortical neurons.” Development 145(1).

Scala, F., D. Kobak, M. Bernabucci, Y. Bernaerts, C. R. Cadwell, J. R. Castro, L. Hartmanis, X. Jiang, S. Laturnus, E. Miranda, S. Mulherkar, Z. H. Tan, Z. Yao, H. Zeng, R. Sandberg, P. Berens and A. S. Tolias (2021). “Phenotypic variation of transcriptomic cell types in mouse motor cortex.” Nature 598(7879): 144–150.

Scardigli, R., N. Bäumer, P. Gruss, F. Guillemot and I. Le Roux (2003). “Direct and concentration-dependent regulation of the proneural gene Neurogenin2 by Pax6.” Development 130(14): 3269–3281.

Schuurmans, C., O. Armant, M. Nieto, J. M. Stenman, O. Britz, N. Klenin, C. Brown, L. M. Langevin, J. Seibt, H. Tang, J. M. Cunningham, R. Dyck, C. Walsh, K. Campbell, F. Polleux and F. Guillemot (2004). “Sequential phases of cortical specification involve Neurogenin-dependent and -independent pathways.” Embo j 23(14): 2892–2902.

Seibt, J., O. Armant, A. Le Digarcher, D. Castro, V. Ramesh, L. Journot, F. Guillemot, P. Vanderhaeghen and T. Bouschet (2012). “Expression at the imprinted dlk1-gtl2 locus is regulated by proneural genes in the developing telencephalon.” PLoS One 7(11): e48675.

Sessa, A., E. Ciabatti, D. Drechsel, L. Massimino, G. Colasante, S. Giannelli, T. Satoh, S. Akira, F. Guillemot and V. Broccoli (2017). “The Tbr2 Molecular Network Controls Cortical Neuronal Differentiation Through Complementary Genetic and Epigenetic Pathways.” Cereb Cortex 27(6): 3378–3396.

Sheng, M. and M. E. Greenberg (1990). “The regulation and function of c-fos and other immediate early genes in the nervous system.” Neuron 4(4): 477–485.

Shikata, Y., T. Okada, M. Hashimoto, T. Ellis, D. Matsumaru, T. Shiroishi, M. Ogawa, B. Wainwright and J. Motoyama (2011). “Ptch1-mediated dosage-dependent action of Shh signaling regulates neural progenitor development at late gestational stages.” Dev Biol 349(2): 147–159.

Shimamura, K., D. J. Hartigan, S. Martinez, L. Puelles and J. L. Rubenstein (1995). “Longitudinal organization of the anterior neural plate and neural tube.” Development 121(12): 3923–3933.

Shimamura, K. and J. L. Rubenstein (1997). “Inductive interactions direct early regionalization of the mouse forebrain.” Development 124(14): 2709–2718.

Simpson, T. I., T. Pratt, J. O. Mason and D. J. Price (2009). “Normal ventral telencephalic expression of Pax6 is required for normal development of thalamocortical axons in embryonic mice.” Neural Dev 4: 19.

Simpson, T. I. and D. J. Price (2002). “Pax6; a pleiotropic player in development.” Bioessays 24(11): 1041–1051.

Smart, I. H. and M. Smart (1982). “Growth patterns in the lateral wall of the mouse telencephalon: I. Autoradiographic studies of the histogenesis of the isocortex and adjacent areas.” J Anat 134(Pt 2): 273–298.

Soldado-Magraner, S., F. Brandalise, S. Honnuraiah, M. Pfeiffer, M. Moulinier, U. Gerber and R. Douglas (2020). “Conditioning by subthreshold synaptic input changes the intrinsic firing pattern of CA3 hippocampal neurons.” J Neurophysiol 123(1): 90–106.

Stenman, J., H. Toresson and K. Campbell (2003). “Identification of two distinct progenitor populations in the lateral ganglionic eminence: implications for striatal and olfactory bulb neurogenesis.” J Neurosci 23(1): 167–174.

Stoykova, A. and P. Gruss (1994). “Roles of Pax-genes in developing and adult brain as suggested by expression patterns.” J Neurosci 14(3 Pt 2): 1395–1412.

Street, K., D. Risso, R. B. Fletcher, D. Das, J. Ngai, N. Yosef, E. Purdom and S. Dudoit (2018). “Slingshot: cell lineage and pseudotime inference for single-cell transcriptomics.” BMC Genomics 19(1): 477.

Sultan, K. T., K. N. Brown and S. H. Shi (2013). “Production and organization of neocortical interneurons.” Front Cell Neurosci 7: 221.

Sun, J., S. Rockowitz, Q. Xie, R. Ashery-Padan, D. Zheng and A. Cvekl (2015). “Identification of in vivo DNA-binding mechanisms of Pax6 and reconstruction of Pax6-dependent gene regulatory networks during forebrain and lens development.” Nucleic Acids Res 43(14): 6827–6846.

Svärd, J., K. Heby-Henricson, M. Persson-Lek, B. Rozell, M. Lauth, A. Bergström, J. Ericson, R. Toftgård and S. Teglund (2006). “Genetic elimination of Suppressor of fused reveals an essential repressor function in the mammalian Hedgehog signaling pathway.” Dev Cell 10(2): 187–197.

Toresson, H., S. S. Potter and K. Campbell (2000). “Genetic control of dorsal-ventral identity in the telencephalon: opposing roles for Pax6 and Gsh2.” Development 127(20): 4361–4371.

Tritschler, S., M. Büttner, D. S. Fischer, M. Lange, V. Bergen, H. Lickert and F. J. Theis (2019). “Concepts and limitations for learning developmental trajectories from single cell genomics.” Development 146(12).

Tuoc, T. C., K. Radyushkin, A. B. Tonchev, M. C. Piñon, R. Ashery-Padan, Z. Molnár, M. S. Davidoff and Stoykova (2009). “Selective cortical layering abnormalities and behavioral deficits in cortex-specific Pax6 knock-out mice.” J Neurosci 29(26): 8335–8349.

Turque, N., S. Plaza, F. Radvanyi, C. Carriere and S. Saule (1994). “Pax-QNR/Pax-6, a paired box-and homeobox-containing gene expressed in neurons, is also expressed in pancreatic endocrine cells.” Mol Endocrinol 8(7): 929–938.

Tuvikene, J., P. Pruunsild, E. Orav, E. E. Esvald and T. Timmusk (2016). “AP-1 Transcription Factors Mediate BDNF-Positive Feedback Loop in Cortical Neurons.” J Neurosci 36(4): 1290–1305.

Tyzio, R., A. Ivanov, C. Bernard, G. L. Holmes, Y. Ben-Ari and R. Khazipov (2003). “Membrane potential of CA3 hippocampal pyramidal cells during postnatal development.” J Neurophysiol 90(5): 2964–2972.

Van den Berge, K., H. Roux de Bézieux, K. Street, W. Saelens, R. Cannoodt, Y. Saeys, S. Dudoit and L. Clement (2020). “Trajectory-based differential expression analysis for single-cell sequencing data.” Nat Commun 11(1): 1201.

Vasistha, N. A., F. García-Moreno, S. Arora, A. F. Cheung, S. J. Arnold, E. J. Robertson and Z. Molnár (2015). “Cortical and Clonal Contribution of Tbr2 Expressing Progenitors in the Developing Mouse Brain.” Cereb Cortex 25(10): 3290–3302.

Villalba, A., M. Götz and V. Borrell (2021). “The regulation of cortical neurogenesis.” Curr Top Dev Biol 142: 1–66.

Viti, J., A. Gulacsi and L. Lillien (2003). “Wnt regulation of progenitor maturation in the cortex depends on Shh or fibroblast growth factor 2.” J Neurosci 23(13): 5919–5927.

Voronova, A., S. A. Yuzwa, B. S. Wang, S. Zahr, C. Syal, J. Wang, D. R. Kaplan and F. D. Miller (2017). “Migrating Interneurons Secrete Fractalkine to Promote Oligodendrocyte Formation in the Developing Mammalian Brain.” Neuron 94(3): 500–516.e509.

Waclaw, R. R., B. Wang, Z. Pei, L. A. Ehrman and K. Campbell (2009). “Distinct temporal requirements for the homeobox gene Gsx2 in specifying striatal and olfactory bulb neuronal fates.” Neuron 63(4): 451–465.

Waddington, C.H. (1932). “Experiments on the Development of Chick and Duck Embryos, Cultivated in Vitro.” Philos Trans R Soc Lond B Biol Sci 221: 179–230.

Waddington, C.H. (1934). “Experiments on Embryonic Induction: III. A Note on Inductions by Chick Primitive Streak Transplanted to the Rabbit Embryo.” J Exp Biol 11: 224–227.

Waddington, C.H. (1957). “The Strategy of the Gene.” London: George Allen and Unwin.

Walther, C. and P. Gruss (1991). “Pax-6, a murine paired box gene, is expressed in the developing CNS.” Development 113(4): 1435–1449.

Wang, B., J. E. Long, P. Flandin, R. Pla, R. R. Waclaw, K. Campbell and J. L. Rubenstein (2013). “Loss of Gsx1 and Gsx2 function rescues distinct phenotypes in Dlx1/2 mutants.” J Comp Neurol 521(7): 1561–1584.

Ward, J. H. (1963). “Hierarchical grouping to optimize an objective function.” Journal of the American Statistical Association, 58, 236–244.

Wechsler-Reya, R. J. and M. P. Scott (1999). “Control of neuronal precursor proliferation in the cerebellum by Sonic Hedgehog.” Neuron 22(1): 103–114.

Wei, S., H. Du, Z. Li, G. Tao, Z. Xu, X. Song, Z. Shang, Z. Su, H. Chen, Y. Wen, G. Liu, Y. You, Z. Zhang and Z. Yang (2019). “Transcription factors Sp8 and Sp9 regulate the development of caudal ganglionic eminence-derived cortical interneurons.” J Comp Neurol 527(17): 2860–2874.

Weisenhorn, D. M., E. W. Prieto and M. R. Celio (1994). “Localization of calretinin in cells of layer I (Cajal-Retzius cells) of the developing cortex of the rat.” Brain Res Dev Brain Res 82(1-2): 293–297.

Wijgerde, M., J. A. McMahon, M. Rule and A. P. McMahon (2002). “A direct requirement for Hedgehog signaling for normal specification of all ventral progenitor domains in the presumptive mammalian spinal cord.” Genes Dev 16(22): 2849–2864.

Winkler, C. C., O. R. Yabut, S. P. Fregoso, H. G. Gomez, B. E. Dwyer, S. J. Pleasure and S. J. Franco (2018). “The Dorsal Wave of Neocortical Oligodendrogenesis Begins Embryonically and Requires Multiple Sources of Sonic Hedgehog.” J Neurosci 38(23): 5237–5250.

Wolpert, L. (1969). “Positional information and the spatial pattern of cellular differentiation.” J Theor Biol 25(1): 1–47.

Wong, F. K., J. F. Fei, F. Mora-Bermúdez, E. Taverna, C. Haffner, J. Fu, K. Anastassiadis, A. F. Stewart and W. B. Huttner (2015). “Sustained Pax6 Expression Generates Primate-like Basal Radial Glia in Developing Mouse Neocortex.” PLoS Biol 13(8): e1002217.

Xie, Q., Y. Yang, J. Huang, J. Ninkovic, T. Walcher, L. Wolf, A. Vitenzon, D. Zheng, M. Götz, D. C. Beebe, J. Zavadil and A. Cvekl (2013). “Pax6 interactions with chromatin and identification of its novel direct target genes in lens and forebrain.” PLoS One 8(1): e54507.

Xu, Q., L. Guo, H. Moore, R. R. Waclaw, K. Campbell and S. A. Anderson (2010). “Sonic hedgehog signaling confers ventral telencephalic progenitors with distinct cortical interneuron fates.” Neuron 65(3): 328–340.

Yabut, O. R., G. Fernandez, T. Huynh, K. Yoon and S. J. Pleasure (2015). “Suppressor of Fused Is Critical for Maintenance of Neuronal Progenitor Identity during Corticogenesis.” Cell Rep 12(12): 2021–2034.

Yabut, O. R., H. X. Ng, K. Yoon, J. C. Arela, T. Ngo and S. J. Pleasure (2020). “The Neocortical Progenitor Specification Program Is Established through Combined Modulation of SHH and FGF Signaling.” J Neurosci 40(36): 6872–6887.

Yabut, O. R. and S. J. Pleasure (2018). “Sonic Hedgehog Signaling Rises to the Surface: Emerging Roles in Neocortical Development.” Brain Plast 3(2): 119–128.

Yang, J. W., W. Kilb, S. Kirischuk, P. Unichenko, M. C. Stüttgen and H. J. Luhmann (2018). “Development of the whisker-to-barrel cortex system.” Curr Opin Neurobiol 53: 29–34.

Yao, J., Y. Liu, J. Husain, R. Lo, A. Palaparti, J. Henderson and S. Stifani (1998). “Combinatorial expression patterns of individual TLE proteins during cell determination and differentiation suggest non-redundant functions for mammalian homologs of Drosophila Groucho.” Dev Growth Differ 40(2): 133–146.

Young, M. D. and S. Behjati (2020). “SoupX removes ambient RNA contamination from droplet-based single-cell RNA sequencing data.” Gigascience 9(12).

Yun, K., S. Fischman, J. Johnson, M. Hrabe de Angelis, G. Weinmaster and J. L. Rubenstein (2002). “Modulation of the notch signaling by Mash1 and Dlx1/2 regulates sequential specification and differentiation of progenitor cell types in the subcortical telencephalon.” Development 129(21): 5029–5040.

Yun, K., S. Potter and J. L. Rubenstein (2001). “Gsh2 and Pax6 play complementary roles in dorsoventral patterning of the mammalian telencephalon.” Development 128(2): 193–205.

Zhang, Y., G. Liu, T. Guo, X. G. Liang, H. Du, L. Yang, A. Bhaduri, X. Li, Z. Xu, Z. Zhang, Z. Li, M. He, J. Tsyporin, A. R. Kriegstein, J. L. Rubenstein, Z. Yang and B. Chen (2020). “Cortical Neural Stem Cell Lineage Progression Is Regulated by Extrinsic Signaling Molecule Sonic Hedgehog.” Cell Rep 30(13): 4490–4504.e4494.

Zimmer, C., M. C. Tiveron, R. Bodmer and H. Cremer (2004). “Dynamics of Cux2 expression suggests that an early pool of SVZ precursors is fated to become upper cortical layer neurons.” Cereb Cortex 14(12): 1408–1420.

